# Old fibroblasts secrete inflammatory cytokines that drive variability in reprogramming efficiency and may affect wound healing between old individuals

**DOI:** 10.1101/448431

**Authors:** Salah Mahmoudi, Elena Mancini, Alessandra Moore, Lucy Xu, Fereshteh Jahanbani, Katja Hebestreit, Rajini Srinivasan, Xiyan Li, Keerthana Devarajan, Laurie Prélot, Cheen Euong Ang, Yohei Shibuya, Bérénice A. Benayoun, Anne Lynn S. Chang, Marius Wernig, Joanna Wysocka, Michael T. Longaker, Michael P. Snyder, Anne Brunet

**Author notes:** Correspondence: Anne Brunet. These authors contributed equally to this work (Salah Mahmoudi, Elena Mancini).

## Abstract

Age-associated chronic inflammation (inflammaging) has emerged as a central hallmark of aging^1-3^, but its impact on specific cells is still largely unknown. Fibroblasts are present in all tissues and contribute to wound healing^4-6^. They are also the cell type that is mostly used for induced pluripotent stem cell (iPSC) reprogramming^7^ – a process that has implications for regenerative medicine and rejuvenation strategies^8-17^. Here we show that primary fibroblasts from old mice secrete inflammatory cytokines and that there is an increased variability in reprogramming efficiency between fibroblast cultures from old individuals. Individual-to-individual variability is emerging as a key feature of old age^18-21^, which could reflect distinct aging trajectories, but the underlying causes remain unknown. To identify drivers of this variability, we perform a multi-omic assessment of young and old fibroblast cultures with different reprogramming efficiency. This approach, coupled with single cell transcriptomics, reveals that old fibroblast cultures are heterogeneous and show a greater proportion of ‘activated fibroblasts’ that secrete inflammatory cytokines, which correlates with reprogramming efficiency. We experimentally validate that activated fibroblasts express inflammatory cytokines *in vivo* and that their presence is linked to enhanced reprogramming efficiency in culture. Conditioned-media swapping experiments show that extrinsic factors secreted by activated fibroblasts are more critical than intrinsic factors for the individual-to-individual variability in reprogramming efficiency, and we identify TNFα as a key inflammatory cytokine underlying this variability. Interestingly, old mice also exhibit variability in wound healing efficiency *in vivo* and old wounds show an increased subpopulation of activated fibroblasts with a unique TNFα signature. Our study shows that a switch in fibroblast composition, and the ratio of inflammatory cytokines they secrete, drives variability in reprogramming *in vitro* and may influence wound healing *in vivo*. These findings could help identify personalized strategies to improve iPSC generation and wound healing in older individuals.

## Main

A key feature of old age is a chronic inflammatory state with elevated levels of inflammatory cytokines^1-3^, also known as ‘inflammaging’. Inflammaging is linked to many age-related diseases and conditions, including cardiovascular diseases, cancer, and fibrosis^2,22,23^. Thus, understanding how inflammaging influences cells and tissues is critical for identifying new strategies to counteract aging and age-related diseases. Fibroblasts are present in all tissues and are known to respond to systemic and local inflammatory signals, notably during wounding^4-6^. Accordingly, fibroblasts play an important role in wound healing as well as in pathological processes following injury such as fibrosis^4-6^. Senescent fibroblasts have been shown to secrete cytokines that negatively impact tissue function during aging^22,24-28^. However, how normal fibroblasts respond to the chronic inflammation that accompanies aging, and how this impacts their function, remains unknown.

Fibroblasts are also the most frequently used cell type for iPSC reprogramming in mice and humans^7^. Reprogramming to an induced pluripotent stem cell (iPSC) state has important implications for regeneration and ‘rejuvenation’^8-17^. Because iPSCs can generate many different cell types, reprogramming of cells from patients could facilitate replacement of cells in deteriorated tissues in aging and age-related diseases^29,30^. Reprogramming might also be useful as a ‘rejuvenation strategy’, as it can erase hallmarks of aging *in vitro*^8-14^ and cycles of reprogramming can ameliorate tissue function in old mice^15^. Several studies have probed the impact of aging on reprogramming, but the results are not yet clear, with some studies reporting a decline in reprogramming efficiency with age^31-36^ and others reporting no change^13,37^ or an increase^38-40^. In addition, senescent fibroblasts have been reported to have both positive and negative effects on reprogramming^33,39,41,42^. Hence, a systematic evaluation of how aging influences reprogramming, and how inflammaging affects this process, is lacking.

We examined the impact of old age on the inflammatory profile of fibroblasts and their ability to reprogram (Fig. 1a). Using cytokine profiling, we compared the systemic milieu (plasma), conditioned media from primary fibroblast cultures, and conditioned media from resulting iPSCs from young (3 months) and old (28-29 months) mice (Fig. 1a). Plasma from old mice showed increased levels of pro-inflammatory cytokines (e.g. IL6 and TNFα), anti-inflammatory cytokines (e.g. IL4 and IL5), and chemokines and growth factors (e.g. MIP2 and MCSF) compared to plasma from young mice (Fig. 1b, Extended Data Fig. 1a, Supplementary Table 1a). Conditioned media from low-passage (P3) primary fibroblast cultures from the ears of old mice also showed enhanced levels of both pro-and anti-inflammatory cytokines (e.g. IL6 and TNFα, and IL4, respectively) (Fig. 1c, Extended Data Fig. 1b, c, Supplementary Table 1b). This increase in inflammatory cytokines was also observed in conditioned media from other types of fibroblasts (lung) in old mice (Extended Data Fig. 1d, Supplementary Table 1c), and from human primary fibroblasts (Fig. 1g, Extended Data Fig. 1e, Supplementary Table 1d). Thus, primary cultures of fibroblasts from old individuals exhibit a secretory inflammatory profile that overlaps in part with that of the systemic milieu (Fig. 1g, Extended Data Fig. 1c, see Fig. 3 and 4 for *in vivo* data). In contrast, conditioned media from iPSC lines derived from old fibroblast cultures (see Experimental Procedures) no longer showed increased inflammatory cytokines compared to conditioned media of iPSCs derived from young cultures (Fig. 1f, g, Supplementary Table 1e). Therefore, old fibroblasts exhibit an inflammatory profile in culture that can be erased by reprogramming.

**Figure 1.**
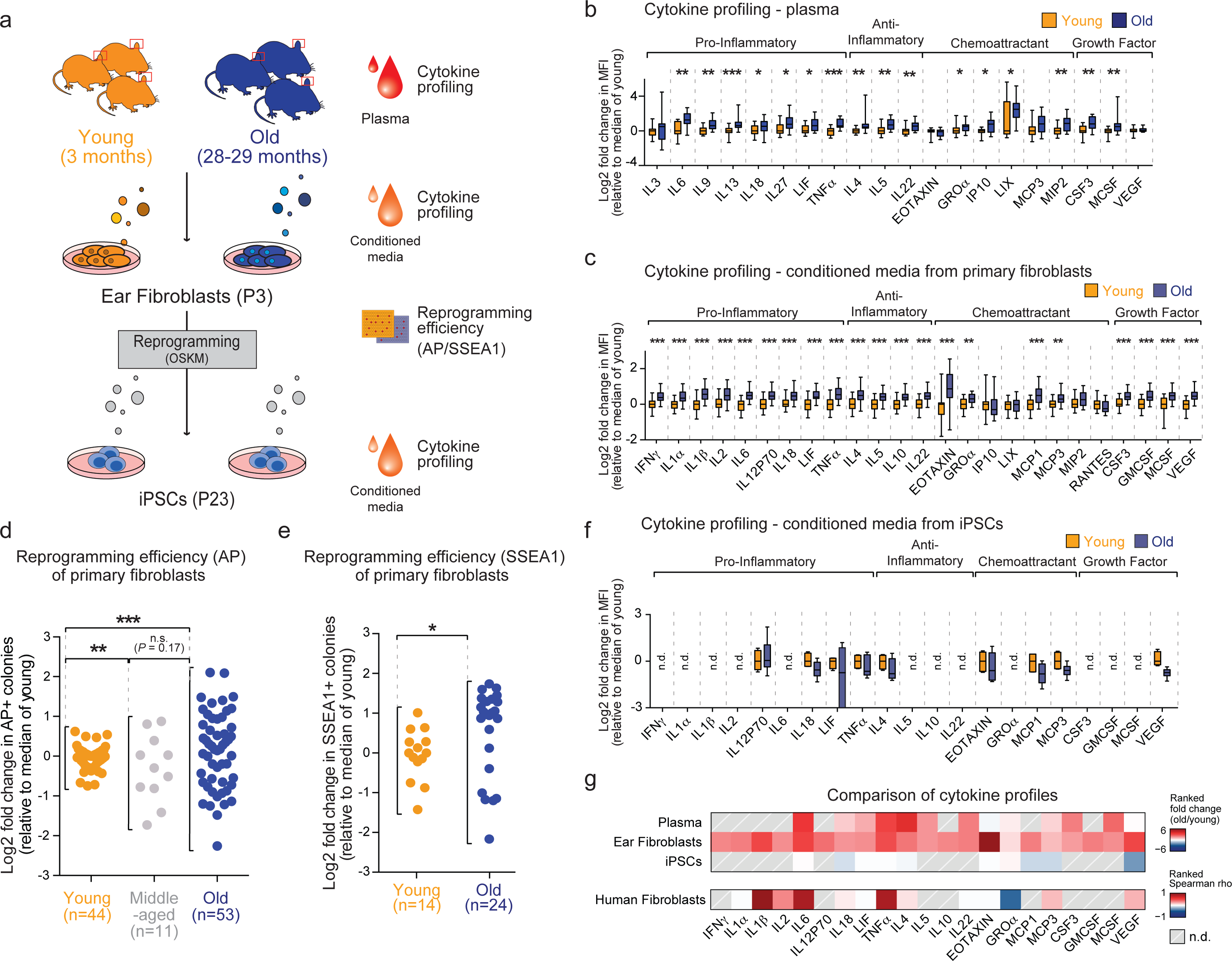
Primary fibroblasts from old mice secrete inflammatory cytokines and exhibit increased individual-to-individual variability in reprogramming efficiency. **a**, Schematic illustration of inflammatory cytokine profiling of the systemic milieu (plasma) and conditioned media from primary cultures of fibroblasts and their resulting induced pluripotent stem cells (iPSCs), and assessment of reprogramming efficiency. Blood was collected from young (3 months) and old (28-29 months) male C57BL/6 mice by cardiac puncture, and plasma (without blood cells) was analysed for cytokine profiling. In parallel, primary fibroblasts were derived from the ears of the mice, and reprogrammed at passage 3 (P3) using Oct3/4, Sox2, Klf4 and c-Myc (OSKM). Conditioned media was collected from the fibroblasts at P3 and the resulting iPSCs at P23 for cytokine profiling. Reprogramming efficiency was determined by staining for alkaline phosphatase (AP) and/or stage-specific embryonic antigen 1 (SSEA1). **b**, Inflammatory cytokine profiles of plasma from young (3 months) and old (28-29 months) mice measured by Luminex multiplex cytokine assay. Results are shown as log2 fold change in mean fluorescence intensity (MFI) over median of young plasma, using a box-and-whisker plot to indicate the median and interquartile range, with whiskers indicating minimum and maximum values. Data are from a total of 21 young and 19 old mice combined from 2 independent experiments. *P*-values were calculated using a two-tailed Wilcoxon rank sum test and were adjusted for multiple hypothesis testing using Benjamini-Hochberg correction. * *P* < 0.05, ** *P* < 0.01, *** *P* < 0.001. **c**, Inflammatory cytokine profiles of conditioned media collected from primary cultures of ear fibroblasts at passage 3 from young (3 months) and old (28-29 months) mice. Results are shown as log2 fold change in MFI over median of young fibroblasts and represented as in b. Data are from a total of 24 young and 24 old mice combined from 3 independent experiments (4 cohorts of mice). For individual experiments, see Supplementary Table 7. *P*-values were calculated using a two-tailed Wilcoxon rank sum test and were adjusted for multiple hypothesis testing using Benjamini-Hochberg correction. ** *P* < 0.01, *** *P* < 0.001. **d**, Reprogramming efficiency (RE), assessed by alkaline phosphatase (AP) staining, of young (3 months), middle-aged (12 months) and old (28-29 months) ear fibroblast cultures at P3. Results are shown as log2 fold change in RE over the median RE of young fibroblasts. Data shown are from a total of 44 young, 11 middle-aged, and 53 old mice, combined from 7 independent experiments (for individual experiments, see Supplementary Table 7). Each dot represents cells from one mouse. Statistical differences in variance between age groups were calculated using the non-parametric Fligner-Killeen test, and *P* values are adjusted for multiple hypotheses testing using Benjamini-Hochberg correction. ** *P* < 0.01, *** *P* < 0.001. n.s. = not significant. Variability was not due to combining different cohorts (see Experimental Procedures for statistical testing). This lack of difference was also unlikely to be due to extensive passage of the cells, as old ear fibroblast cultures at passage 33 still exhibited increased secretory inflammatory profile (see Supplementary Fig. 2c, Supplementary Table 1f) and inflammatory signature (see Supplementary Fig. 4j-l). **e**, Reprogramming efficiency (RE), assessed by SSEA1 staining, of young and old ear fibroblast cultures at P3. Results are shown as log2 fold change in RE over the median RE of young fibroblasts. Data shown are from a total of 14 young and 24 old mice, combined from 3 independent experiments (for individual experiments, see Supplementary Table 7). Each dot represents cells from an individual mouse. *P* value was calculated using the non-parametric Fligner-Killeen test. * *P* < 0.05. **f**, Inflammatory cytokine profiles of conditioned media collected from passage 23 cultures of iPSC lines derived from young and old fibroblasts. Results are shown as log2 fold change in MFI over median of young iPSC from one experiment and represented as in b. Data are from 4 young and 6 old iPSC lines derived from one experiment (see Supplementary Table 1f for more details). Only cytokines that were detected at significantly different levels in young and old fibroblasts are shown (for a complete cytokine list, see Supplementary Table 1e). *P*-values were calculated using a one-tailed Wilcoxon rank sum test and were adjusted for multiple hypothesis testing using Benjamini-Hochberg correction. n.d. = not detected. **g**, Comparison of age-dependent changes in cytokine levels between plasma and conditioned media from mouse fibroblasts, their derived iPSCs, and from human fibroblasts. Based on cytokines that are significantly different in conditioned media from fibroblasts (Fig. 1c). Upper panel: Ranked fold change (old/young) in levels of the indicated cytokines in plasma, conditioned media from mouse primary fibroblasts and iPSCs. Lower panel: Ranked Spearman rho (i) correlations for the indicated cytokines in conditioned media from human primary fibroblasts (see Extended Data Fig. 1e for individual ρ). n.d. = not detected.

**Extended Data Figure 1.**
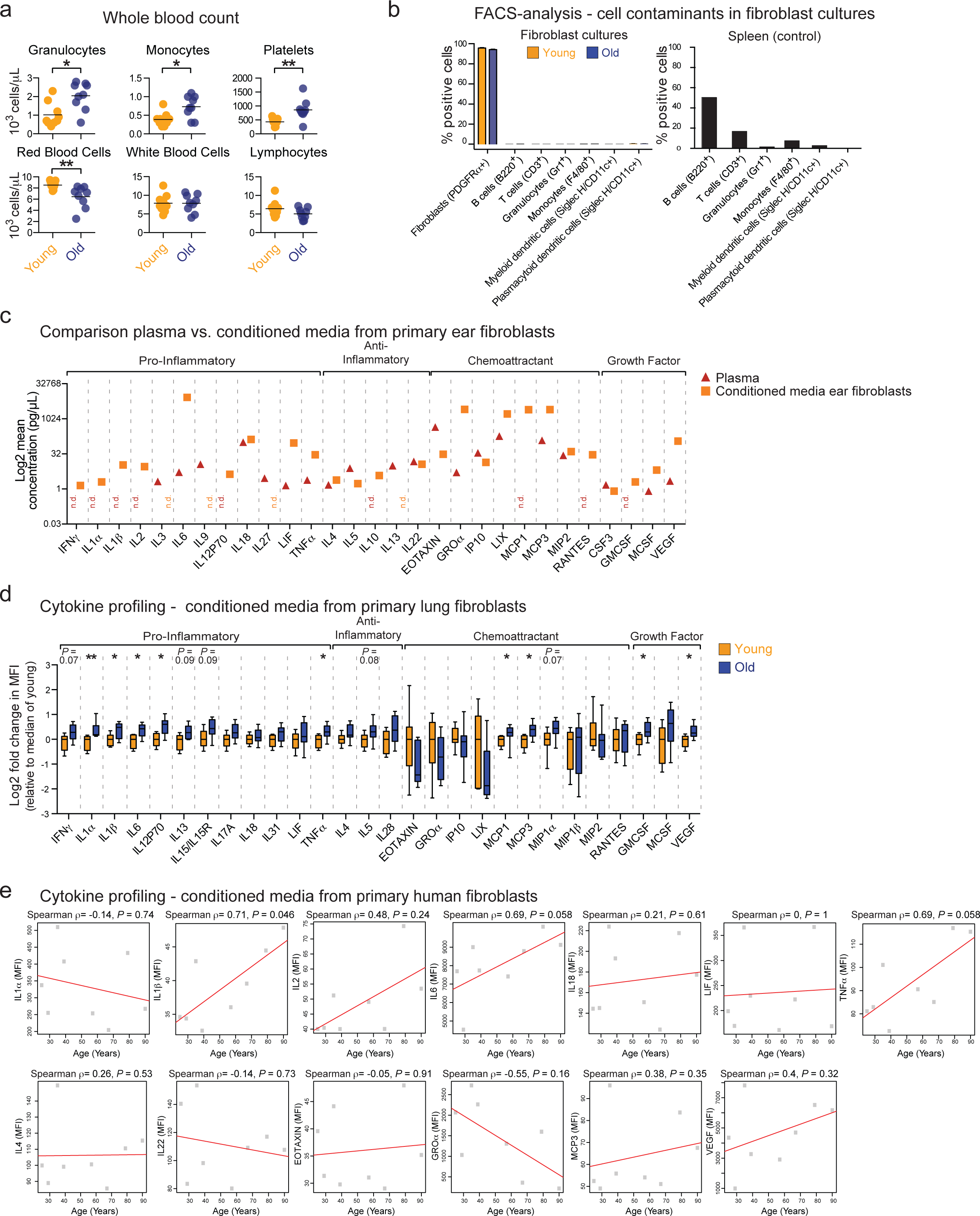
Primary old fibroblasts from mouse ear, mouse lungs, and human skin secrete high levels of inflammatory cytokines. **a**, To gain a better understanding of the systemic changes that occur with age, we analyzed blood cell composition in young (3 months) and old (28-29 months) male C57BL/6 mice using Hemavet Multispecies Hematology Analyzer. Results are shown as cell numbers of the indicated cell types per µL in whole blood samples. Data are from a total of 9 young and 9 old mice combined from 2 independent experiments. Each dot represents cells from one mouse. Lines depict median. *P*-values were calculated using a two-tailed Wilcoxon rank sum test and were adjusted for multiple hypothesis testing using Benjamini-Hochberg correction. * *P*< 0.05, ** *P* < 0.01. **b**, Passage 3 fibroblast cultures contained a high percentage of fibroblast cells (PDGFRα+) and virtually no detectable immune cells, as determined by flow cytometry using the indicated cell-specific surface markers on young and old fibroblast cultures (left panel). Data are from a total of 8 young and 8 old mice from one experiment. Bars represent the mean −/+ SEM. Primary splenocytes from an 8-week-old mouse were used as positive controls for the indicated immune cell-specific surface markers (right panel). **c**, Comparison between inflammatory cytokine profiling of plasma (red triangles) and of conditioned media from cultured ear fibroblasts at passage 3 (orange squares). Results are shown as the mean log2 concentrations (pg/µL) of cytokines detected. For exact concentrations, see Supplementary Tables 1a-b. n.d. = not detected. **d**, Primary fibroblast cultures from the lungs of young (3 months) and old (20-24 months) mice exhibit an increased secretion of inflammatory cytokines with age. Inflammatory cytokine profiles of conditioned media collected from cultures of young and old fibroblasts at passage 3. Results are shown as log2 fold change in mean fluorescence intensity (MFI) over median of young fibroblasts, using a box-and-whisker plot to indicate the median and interquartile range with whiskers indicating minimum and maximum values. Data are from a total of 8 young and 9 old fibroblast cultures combined from 2 independent experiments (2 cohorts of mice). For individual experiments, see Supplementary Table 7. *P-* values were calculated using a two-tailed Wilcoxon rank sum test and were adjusted for multiple hypothesis testing using Benjamini-Hochberg correction. * *P* < 0.05, ** *P* < 0.01. **e**, Primary fibroblasts cultures isolated from punch biopsy of pre-auricular skin of healthy human subjects exhibit increased secretion of several inflammatory cytokines with age. Results are shown as Spearman rank correlation coefficient (ρ) between donor age (years, x-axis) and the levels of individual cytokines (mean fluorescence intensity (MFI), y-axis) in human fibroblast cultures at passage 3. Data are from a total of 8 healthy human samples from 1 experiment. *P*-values were computed using algorithm AS 89 and were adjusted for multiple hypothesis testing using Benjamini-Hochberg correction. For multiple hypothesis testing see Supplementary Table 7.

We next determined how old age impacts the efficiency of reprogramming – a pivotal parameter in the generation of iPSCs from aged patients. To systematically test the effect of age on iPSC reprogramming, we used large cohorts of mice. We derived independent fibroblast cultures from a total of 108 mice from three age groups: young (3 months), middle-aged (12 months), and old (28-29 months) in 7 independent cohorts (Fig. 1a). We induced reprogramming by expressing *OCT4*, *KLF4*, *SOX2*, and *c-MYC* in a single lentiviral construct. For each independent fibroblast culture, reprogramming efficiency was determined by counting, in 96-well plates seeded at low density, the number of forming colonies that are positive for alkaline phosphatase (AP), an early marker of pluripotency^43^, and the stage-specific embryonic antigen 1 (SSEA1), a later marker of pluripotency^43^. We did not observe a significant change in the mean reprogramming efficiency of cells from older individuals (Fig. 1d-e). In contrast, there was a significant increase in the individual-to-individual variability of reprogramming efficiency in cells from old donors, with cells from some old individuals reprogramming either better or worse than those from young individuals (Fig. 1d-e). A similar age-dependent increase in variability in reprogramming efficiency was also observed in cultures of fibroblasts derived from the chest (Extended Data Fig. 2d). This variability appeared to be linked to each fibroblast culture (derived from an individual mouse), as reprogramming efficiency of the same culture was largely consistent in independent reprogramming experiments (Extended Data Fig. 2e) and even in direct reprogramming to induced neurons (Extended Data Fig. 2f). These results indicate an increased variability in reprogramming efficiency between fibroblast cultures from old individuals.

**Extended Data Figure 2.**
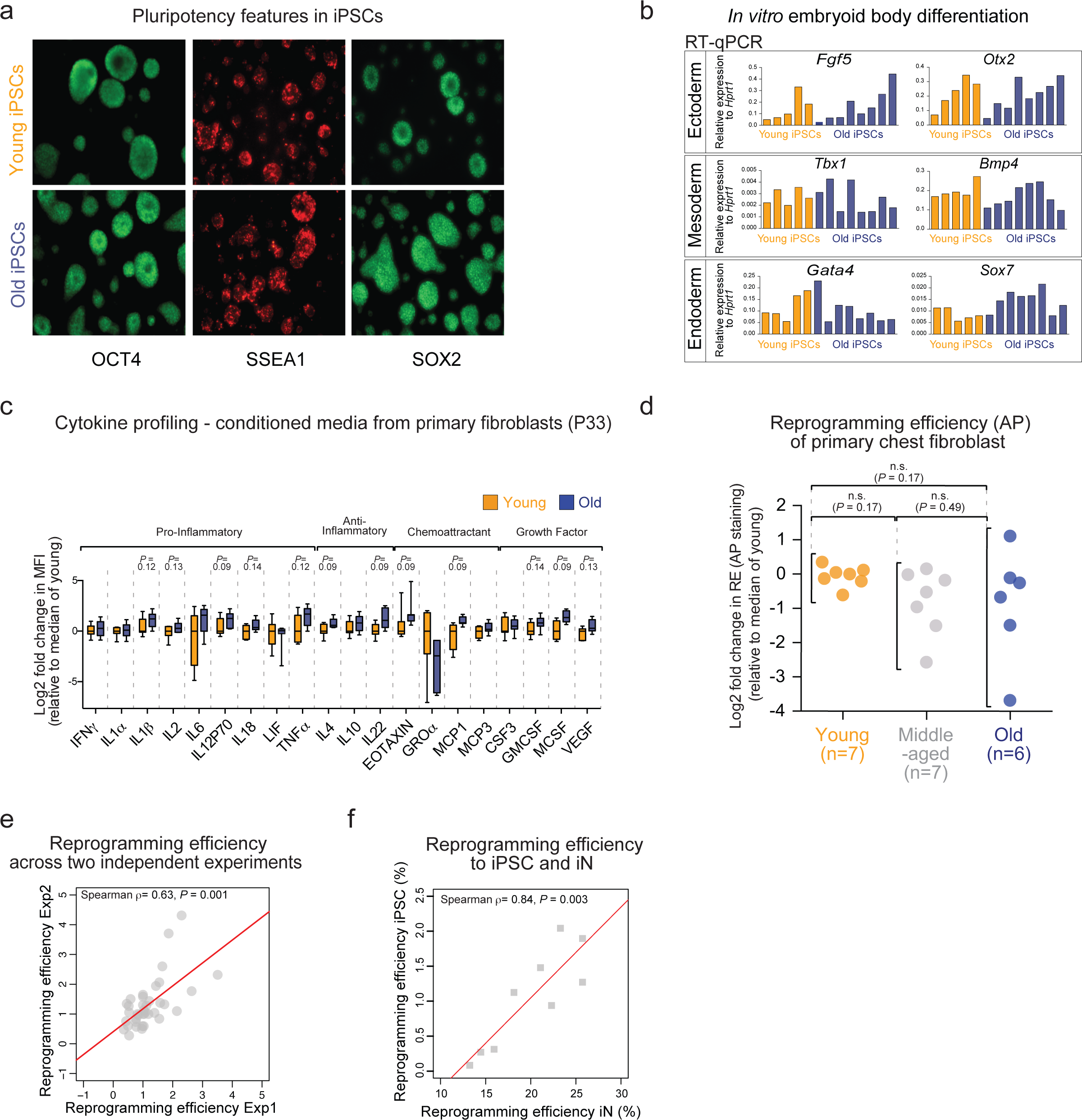
iPSC lines derived from young and old fibroblasts express pluripotency markers and give rise to cell types from all three germ layers, and the ability of individual fibroblast cultures to reprogram is stereotypical. **a-b**, Young and old derived iPSC lines show typical morphologies of mouse iPSCs, express comparable levels of pluripotency markers, and can give rise to cell types from all three germ layers upon embryoid body formation. **a**, Representative immunofluorescence (IF) images of young and old derived iPSC lines at passage 23, stained with the indicated antibodies. **b**, Real-time quantitative PCR (RT-qPCR) on the indicated genes in embryoid bodies *in vitro* differentiated from 5 young and 8 old iPSC lines at passage 23. Expression is presented as expression relative to the housekeeping gene *Hprt1*. Each bar represents expression of one iPSC line. Data are from a total of 13 iPSC lines from 1 experiment. **c**, To test if the extensive *in vitro* culturing, rather than the reprogramming process *per se*, erases cytokine signatures, young and old ear fibroblasts were passaged to passage 33 (P33) and their inflammatory cytokine profile was analyzed. Inflammatory cytokine profiles of conditioned media collected from cultures of young and old ear fibroblasts at passage 33. Results are shown as log2 fold change in mean fluorescence intensity (MFI) over median of young fibroblasts at P33, using a box-and-whisker plot to indicate the median and interquartile range with whiskers indicating minimum and maximum values. Data are from a total of 7 young and 7 old fibroblast cultures from 1 independent experiment. Only the cytokines that exhibited significant difference in levels in conditioned media from young and old at fibroblast passage 3 are shown. For a complete cytokine list, see Extended Data Table 1f. *P-* values were calculated using a one-tailed Wilcoxon rank sum test and adjusted for multiple hypothesis testing using Benjamini-Hochberg correction. Note that the experiments in fibroblasts at passage 3 and 33 were conducted independently, and therefore statistical comparisons were restricted to within experiments. However, a direct comparison between the levels of secreted factors at passage 3 to 33 found that the levels of a majority of cytokines decrease upon passaging. **d**, Reprogramming efficiency (RE), assessed by alkaline phosphatase (AP) staining, of young (3 months), middle-aged (12-13 months) and old (28-30 months) chest fibroblast cultures at P3. Results are shown as log2 fold change in RE over median RE of young fibroblasts. Data shown are from a total of 7 young, 7 middle-aged, and 6 old mice, combined from 2 independent experiments. Each dot represents cells from one mouse. Statistical differences in variance between the age groups calculated using the non-parametric Fligner-Killeen test and adjusted for multiple hypotheses testing using Benjamini-Hochberg correction were not significant (n.s.). **e**, RE of fibroblast cultures are mainly stereotypical to fibroblast cultures from an individual mouse. Correlation plot depicts the RE (assessed by alkaline phosphatase staining) of fibroblast cultures reprogrammed in two independent experiments (separated by > 1 month), with data from experiment 1 on the x-axis, and data from experiment 2 on the y-axis. Data shown are from a total of 14 young, 6 middle-aged, and 18 old mice, combined from 3 independent experiments (for individual experiments, see Supplementary Table 7). Each dot represents cells from an individual mouse. There was a positive correlation (Spearman rank correlation, ρ = 0.63, *P*-value < 0.001) between RE of fibroblast cultures from the same individual. **f**, Correlation between the ability of individual young and old fibroblast cultures at passage 3 to reprogram into neurons (RE iN, x-axis) or to iPSCs (RE iPSC, y-axis). iN and iPSC RE were assessed by PSA-NCAM and alkaline phosphatase staining, respectively. Data shown are from a total of 4 young and 5 old fibroblast cultures from 1 independent experiment. Each dot represents cells from an individual mouse. There was a significant positive correlation (Spearman rank correlation, ρ = 0.84, *P*-value = 0.003) between these two features.

Variability between old individuals has been observed for several biological features, including blood metabolites^18^, methylation status^19^, and physical and cognitive functions^20,21^, and it could reflect distinct aging trajectories. Within an individual, variability between old cells has also been reported for the transcriptome of cardiomyocytes, immune cells, pancreatic cells, and senescent cells^44-50^. However, the mechanism underlying variability between old individuals, and whether it is linked to variability between old cells, is unknown. To understand in an unbiased manner the variability in reprogramming efficiency of old fibroblast cultures, we used a ‘multiomics’ approach and profiled the transcriptome, chromatin landscape of two histone marks (H3K4me3 and H3K27me3), and metabolome of young and old fibroblasts that reprogrammed well or poorly (hereinafter referred to as ‘good’ or ‘bad’ old, respectively) (Fig. 2a, Supplementary Table 2a). Principal component analysis (PCA) and unsupervised hierarchical clustering showed clear separation between the transcriptomes, epigenomes, and metabolomes of young and old fibroblasts (Fig. 2b, Extended Data Fig. 3a-h). PCA analysis also showed some separation between the transcriptomes and metabolomes of good and bad old fibroblast cultures (Fig. 2c, Extended Data Fig. 3i-j).

**Figure 2.**
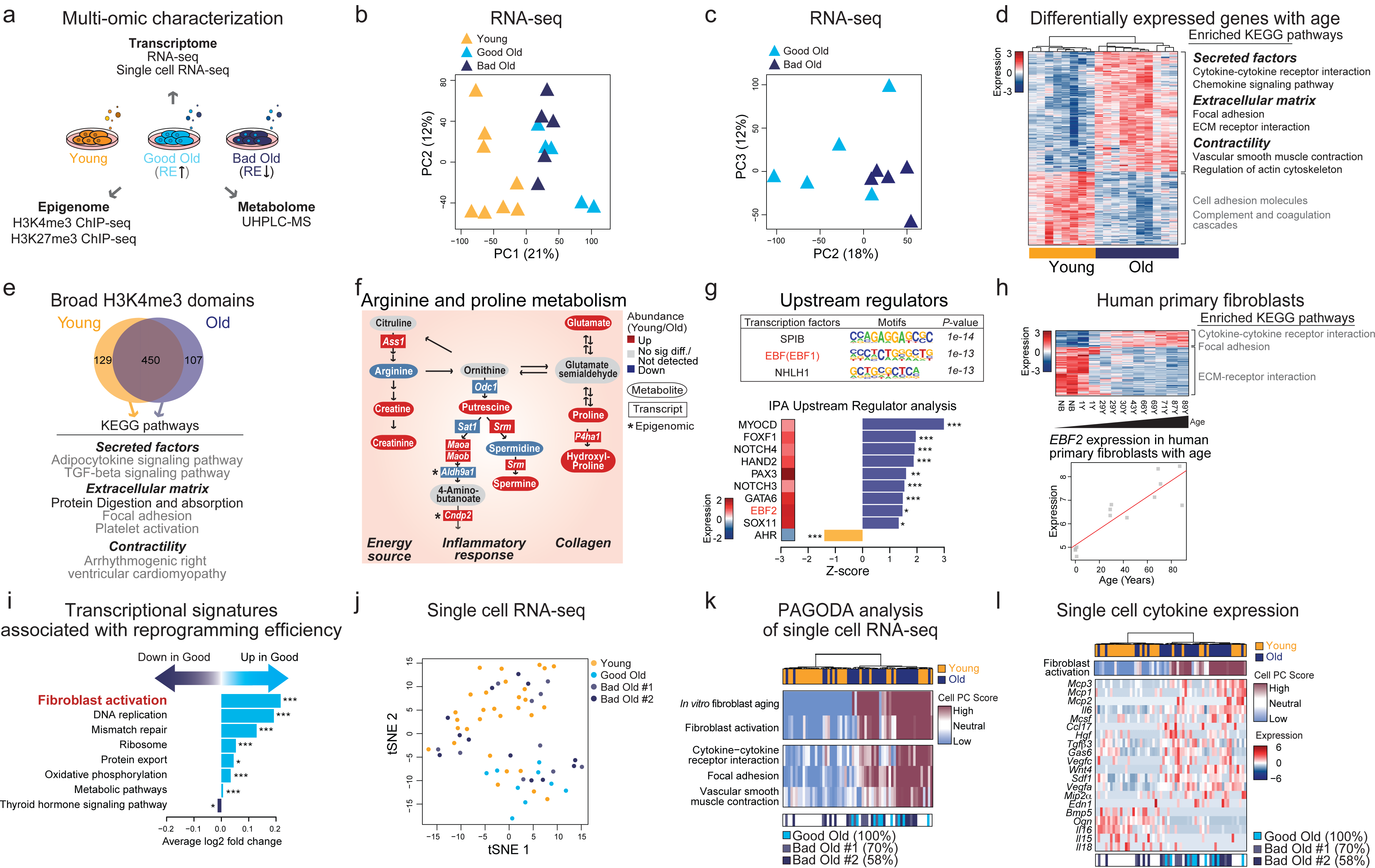
Old fibroblast cultures exhibit a signature of an inflammatory activated state, which is associated with variability in reprogramming efficiency. **a**, Schematic illustration of the transcriptomic (population and single cell RNA-seq), epigenomic, and metabolomic profiling of young (3 months) and old (28-29 months) (‘good’ and ‘bad’ for reprogramming efficiency) fibroblasts at passage 3. **b**, Principal component analysis (PCA) of whole transcriptomes from RNA-seq data of young and old ear fibroblast cultures. PC1 and 2 are shown. Based on reprogramming efficiency (RE), old cultures are depicted either as ‘good’ (high RE) or ‘bad’ (low RE) (see Supplementary Table 2a). Each triangle represents a transcriptome from an individual mouse. Data shown are from a total of 8 young and 10 old mice combined from 3 independent experiments. **c**, PCA of whole transcriptomes from RNA-seq data of only good and bad old ear fibroblast cultures. PC 2 and 3 are shown. PC 1 and 2 are depicted in Extended Data Fig. 3i. Each triangle represents a transcriptome from an individual mouse. **d**, Heatmap of significantly differentially expressed genes between young and old fibroblasts (FDR-values < 0.05, absolute fold change > 1.5) that are part of the indicated KEGG pathways. Expression is depicted as log2 VST (variance stabilizing transformation, DESeq2) normalized read counts, and scaled row-wise. The depicted KEGG pathways are color-coded for significance (black = FDR < 0.05, grey = FDR < 0.15), and further grouped into three classes: Secreted factors, Extracellular matrix and Contractility. For a complete list of significant KEGG terms, see Supplementary Table 2c-e. **e**, Venn diagram depicting the overlap of broad H3K4me3 domains within promoters regions of young and old fibroblasts. In this analysis, the broad H3K4me3 domains that are consistently present in young samples were compared to those that are consistently present in old samples (see Experimental Procedures). The bottom panel depicts a subset of KEGG pathways that are enriched by the genes associated with the broad H3K4me3 domains that change with age. The depicted KEGG pathways are color-coded for significance (black = FDR < 0.05, grey = FDR < 0.15), and are further grouped into two classes: Extracellular matrix and Contractility. For a complete list of significant KEGG terms, see Supplementary Table 2j. **f**, Schematic representation of the biological functions of key metabolites and genes in the arginine and proline metabolic pathway. Abundance of putative metabolites and gene transcripts in old fibroblasts is color-coded (red = higher in old, blue = lower in old, grey = not significantly different/not detected). Metabolites and gene transcripts are depicted in ovals and boxes, respectively. Epigenomic changes in the associated genes are indicated with black stars. **g**, Upper panel: Top 3 motifs found in the promoters of differentially expressed genes between young and old fibroblasts, using HOMER motif analysis. Lower panel: Top 10 putative upstream regulators identified by the Ingenuity Pathway Analysis (IPA) database that are also differentially expressed between young and old fibroblasts (FDR < 0.05, absolute fold change > 1.5). Heatmap depicts log2 fold change in expression (old/young) calculated using DESeq2. The transcription factor identified across both analyses (EBF2) is marked in red. For a complete list of significant motifs and upstream regulators, see Supplementary Table 2p. * *P* < 0.05, ** *P* < 0.01, *** *P* < 0.001. **h**, Upper panel: Heatmap of differentially expressed genes in a regression analysis from young to old healthy human primary fibroblasts^10^ (FDR < 0.05) (Supplementary Table 2q). Expression is depicted as log2 VST (variance stabilizing transformation, DESeq2) normalized read counts, and scaled row-wise. The depicted KEGG pathways are color-coded for significance (black = FDR < 0.05, grey = FDR < 0.15) (Supplementary Table 2r). Lower panel: Expression of *EBF2* across the human samples. The y-axis denotes the log2 VST expression, and the x-axis denotes the age in years. Each square represents a transcriptome from an individual human patient. NB = newborn. **i**, Pathway enrichment analysis of KEGG pathways that are associated with increased (good) or decreased (bad) reprogramming efficiency. The graph shows the pathways that were identified in a regression analysis from bad to good reprogramming efficiency, and that were also identified in a separate analysis comparing the top five to the bottom five cultures in terms of reprogramming efficiency. All depicted KEGG pathways were significantly enriched (FDR < 0.05). For a complete list of pathways, see Supplementary Table 3b, d. * *P* < 0.05, *** *P* < 0.001. **j**, Single cell RNA-seq of young and good and bad old ear fibroblast cultures. tSNE was performed on the log2 VST (variance stabilizing transformation, DESeq2) normalized read counts of all detected genes. Each dot represents a single fibroblast transcriptome. **k**, Pathway and gene set overdispersion analysis (PAGODA) of single cell RNA-seq data from young and good and bad old fibroblasts performed using all KEGG pathways, the “*In vitro* fibroblast aging” and “Fibroblast activation” signature (see Supplementary Table 2b, f), and de novo gene sets after accounting for cell cycle phases (see Extended Data Fig. 5b for data without accounting for cell cycle phases). Hierarchical clustering is based on 84 significantly overdispersed gene sets (see Extended Data Fig. 5c for full list of significant KEGG pathways). Upper heatmap depicts single cells from young and old fibroblast cultures. Middle heatmap shows separation of cells based on their PC scores for a subset of the top significantly overdispersed gene sets. The PC scores are oriented so that high (maroon) and low (blue) values generally correspond to increased and decreased expression, respectively, of the associated gene sets. Lower heatmap depicts single cells from good and bad old fibroblast cultures. **l**, Upper heatmap shows PAGODA of single cell RNA-seq data as described in k. Middle heatmap shows the expression of a subset of the cytokine genes. Expression is depicted as VST-normalized read counts, and scaled row-wise. See Extended Data Fig. 5g for the full list of genes. Lower heatmap depicts single cells from good and bad old fibroblast cultures.

**Extended Data Figure 3.**
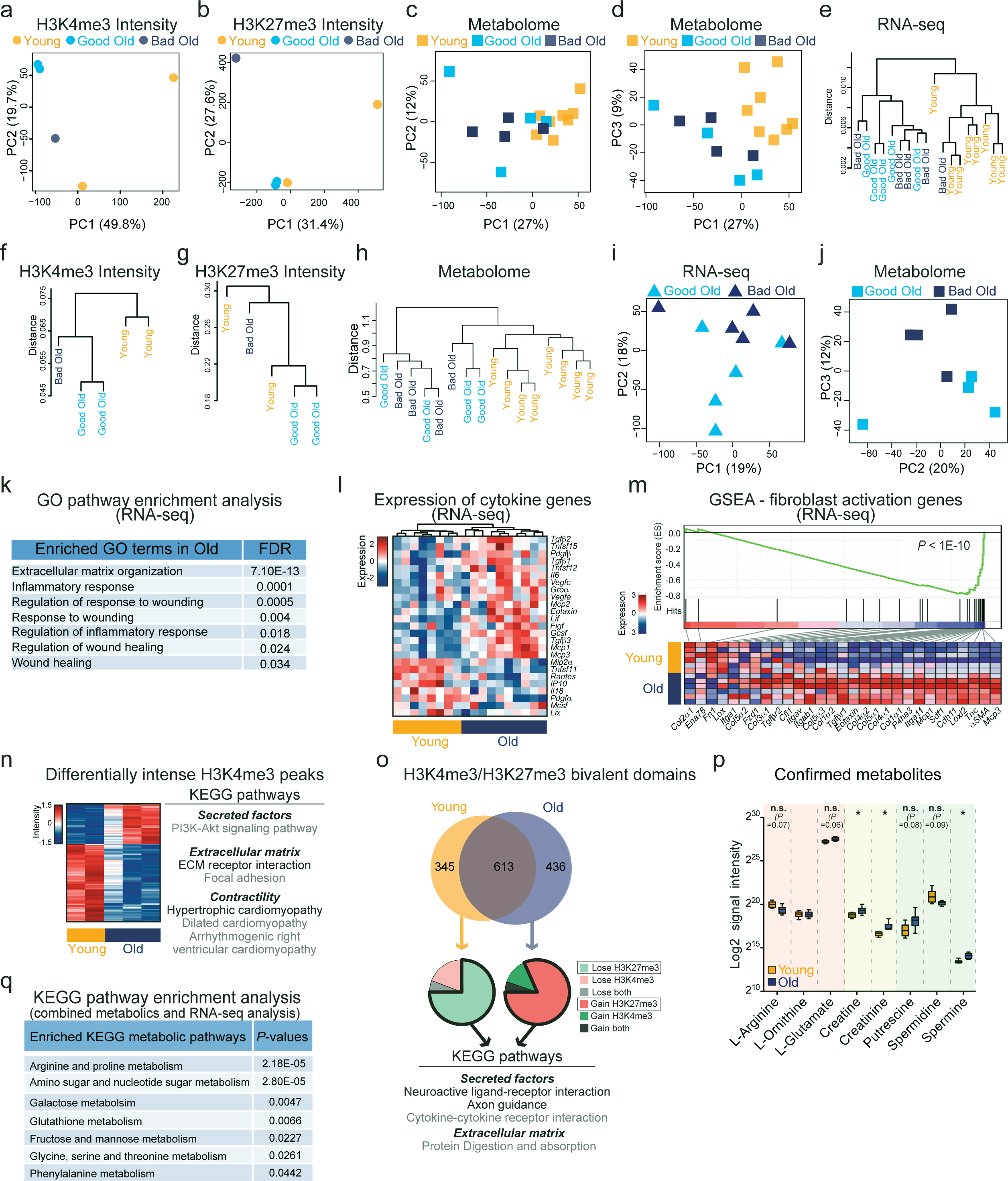
Old fibroblasts exhibit distinct transcriptomic, epigenomic and metabolomic profiles compared to young fibroblasts. Using a multi-omic approach, young and old fibroblasts, including those that reprogrammed well (Good Old) or poorly (Bad Old), were profiled in regard to their transcriptome (RNA-seq), epigenome (ChIP-seq of several histone marks) and metabolome (untargeted metabolomics). **a**, Principal component analysis (PCA) of all H3K4me3 peak intensities from young (n=2), good old (n=2) and bad old (n=1) ear fibroblast cultures from one cohort. Principal components (PC) 1 and 2 are shown. **b**, PCA of all H3K27me3 peak intensities from young (n=2), good old (n=2) and bad old (n=1) ear fibroblast cultures from one cohort. PC 1 and 2 are shown. **c**, PCA of all metabolic features of young (n=8), good old (n=4) and bad old (n=4) ear fibroblast cultures from one cohort. PC 1 and 2 are shown. **d**, PCA of all metabolic features of young (n=8), good old (n=4) and bad old (n=4) ear fibroblast cultures from one cohort. PC 1 and 3 are shown. **e**, Unsupervised hierarchical clustering of transcriptomes of young (n=8), good old (n=5) and bad old (n=5) ear fibroblast cultures from 3 cohorts. The hierarchical clustering (average linkage) was performed using correlation-based dissimilarity (Pearson’s) as distance measure and average for linkage analysis. Y-axis indicates the similarity between samples. **f**, Unsupervised hierarchical clustering of all H3K4me3 peaks from young (n=2), good old (n=2) and bad old (n=1) ear fibroblast cultures from one cohort. The hierarchical clustering was performed as in e. **g**, Unsupervised hierarchical clustering of all H3K27me3 peaks from young (n=2), good old (n=2) and bad old (n=1) ear fibroblast cultures from one cohort. The hierarchical clustering was performed as in e. **h**, Unsupervised hierarchical clustering of metabolic profiles of young (n=8), good old (n=4) and bad old (n=4) ear fibroblast cultures from one cohort. The hierarchical clustering was performed as in e. **i**, PCA of whole transcriptomes from RNA-seq data of Good and Bad Old ear fibroblast cultures. PC 1 and 2 are shown. **j**, PCA of all metabolic features of Good Old (n=4) and Bad Old (n=4) ear fibroblast cultures. PC 2 and 3 are shown. **k**, Table listing a subset of GO-terms that were enriched in the old transcriptomes, with corresponding FDR-adjusted *P*-values. For a complete list of GO terms, see Supplementary Table 2d. **l**, Heatmap of a subset of cytokine genes that are associated with young or old fibroblast cultures *in vitro*. Expression is depicted as log2 VST normalized read counts, and scaled row-wise. The scale for expression fold changes is indicated on the left. **m**, Gene Set Enrichment Analysis (GSEA) comparing transcriptional profiles of young and old fibroblasts, in regard to genes that have previously been associated with fibroblast activation (see Supplementary Table 2f). The upper panel shows the enrichment score plot corresponding to fibroblast activation (*P* < 1E-10). The lower heatmap shows expression of the genes within the fibroblast activation gene set, where expression is depicted as VST-normalized read counts (derived from DESeq2), and scaled row-wise. The scale for expression fold changes is indicated on the left. **n**, The left heatmap shows the H3K4me3 peaks within promoters regions that exhibit a significant difference in intensity with age, assessed by Diffbind. Expression is depicted as log2 VST (variance stabilizing transformation, DESeq2) normalized read counts, and scaled row-wise. The scale for expression fold changes is indicated on the left. The right panel depicts a subset of KEGG pathways enriched by the genes associated with these peaks, and are color-coded for significance (black = FDR < 0.05, grey = FDR < 0.15). These pathways are further grouped into two classes: Extracellular matrix and Contractility. For a complete list of significant KEGG terms, see Supplementary Table 2h. **o**, The upper Venn diagram depicts the overlap of bivalent domains within promoters regions of young and old fibroblasts. In this analysis, the H3K4me3 and H3K27me3 peaks that are consistently present in young samples were compared to the ones that are consistently present in old samples (see Experimental Procedures). The middle pie charts show how the unique bivalent domains in young fibroblasts change in old fibroblasts (left pie chart), and vice versa (right pie chart). The bottom panel depicts a subset of KEGG pathways enriched by the genes associated with the bivalent domains that change with age, and are color-coded for significance (black = FDR < 0.05, grey = FDR < 0.15). The pathway enrichment analysis was restricted to the bivalent domains in young that lose H3K27me3 in old, and the H3K4me3 peaks in young that gain H3K27me3 in old, as these domains are likely to alter expression of their associated genes. These pathways are further grouped into two classes: Secreted factors and Extracellular matrix. For a complete list of KEGG terms, see Supplementary Table 2k-n. **p**, Boxplots showing the log2 signal intensities of selected metabolites in the arginine and proline pathway from the metabolic profiling of young and old fibroblasts cultures at passage 3, whose identity were confirmed using commercially available standards. The boxplots depict the median and interquartile range, with whiskers indicating minimum and maximum values. *P-* values were calculated using a two-tailed Wilcoxon rank sum test and adjusted for multiple hypothesis testing using a q-value correction. * *P* < 0.05. n.s. = not significant. Note that multiple hypothesis correction was performed on all acquired metabolic features together and not solely on the depicted metabolites. **q**, Pathway analysis of all the putatively identified metabolites that were significantly different between young and old fibroblasts, as well as the differentially expressed genes (FDR < 0.05, absolute fold change > 1.5), using the MetaboAnalyst online tool^98^. Note that MetaboAnalyst online tool does not provide multiple-hypothesis corrected *P-* values.

Old fibroblasts exhibited a significant enrichment for transcriptional pathways involving secreted factors (e.g. cytokine signaling), extracellular matrix (ECM) components, contractility-related features, inflammatory and wound healing responses (Fig. 2d, Extended Data Fig. 3k-l, Supplementary Table 2b-e). Together, these features are characteristic of ‘activated fibroblasts’ (also termed myofibroblasts), which are known to be critical for tissue repair after injury but can also lead to fibrosis^4-6,51^. Consistently, the ‘Fibroblast activation’ gene set was enriched in the old fibroblast transcriptome (Extended Data Fig. 3m, Supplementary Table 2f). In line with the transcriptomic data, age-dependent changes in epigenomic landscape (e.g. H3K4me3 intensity and breadth) also revealed enrichment for pathways involved in activated fibroblasts, such as cytokines, ECM components, and contractility-related features (Fig. 2e, Extended Data Fig. 3n-o, Supplementary Table 2g-n). Similarly, untargeted metabolomics uncovered changes in arginine and proline metabolism (Fig. 2f, Extended Data Fig. 3p-q, Supplementary Table 2o), which has been implicated in the regulation of inflammatory cytokines and ECM synthesis (Fig. 2d)^52-54^, consistent with the characteristics of the activated state. We identified several transcriptional regulators, notably the transcription factor EBF2, as increased in old fibroblasts and potentially driving the ‘activated fibroblast’ signature in these cells (Fig. 2g, Supplementary Table 2p). Primary fibroblast cultures from elderly humans also exhibited increased cytokine-related pathways, with increased expression of EBF2 (Fig. 2h, Supplementary Table 2q-r). The inflammatory signature of old fibroblasts persists through passaging but can be erased by reprogramming (Extended Data Fig. 4a-l, Supplementary Table 2s-t). Thus, old fibroblasts exhibit characteristics of fibroblast activation, with a persistent inflammatory signature, consistent with cytokine profiling.

**Extended Data Figure 4.**
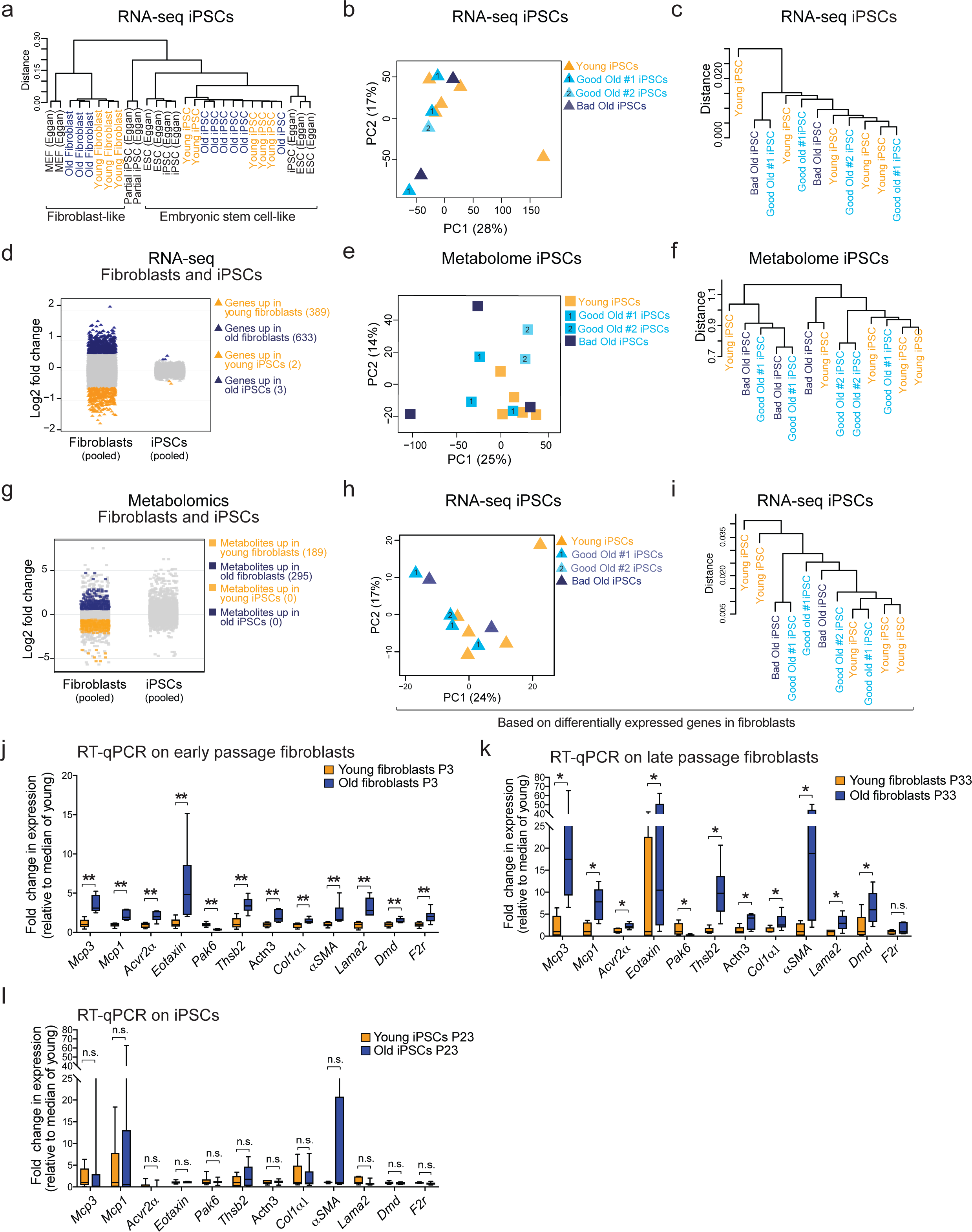
Reprogramming erases features of inflammaging and individual-to-individual variability. To test whether transcriptomic and metabolomic features of inflammaging could be erased by reprogramming, iPSC lines from young and old fibroblasts at passage 23 were profiled in regard to their transcriptome and metabolome. **a**, All iPSC lines cluster with previously established bona fide iPSC lines and ESCs. Unsupervised hierarchical clustering based on overall gene expression of transcriptomes of the indicated cell types. The hierarchical clustering (average linkage) was performed using correlation-based dissimilarity (Pearson’s) as distance measure and average for linkage analysis. Y-axis indicates the similarity between samples. **b**, Principal component analysis (PCA) of whole transcriptomes from RNA-seq data of young (n=5) and old (n=6) derived iPSC lines at passage 23. Principal components (PC) 1 and 2 are shown. **c**, Unsupervised hierarchical clustering of transcriptomes of young (n=5) and old (n=6) derived iPSC lines. The hierarchical clustering was performed as in a. **d**, Strip plot illustrating the log2 fold expression changes of all genes with age for fibroblasts (left) and iPSCs (right). Genes detected as significantly up-or down-regulated (DESeq2, FDR-values < 0.05, absolute fold change > 1.5) with age, are depicted in blue and yellow, respectively. **e**, PCA of metabolomes of young (n=5) and old (n=8) derived iPSC lines at passage 23. Untargeted metabolomic profiles were generated using ultra-high performance liquid chromatography mass spectrometry. PC 1 and 2 are shown. **f**, Unsupervised hierarchical clustering of metabolic profiles of young (n=5) and old (n=8) derived iPSC lines. The hierarchical clustering was performed as in a. **g**, Strip plot illustrating the log2 fold change in signal intensity of all metabolic features with age for fibroblasts (left) and iPSCs (right). Metabolic features detected as significantly up or down (using a two-tailed Wilcoxon rank sum test and adjusting for multiple hypotheses testing using Q-value correction) with age are depicted in blue and yellow, respectively. **h**, PCA of young (n=5) and old (n=6) derived iPSC lines at passage 23, based on solely the genes that were significantly differentially expressed between young and old at fibroblast level. PC 1 and 2 are shown. **i**, Unsupervised hierarchical clustering of transcriptomes of young (n=5) and old (n=6) derived iPSC lines, based on solely the genes that were significantly differentially expressed at fibroblast level. The hierarchical clustering was performed as in a. **j-l**, Real-time quantitative PCR (RT-qPCR) of the indicated genes in fibroblasts cultures at passage 3 (j) and 33 (k), and iPSC cultures at passage 23 (l). The genes shown represent the three major groups of features associated with fibroblast activation and that change with age in fibroblasts. Expression is presented as fold change in expression compared to median expression of young, using a box-and-whisker plot to indicate the median and interquartile range with whiskers indicating minimum and maximum values. Data shown are from total of 6 young and 6 old fibroblast cultures at passage 3 (P3), 5 young and 6 old fibroblast cultures at passage 33 (P33), 6 young and 7 old iPSC cultures at passage 23 (P23). *P-* values were calculated using a one-tailed Wilcoxon rank sum test and were adjusted for multiple hypothesis using a Benjamini-Hochberg correction. * *P* < 0.05, ** *P* < 0.01. n.s. = not significant. Note that the experiments were conducted independently in fibroblasts at passage 3, 33 and iPSCs, and therefore the statistical comparisons indicated were restricted to each independent experiment. However, a comparison between the expressions of secreted factors at passage 3 to 33 shows that expression of *Eotaxin*, but not *Mcp1* and *Mcp3,* significantly decreases upon passaging.

To determine if certain signatures of old fibroblasts correlate with the variability in reprogramming efficiency, we performed a regression analysis with the transcriptomic datasets from all samples as well as with the five best or worst reprogrammers. Both analyses showed fibroblast activation as a top feature associated with ‘good’ cellular reprogramming of old fibroblasts (other signatures included DNA replication, mismatch repair, protein export) (Fig. 2i, Supplementary Table 3a-d). Our epigenomic datasets further corroborated these findings (Extended Data Fig 5a, Supplementary Table 3e-f). Hence, the fibroblast activation signature correlates with the individual-to-individual variability in reprogramming, with a link to enhanced reprogramming efficiency.

**Extended Data Figure 5.**
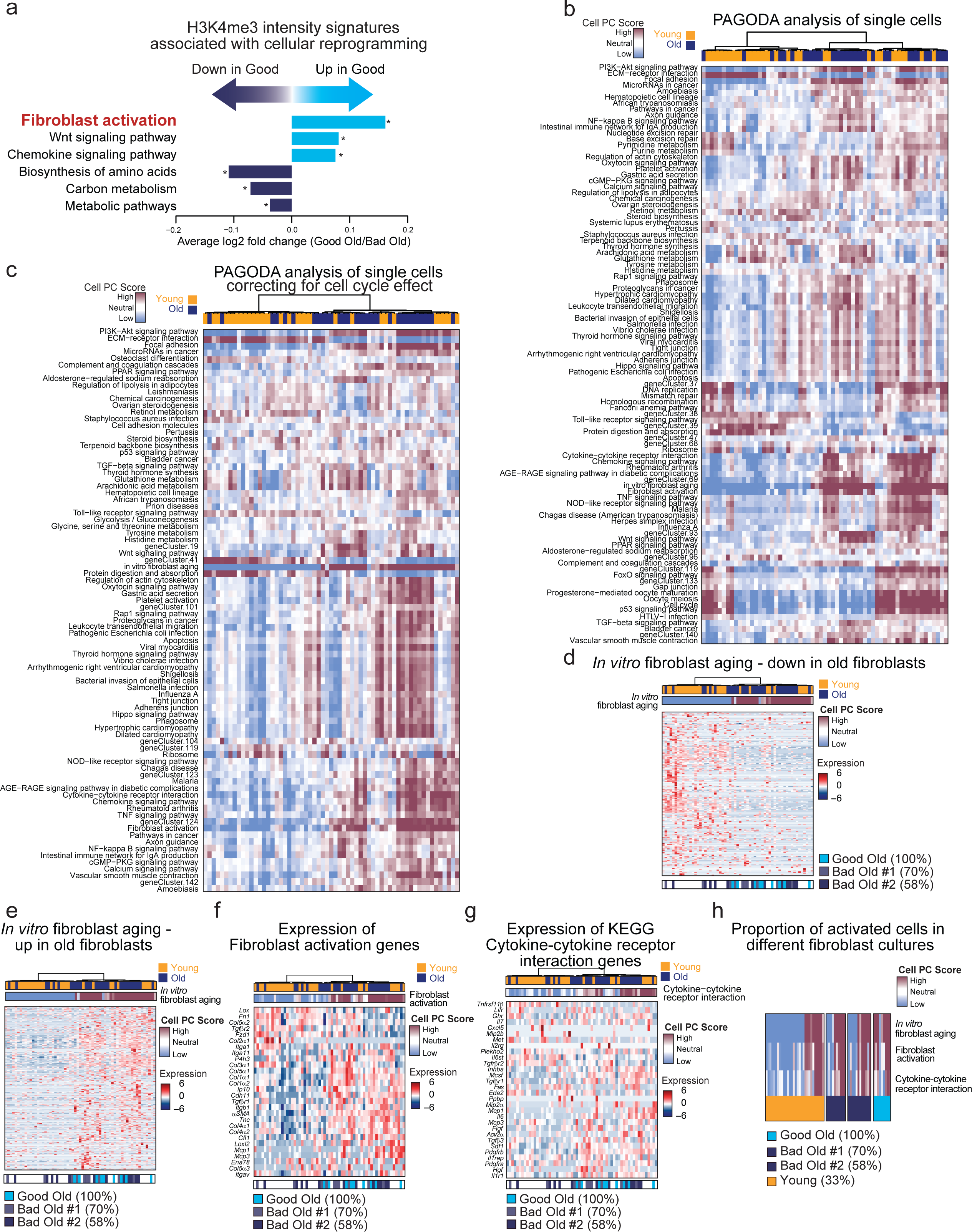
Correlation between the fibroblast activation signature and reprogramming efficiency in epigenomic and single cell RNA-seq data. **a**, Pathway enrichment analysis of KEGG pathways that are associated with enhanced (up) or reduced (down) reprogramming. These features are based on the comparison of H3K4me3 peak intensities between the top and the bottom old reprogramming culture in our datasets (see Supplementary Table 2a for more information fibroblast cultures). All depicted KEGG pathways were significantly enriched (FDR < 0.05). For a complete list of KEGG terms, see Supplementary Table 3f. **b**, Pathway and gene set overdispersion analysis (PAGODA) of single cell RNA-seq data from young and old fibroblasts at passage 3 performed using all KEGG pathways, the “*In vitro* fibroblast aging”, the “fibroblast activation”, and *de novo* gene sets. Hierarchical clustering is based on 97 significantly overdispersed gene sets and the 405 genes driving the significantly overdispersed gene sets. Upper heatmap depicts single cells from young and old fibroblast cultures. Middle heatmap shows separation of cells based on their PC scores for the significantly overdispersed gene sets. The PC scores are oriented so that high (maroon) and low (blue) values generally correspond, respectively, to increased and decreased expression of the associated gene sets. **c**, PAGODA of single cell RNA-seq data from young and old fibroblasts at passage 3 performed using all KEGG pathways, the “*In vitro* fibroblast aging”, the “fibroblast activation”, and *de novo* gene sets after accounting for cell cycle. Hierarchical clustering is based on 84 significantly overdispersed gene sets and the 248 genes driving the significantly overdispersed gene sets. Upper heatmap depicts single cells from young and old fibroblast cultures. Middle heatmap shows separation of cells based on their PC scores for the significantly overdispersed gene sets. The PC scores are oriented so that high (maroon) and low (blue) values generally correspond, respectively, to increased and decreased expression of the associated gene sets. **d**, PAGODA as described in c. The heatmap shows expression of the genes that are part of the “*In vitro* fibroblast aging” signature, and go down with age, where expression is depicted as VST normalized read counts, and scaled row-wise. The scale for expression fold changes is indicated on the right. The lower heatmap indicates the cells that originate from good and bad old cultures. **e**, PAGODA as described in c. The heatmap shows expression of the genes that are part of the “*In vitro* fibroblast aging” signature, and go up with age, where expression is depicted as VST normalized read counts, and scaled row-wise. The scale for expression fold changes is indicated on the right. The lower heatmap indicates the cells that originate from good and bad old cultures. **f**, PAGODA as described in c. The heatmap shows expression of the genes that are part of the “fibroblast activation” gene set, where expression is depicted as VST normalized read counts, and scaled row-wise. The scale for expression fold changes is indicated on the right. The lower heatmap indicates the cells that originate from good and bad old cultures. **g**, PAGODA as described in c. The heatmap shows expression of the top 30 overdispersed genes in the KEGG cytokine-cytokine receptor interaction pathway, where expression is depicted as VST normalized read counts, and scaled row-wise. The scale for expression fold changes is indicated on the right. The lower heatmap indicates the cells that originate from good and bad old cultures. **h**, PAGODA as described in c, in which cells are separated by the culture they originate from. Heatmap shows separation of cells based on their PC scores for a subset of significantly overdispersed gene sets. The PC scores are oriented so that high (maroon) and low (blue) values generally correspond, respectively, to increased and decreased expression of the associated gene sets. The lower heatmap indicates the cells that originate from good and bad old cultures.

We asked whether fibroblast cultures are heterogeneous and if the proportion of activated fibroblasts might differ between them. To this end, we performed single cell RNA-seq on 30 young and 31 old cells from 3 young and 3 old fibroblast cultures exhibiting either good (1 culture) or bad (2 cultures) reprogramming potential (Fig. 2j, Supplementary Table 3g). tSNE and Pathway and gene set overdispersion analysis (PAGODA)^55^ identified two main clusters: one predominantly composed of young cells and the other of old cells (Fig. 2k-l, Extended Data Fig. 5b-e). The subpopulation containing most old cells indeed exhibited a signature of fibroblast activation (Fig. 2k-l, Extended Data Fig. 5f), and those cells had high expression of inflammatory cytokines such as *Il6*, *Mcp1*, and *Mcp3* (Fig. 2l, Extended Data Fig. 5g). The proportion of activated fibroblasts varied between fibroblast cultures, and although relatively few cells were queried, there seems to be a higher proportion of activated cells in the good old culture (9/9 cells; Fig. 2k-l, Extended Data Fig. 5f, h) compared to the two bad old cultures (6/12 and 6/10 cells; Fig. 2k-l, Extended Data Fig. 5f, h). Collectively, these data suggest that the proportion of activated fibroblasts is linked to the variability in reprogramming between individual cultures, with a higher number of activated fibroblasts in old cultures with good reprogramming efficiency.

We next experimentally assessed the presence of activated fibroblasts in old individuals *in vitro* and *in vivo* and tested their causal role in differences in reprogramming efficiency. We first validated that old fibroblast cultures harbored a high number of cells positive for α smooth muscle actin (αSMA) a marker of activated fibroblasts (Fig. 3a, Extended Data Fig. 6a) and that these cells, unlike senescent fibroblasts, actively proliferate (Fig. 3a, Extended Data Fig. 6b) and do not exhibit signatures or markers of senescence (e.g. *p16^INK4^*) (Extended Data Fig. 6c-e). We next identified a surface marker that correlates with the activated fibroblast signature to facilitate the isolation of these cells for further experiments. *Thy1* (also known as *Cd90*) encodes a surface protein and is one of the most significant differentially expressed genes in activated fibroblasts compared to non-activated ones (Fig. 3b, Extended Table 4a-c). THY1 has been associated with populations of fibroblasts with activated features in tissues^56,57^ and with secretion of pro-inflammatory cytokines in the context of diseases^58,59^. FACS analysis using antibodies to THY1 and to the pan-fibroblast marker PDGFRα^60^ confirmed that old fibroblast cultures in general contained a higher proportion of THY1-positive/PDGFRα-positive cells (hereafter termed THY1+ cells) (Fig. 3b), and that these cells express markers of fibroblast activation (Extended Data Fig. 6f). THY1+ cells also expressed higher levels of the transcription factor EBF2 (Extended Data Fig. 6g), a potential driver of the inflammatory signature (see Fig. 2), and knockdown of EBF2 in these cells impaired expression of cytokines (e.g. *Il6*, *Eotaxin*) and other genes related to the activated state (e.g. *αSMA*, *Col1αl*) (Fig. 3c, Extended Data Fig. 6g). These experiments validate the presence of activated fibroblasts that express inflammatory cytokines in old fibroblast cultures, and further suggest that the transcription factor EBF2 drives the expression of key cytokine genes in these cells.

**Figure 3.**
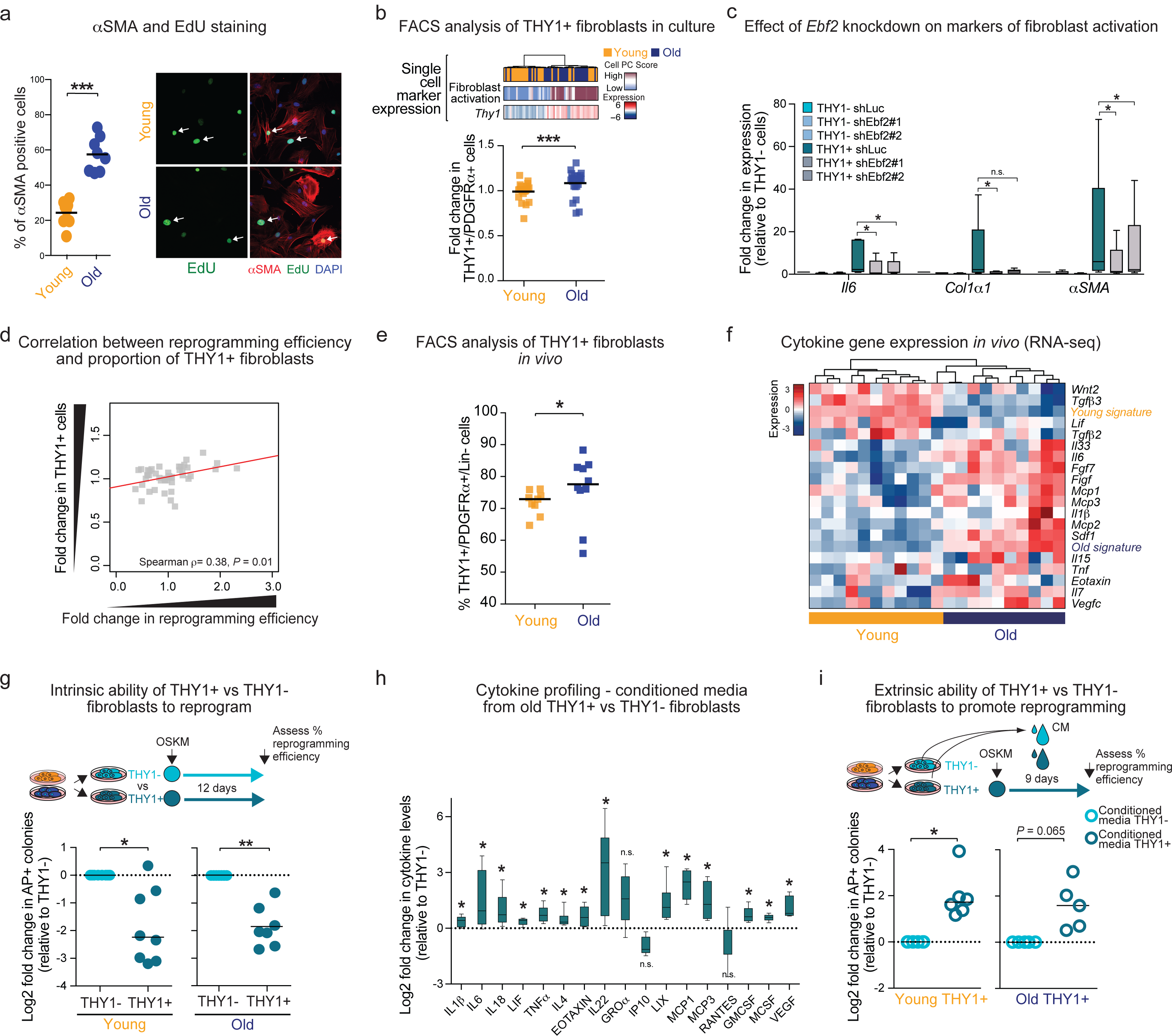
Activated fibroblasts increase with age *in vitro* and *in vivo*, and they enhance reprogramming efficiency extrinsically through secreted factors. **a**, Left panel: Quantification of the percentage of cells positive for α smooth muscle actin (αSMA), a marker of activated fibroblasts, assessed by immunofluorescence staining of young and old fibroblasts cultures at passage 3. For representative images, see Extended Data Fig. 6a. Data shown are percentage of αSMA positive cells from a total of 8 young and 8 old mice (1 cohort) combined from 1 independent experiments. Each dot represents cultures from an individual mouse. Line depicts median percentage. *P*-values were calculated using a two-tailed Wilcoxon rank sum test. *** *P* < 0.001. Right panel: Representative immunofluorescence images of young and old fibroblasts at passage 3 incubated with EdU, a base analog that incorporates during S phase, for 4h, and then stained for αSMA (red, activation marker), EdU (green, proliferation marker), and DAPI (blue, nuclei). The white arrows indicate an activated and a non-activated cell in each respective image. For quantification of EdU-positive cells, see Extended Data Fig. 6b. **b**, Upper panel: Pathway and gene set overdispersion analysis (PAGODA) of single cell RNA-seq data from young and old fibroblasts as in Fig. 2k, showing the expression of *Thy1*, a surface marker expressed mainly in old activated fibroblasts. Lower panel: Proportion of THY1+/PDGFRα+ (hereinafter referred to as THY1+) fibroblasts in young and old ear fibroblast cultures at passage 3, as measured by FACS. Results are shown as fold change in percentage of THY1+ fibroblasts relative to that of the median of young fibroblasts. Data shown are from a total of 21 young and 23 old mice from 3 cohorts of mice, combined over 3 independent experiments (for individual experiments, see Supplementary Table 7). Each dot represents cultures from an individual mouse. Line depicts median fold change. *P*-values were calculated using a two-tailed Wilcoxon rank sum test. *** *P* < 0.001. No statistical difference in variance between age groups was observed (*P* = 0.54) using the non-parametric Fligner-Killeen test. **c**, Real-time quantitative PCR (RT-qPCR) of the indicated genes in THY1−/PDGFRα+ (hereinafter referred to as THY1−) and THY1+ cells expressing the indicated shRNAs for 72h. Hprt1 was used as housekeeping gene. Expression is presented as fold change in expression compared to THY1− shLuciferase (shLuc) treated cells, using a box-and-whisker plot to indicate the median and interquartile range with whiskers indicating minimum and maximum values. Data shown are from total of 5 old mice from 1 cohort, combined over 4 independent experiments (for individual experiments, see Supplementary Table 7). P-values were calculated using one-tailed Wilcoxon signed rank test and adjusted for multiple hypothesis testing using a Benjamini-Hochberg correction. * P < 0.05. n.s. = not significant. **d**, Positive correlation between the proportion of THY1+ fibroblasts in a given culture, as quantified by FACS, and the reprogramming efficiency of the culture using Spearman rank correlation. The y-axis denotes the fold change in the proportion of THY1+ cells relative to the median of young, and x-axis denotes the fold change in reprogramming efficiency of the culture relative to the median of young. Each dot represents cells from an individual mouse. Data shown are from a total of 21 young and 23 old mice from 3 cohorts of mice, combined over 3 independent experiments (for individual experiments, see Supplementary Table 7). **e**, FACS analysis of fibroblasts freshly isolated in vivo from young and old ears, defined as live PDGFRα+/Lin-(CD45-/CD31-/EpCAM-/TER119-/TIE2-) cells, and then further subdivided into THY1+ or THY1− cells. Results are shown as percentage of THY1+/PDGFRα+/Lin-cells over PDGFRα+/Lin-cells. Data shown are from a total of 9 young and 10 old biological replicates (cells pooled from 2-3 mice) from 3 cohorts of mice, combined over 3 independent experiments (for individual experiments, see Supplementary Table 7). Each dot represents cells pooled from 2-3 mice. Line depicts median percentage. *P*-values were calculated using a two-tailed Wilcoxon rank sum test. * *P* < 0.05. No statistical difference in variance between age groups was observed (*P* = 0.14) using the non-parametric Fligner-Killeen test. Note that the % of PDGFRα+/Lin-/THY1+ is also used in Fig. 4g as basal (non-wounded). **f**, Heatmap of expression of a subset of cytokine genes from population RNA-seq of freshly isolated THY1+/PDGFRα+/Lin-or THY1−/PDGFRα+/Lin-cells from young and old ears. FACS sorting was performed as described in **e**. Two to three young or old mice were pooled together to obtain 500 cells of each population for each biological replicate. Expression is depicted as log2 VST normalized read counts, and scaled row-wise. The scale for expression fold changes is indicated on the left. Young and old signatures refer to the average expression of the genes that go significantly down or up with age, respectively, in this dataset. **g**, Reprogramming efficiency (RE) of FACS-sorted young (left panel) and old (right panel) THY1− and THY1+ fibroblasts at passage 4-6, assessed using alkaline phosphatase (AP) staining. Individual cultures were FACS-sorted using antibodies to THY1 and PDGFRα, and comparisons were made between pairs of THY1− and THY1+ from the same original culture. Results are shown as log2 fold change in RE of the cells over that of THY1− fibroblasts. Data shown are from a total of 8 young and 7 old mice from 3 cohorts of mice, combined over 3 independent experiments (for individual experiments, see Supplementary Table 7). Note that 1 old culture was used in 2 independent experiments. In this case, an average of the resultant measurements was determined so as to not inflate the statistical power. Each dot represents cells from an individual mouse. Line marks median log2 fold change. *P*-values were calculated using two-tailed Wilcoxon signed rank test. * *P* < 0.05, ** *P* < 0.01. **h**, Inflammatory cytokine profiles of conditioned media collected from cultures of THY1− and THY1+ old fibroblasts at passage 4-6. Individual old cultures were FACS-sorted using antibodies to THY1 and PDGFRα, and comparisons were made between pairs of THY1− and THY1+ from the same original culture. Results are shown as log2 fold change in mean fluorescence intensity (MFI) over THY1− fibroblasts, using a box-and-whisker plot to indicate the median and interquartile range with whiskers indicating minimum and maximum values. Data shown are from a total of 6 old mice from 3 cohorts of mice, combined over 3 independent experiments (for individual experiments, see Supplementary Table 7).*P-* values: one-tailed Wilcoxon signed rank test and adjusted for multiple hypothesis testing using a Benjamini-Hochberg correction. * *P* < 0.05, n.s. = not significant. **i**, RE, assessed as in g, of FACS-sorted young (left panel) and old (right panel) THY1+ fibroblasts at passage 4-6 treated with fresh CM every day starting from day one after infection with OSKM. CM was collected every day from the THY1− or THY1+ fibroblasts from the same original culture. Results are shown as log2 fold change in the RE relative to the RE of THY1− fibroblasts treated with CM from THY1− fibroblasts. Data shown are from a total of 6 young and 5 old mice from 2 cohorts of mice, combined from 4 independent experiments (for individual experiments, see Supplementary Table 7). Note that 1 young culture was used in 2 independent experiments. In this case, an average of the resultant measurements was determined so as to not inflate the statistical power. Each dot represents cells from an individual mouse. Line marks median log2 fold change. *P*-values were calculated using two-tailed Wilcoxon signed rank test. *P*-value was not significant for the old (*P* = 0.065) because of low sample number (5). * *P* < 0.05, CM = conditioned media.

**Extended Data Figure 6.**
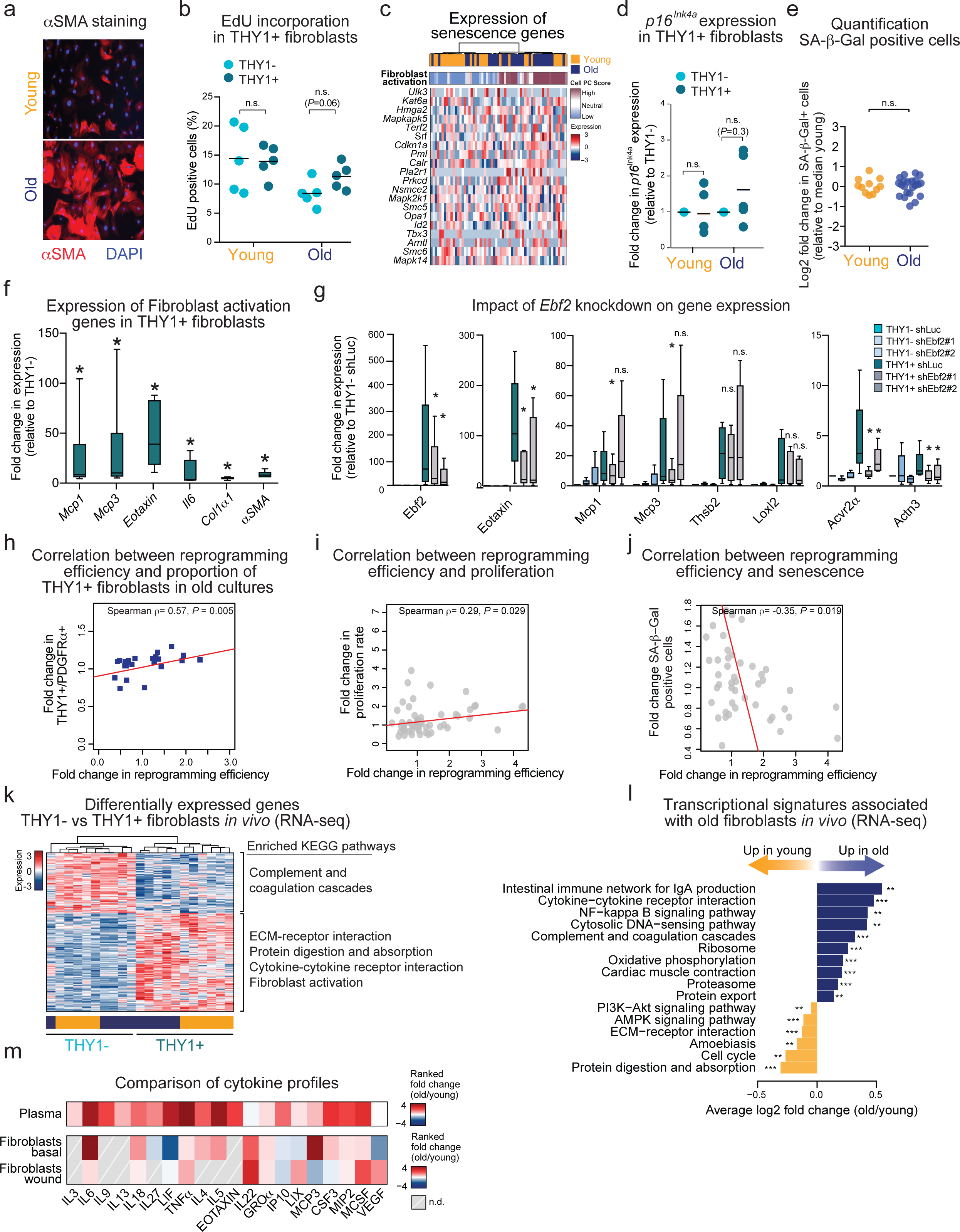
Old fibroblast cultures are enriched for activated fibroblasts that actively proliferate and that are associated with enhanced reprogramming. **a**, Old fibroblast cultures are enriched for α smooth muscle actin (αSMA) positive cells, a marker of activated fibroblasts. Representative immunofluorescence (IF) images of young and old fibroblasts at passage 3 stained for αSMA. Nuclei were stained with DAPI. For quantification see Fig. 3a. **b**, Activated fibroblasts are actively proliferating. FACS quantification of the percentage of EdU-positive THY1−/PDGFRα+ and THY1+/PDGFRα+ cells in young and old cultures at passage 3. Data are from a total of 5 young and 5 old fibroblast cultures combined from 4 independent experiments. Each dot represents cells from an individual mouse. Line indicates median percentage. *P*-values were calculated using a two-tailed Wilcoxon rank sum test. n.s. = not significant. **c**, PAGODA of single cell RNA-seq data from young and old fibroblasts performed using all KEGG pathways, the “*In vitro* fibroblast aging”, the “fibroblast activation”, and *de novo* gene sets after accounting for cell cycle. Hierarchical clustering is based on 84 significantly overdispersed gene sets and the 248 genes driving the significantly overdispersed gene sets. Upper heatmap depicts single cells from young and old fibroblast cultures. Middle heatmap shows separation of cells based on their PC scores for the “Fibroblast activation” signature. The PC scores are oriented so that high (maroon) and low (blue) values generally correspond, respectively, to increased and decreased expression of the associated gene sets. Lower heatmap shows expression of the genes that are part of the “GO Cellular senescence” gene set, where expression is depicted as VST normalized read counts, and scaled row-wise. The scale for expression fold changes is indicated on the right. None of the senescence gene sets (see Extended Data Fig. 5b-c) were significantly overdispersed using PAGODA. **d**, Real-time quantitative PCR of *p16^INK4^* a expression in cultures of THY1− and THY1+ young and old cells at passage 4-6. Individual young and old cultures were FACS sorted into THY1− and THY1+ populations, and comparisons were made between pairs from the same original culture. Results are presented as fold change in expression of *p16 INK4a* between THY1− and THY1+ populations. *Hprt1* was used as housekeeping gene. Data shown are from a total of 5 young and 5 old fibroblast cultures combined from 6 independent experiments. One young and 3 old cultures were used in 2-3 independent experiments. In this case, an average of the resultant measurements was determined so as to not inflate the statistical power. Each dot represents cells from an individual mouse. Line indicates median fold change in expression. *P*-values were calculated using a two-tailed Wilcoxon signed rank test. **e**, The percentage of SA-β-galactosidase positive cells does not significantly differ between young and old fibroblast cultures at passage 3 (*P* value = 0.81, two-tailed Wilcoxon rank sum test). Results are shown as log2 fold change in SA-β-galactosidase positive cells over the median of SA-β-galactosidase positive cells in young fibroblasts. Data shown are from 3 independent experiments, using a total of 11 young and 22 old mice from 3 cohorts. Each dot represents cells from an individual mouse. Line indicates median log2 fold change in expression. *P*-values were calculated using a two-tailed Wilcoxon ranked sum test. n.s. = not significant. **f**, THY1+ fibroblasts exhibit features of activated fibroblasts. Real-time quantitative PCR of the indicated genes in old cultures of THY1+ and THY1− old cells at passage 4-6. Individual old cultures were FACS sorted into THY1+ and THY1− populations, and comparisons were made between pairs from the same original culture. Results are presented as fold change in expression between THY1+ and THY1− populations originating from the same culture, using a box-and-whisker plot to indicate the median and interquartile range with whiskers indicating minimum and maximum values. *Hprt1* was used as housekeeping gene. Data shown are from a total of 6 old mice from 1 cohort combined over 3 independent experiments (for individual experiments, see Supplementary Table 7). *P*-values were calculated using a one-tailed Wilcoxon signed rank test and adjusted for multiple hypothesis testing using a Benjamini-Hochberg correction. * *P* < 0.05. **g**, Real-time quantitative PCR of the indicated genes in THY1−/PDGFRα+ and THY1+/PDGFRα+ cells treated with the indicated shRNA constructs for 72h. Results are presented as fold change in expression compared to THY1−/PDGFRα+ shLuciferase (shLuc) treated cells, using a box-and-whisker plot to indicate the median and interquartile range with whiskers indicating minimum and maximum values. *Hprt1* was used as housekeeping gene. Data shown are from a total of 5 old mice from 1 cohort combined over 4 independent experiments (for individual experiments, see Supplementary Table 7). *P*-values were calculated using a one-tailed Wilcoxon signed rank test and adjusted for multiple hypothesis testing using a Benjamini-Hochberg correction. * P < 0.05. n.s. = not significant. **h**, Correlation between the proportion of THY1+/PDGFRα+ fibroblasts in a given old culture and the reprogramming efficiency of the culture. There is a positive correlation (Spearman rank correlation, ρ = 0.57, *P*-value < 0.005) between these two features. The y-axis denotes the fold change in the proportion of THY1+/PDGFRα+ cells relative to the median of young, and x-axis denotes the fold change in reprogramming efficiency of the culture relative to the median of young. Each dot represents cells from an individual mouse. Data shown are from a total of 23 old mice from 3 cohorts of mice, combined over 3 independent experiments (for individual experiments, see Supplementary Table 7). **i**, The correlation between the proliferation rate of a given fibroblast culture and the reprogramming efficiency of the culture. Proliferation rate was determined by calculating the growth slope of young, middle-aged and old ear fibroblast cultures at passage 3. There is a significant positive correlation (Spearman rank correlation, ρ = 0.29, *P*-value = 0.029) between these two features. The y-axis denotes the fold change in the proliferation rate relative to the median of young, and x-axis denotes the fold change in reprogramming efficiency of the culture relative to the median of young. Each dot represents cells from an individual mouse. Data shown are from a total of 15 young, 10 middle-aged and 27 old mice from 4 cohorts, combined from 4 independent experiments. **j**, The correlation between the percentage of SA-β-galactosidase-positive cells of a given fibroblast culture and the reprogramming efficiency of the culture. Senescence rate was assessed by SA-β-galactosidase staining of young, middle-aged and old ear fibroblast cultures at passage 3. There was a significant negative correlation (Spearman rank correlation, ρ = −0.35, *P*-value = 0.019) between these two features. The y-axis denotes the fold change in the percentage of SA-β-galactosidase-positive cells relative to the median of young, and x-axis denotes the fold change in reprogramming efficiency of the culture relative to the median of young. Each dot represents cells from an individual mouse. Data shown are from a total of 11 young, 11 middle-aged and 22 old mice from 3 cohorts, combined from 3 independent experiments. **k**, Heatmap of significantly differentially expressed genes (FDR-values < 0.05, absolute fold change > 1.5) between freshly FACS sorted THY1+/PDGFRα+/Lin-and THY1−/PDGFRα+/Lin-cells from young and old ears. The panel to the right indicates the KEGG pathways that these genes are enriched for. Expression is depicted as log2 VST normalized read counts, and scaled row-wise. The scale for expression fold changes is indicated on the left. All depicted KEGG pathways were significantly enriched (FDR < 0.05). For a complete list of KEGG terms, see Supplementary Table 4e. **l**, Pathway enrichment analysis of KEGG pathways that are associated with aging in freshly FACS sorted THY1+/PDGFRα+/Lin-and THY1−/PDGFRα+/Lin-cells from young and old ears. All depicted KEGG pathways were significantly enriched (FDR < 0.05). For a complete list of KEGG terms, see Supplementary Table 4g. ** *P* < 0.01, *** *P* < 0.001. **m**, Heatmap of cytokines that are detected in young and old plasma (see Fig. 1b) and cytokine transcripts in non-wounded and wounded THY+/PDGFRα+/Lin-cells. Upper panel: ranked fold change (old/young) in levels of the indicated cytokines in plasma (see Fig. 1b). Lower panel: ranked fold change (old/young) in expression for the indicated cytokines freshly FACS-sorted THY1+/PDGFRα+/Lin-and THY1−/PDGFRα+/Lin-cells from young and old ears. See Experimental Procedures for calculation of ranked fold changes. n.d. = expression not detected. Note that gene expression related to wounded fibroblasts is from the data sets described in Fig. 4e.

Importantly, FACS analysis of fibroblasts cultures corroborated the positive correlation between the proportion of THY1+ activated cells in a given culture and the propensity of the culture to reprogram (Spearman correlation, ρ = 0.38) (Fig. 3d), especially among old cultures (Spearman correlation, ρ = 0.57) (Extended Data Fig. 6h). Hence, even though the proportion of THY1+ cells is not more variable in the old compared to young cultures (Fig. 3b), their increased proportion, particularly in old cultures, correlates with enhanced reprogramming. Reprogramming efficiency also correlated positively with proliferation (as predicted by our multi-omic analysis) and negatively with senescence (Extended Data Fig. 6i-j). These analyses confirm that activated fibroblasts that express cytokines are more numerous in old fibroblast cultures, and that their number is correlated with reprogramming efficiency.

We then asked if activated fibroblasts are also present *in vivo* and if fibroblast composition changes in old tissues. FACS analysis of the ears from young and old mice revealed a significantly higher proportion of THY1+ fibroblasts in old mice (Fig. 3e). Transcriptomic analysis by RNA-seq of THY1+ and THY1− fibroblasts isolated freshly from old ears revealed enrichment for activated fibroblast signatures in THY1+ fibroblasts (Extended Data Fig. 6k, Supplementary Table 4d-e) and a general increase in the expression of inflammatory cytokines in fibroblasts with age (Fig. 3f, Extended Data Fig. 6l, Supplementary Table 4f-g). Some of these cytokines are also present in old plasma (e.g. IL6) (Extended Data Fig. 6m). These findings are consistent with our *in vitro* observations that aging is associated with an increased proportion of fibroblasts with activated features and an inflammatory signature that partially overlaps with the aging inflammatory milieu.

We next determined how activated fibroblasts influence reprogramming efficiency, and whether intrinsic or extrinsic factors are involved. Surprisingly, even though the proportion of THY1+ fibroblasts is correlated to enhanced reprogramming efficiency, the intrinsic reprogramming efficiency of THY1+ fibroblasts (activated fibroblasts) isolated by FACS from old mice was ∼4-fold lower than that of THY1− fibroblasts (Fig. 3g). This was confirmed using a non-lentiviral reprogramming approach (Extended Data Fig. 7a). We next examined whether activated fibroblasts could impact reprogramming efficiency by secreting extrinsic factors. Cytokine profiling reveals that the conditioned media from FACS-purified THY1+ fibroblasts shows increased inflammatory profile compared to that of THY1− cells (Fig. 3h, Fig. 7b, Supplementary Table 4h). Interestingly, conditioned media from THY1+ cells boosted the reprogramming of fibroblasts by ∼4 fold (whether they were THY1+ or THY1−) compared to conditioned media from THY1− cells (Fig. 3i, Extended Data Fig. 7c). Together, these results indicate that activated fibroblasts, although intrinsically deficient at undergoing cellular reprogramming, can boost reprogramming of other cells by secreting factors into the media.

**Extended Data Figure 7.**
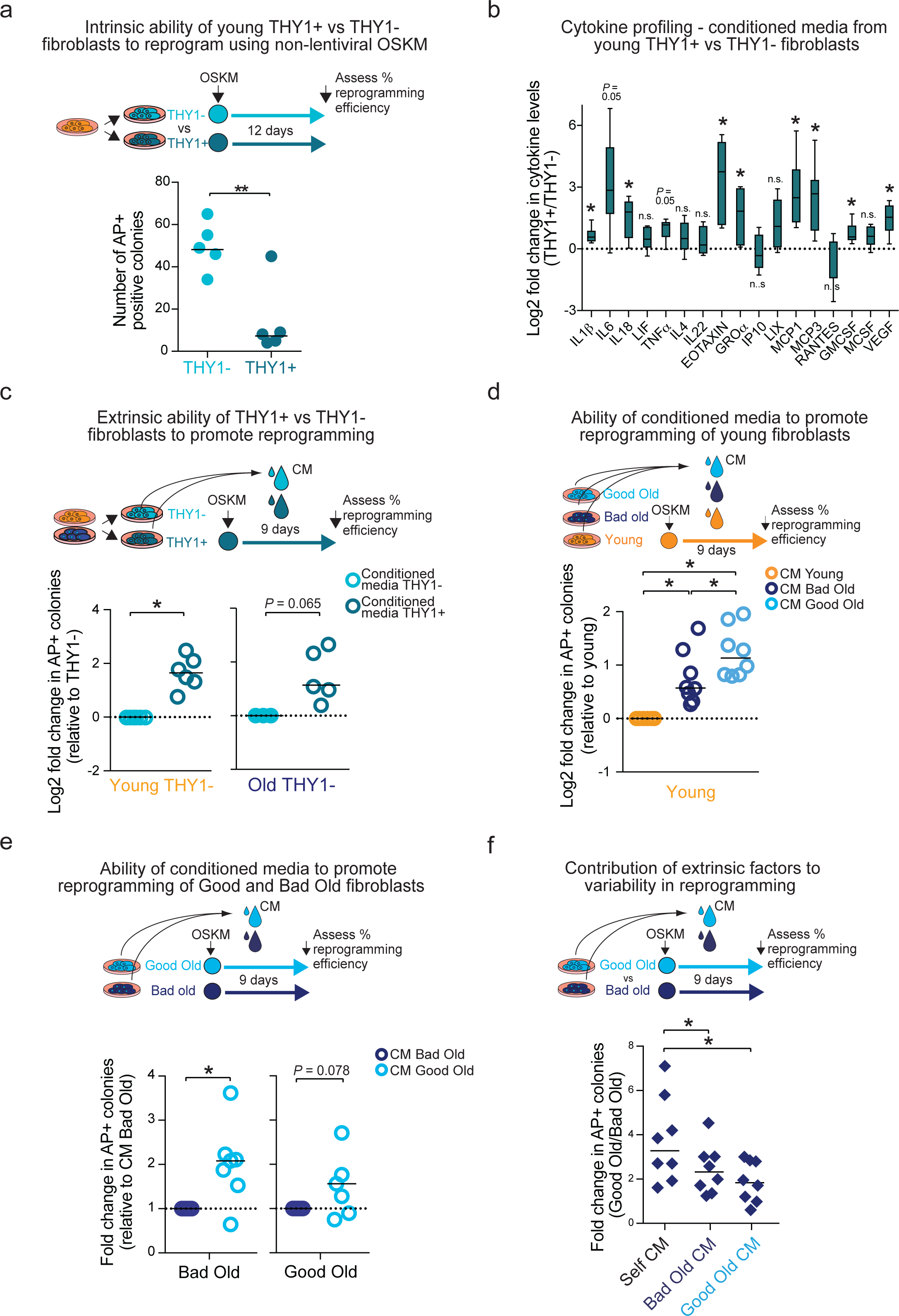
THY1+ fibroblasts are intrinsically poor at reprogramming but facilitate the process extrinsically via secretion of cytokines. **a**, Reprogramming efficiency of FACS-sorted young THY1+ and THY1− fibroblasts at passage 4-6, assessed using alkaline phosphatase (AP) staining. Reprogramming was induced using a non-lentiviral piggyBac transposon system approach. Results are shown as number of AP+ colonies. Data shown are from a total of 5 young mice from 2 cohorts, combined over 3 independent experiments (for individual experiments, see Supplementary Table 7). Each dot represents cells from an individual mouse. Line marks median log2 fold change. *P*-values were calculated using a one-tailed Wilcoxon rank sum test. ** *P* < 0.01. **b**, Inflammatory cytokine profiles of conditioned media collected from cultures of young THY1+ and THY1− fibroblasts at passage 4-6. Individual young cultures were FACS-sorted using antibodies to THY1 and PDGFRα, and comparisons were made between pairs of THY1+ and THY1− from the same original culture. Results are shown as log2 fold change in mean fluorescence intensity (MFI) over THY1− fibroblasts, using a box-and-whisker plot to indicate the median and interquartile range with whiskers indicating minimum and maximum values. Data shown are from a total of 6 young mice from 2 cohorts of mice, combined over 2 independent experiments (for individual experiments, see Supplementary Table 7).*P*-values were calculated using a one-tailed Wilcoxon signed rank test and adjusted for multiple and adjusted for multiple hypothesis testing using a Benjamini-Hochberg correction. * *P* < 0.05. n.s. = not significant. **c**, Reprogramming efficiency (RE), assessed as in a, of young (left panel) and old (right panel) THY1− fibroblasts at passage 4-6 treated with fresh CM every day starting from day one post-infection. CM was collected every day from the THY1− or THY1+ fibroblasts from the same original culture. Results are shown as log2 fold change in the RE relative to the RE of THY1− fibroblasts treated with CM from THY1− fibroblasts. Data shown are from a total of 6 young and 5 old mice from 2 cohorts of mice, combined from 4 independent experiments. Note that 1 young culture was used in 2 independent experiments. In this case, an average of the resultant measurements was determined so as to not inflate the statistical power. Each dot represents cells from an individual mouse. Line marks median log2 fold change. *P*-values were calculated using a two-tailed Wilcoxon rank sum test. *P*-value was not significant for the old (*P* = 0.065) because of low sample number (5). * *P* < 0.05, CM = conditioned media. **d**, RE, assessed as in a, of young fibroblasts at passage 3 treated with CM from day one post-infection. CM was collected from young, good old or bad old fibroblast cultures. Results are shown as fold change in the RE relative to the RE of young fibroblasts treated with young CM. Data shown are from 8 young mice (recipient cells) from 3 cohorts of mice, combined from 4 independent experiments (for individual experiments, see Supplementary Table 7). Each dot represents cells from an individual mouse. Line marks median log2 fold change. *P*-values were calculated using a two-tailed Wilcoxon signed rank test and were adjusted for multiple hypothesis testing using Benjamini-Hochberg correction. * *P* < 0.05, CM = conditioned media. **e**, RE, assessed as in a, of bad old (left panel) or good old (right panel) fibroblast cultures at passage 3 treated with CM from day one post-infection. CM was collected from good or bad old fibroblast cultures. Results are shown as fold change in the RE relative to the RE of the indicated old fibroblasts treated with bad old CM. Data shown are from 7 bad and 6 good old mice (recipient cells) from 2 cohorts of mice, combined from 5 independent experiments. Each dot represents the fold difference in RE between a unique pair of good and bad old cultures. Line marks median fold change. *P*-values were calculated using a two-tailed Wilcoxon signed rank test.* *P* < 0.05, CM = conditioned media. **f**, RE, assessed as in a, of good and bad old fibroblast cultures at passage 3. Pairs of good and bad old fibroblast cultures were treated either with CM from the corresponding bad old (bad old CM) or with CM from the corresponding good old (good old CM). Data shown are from a total of 6 good and 7 bad old cultures isolated from a total of 13 old mice from 2 cohorts, combined from 5 independent experiments. The cultures were combined to generate 8 unique pairs of good and bad old cultures. Note that 1 good and 1 bad old cultures were used in 2 different pairs. Each dot represents the fold difference in RE between a unique pair of good and bad old cultures. Line marks median fold change. *P*-values were calculated using a two-tailed Wilcoxon signed rank test. * *P* < 0.05, CM = conditioned media.

Could factors secreted by activated fibroblasts contribute to the variability in reprogramming efficiency between individual old cultures? We compared the effect of conditioned media from ‘good’ old fibroblasts (high proportion of activated fibroblasts) to media from ‘bad’ old fibroblasts (low proportion of activated fibroblasts) on reprogramming efficiency. We used fibroblasts from young mice as recipient cells to avoid interference from intrinsic factors. Good old conditioned media enhanced reprogramming efficiency more than bad old conditioned media (Extended Data Fig. 7d). We also performed conditioned media swapping experiments to determine the difference in reprogramming efficiency between good and bad old fibroblast cultures in the context of their own conditioned media or swapped conditioned media (Fig. 4a, Extended Data Fig. 7e-f). Interestingly, when good and bad old fibroblast pairs were reprogrammed in the context of the swapped conditioned media, the difference in reprogramming efficiency was reduced by 63% (Fig. 4a). Thus, extrinsic factors play a role in the variability in reprogramming efficiency between old cultures (although intrinsic factors may also contribute).

**Figure 4.**
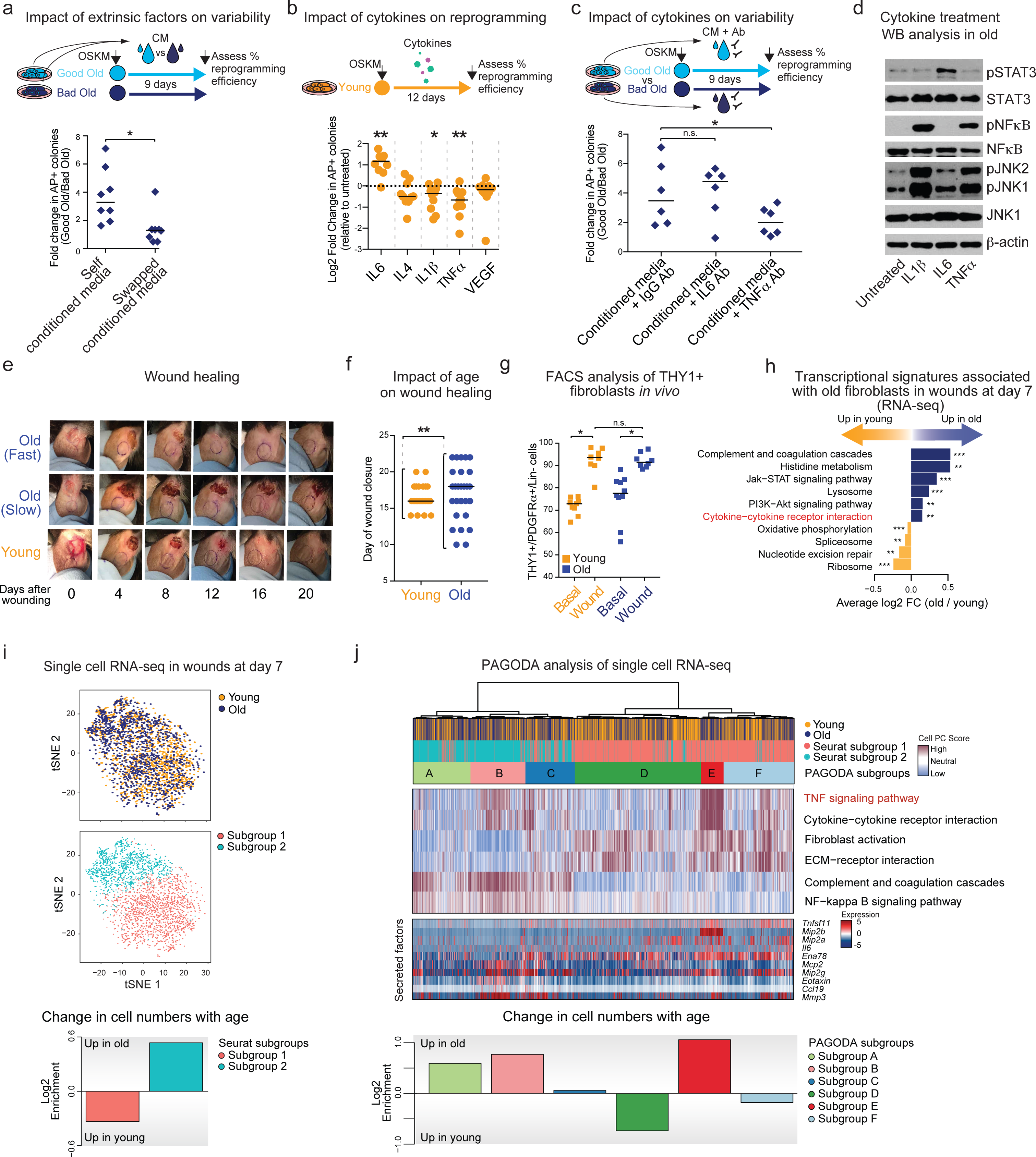
Inflammatory cytokines drive the inter-individual variability in reprogramming and may contribute to the variability in wound healing with age. **a**, Reprogramming efficiency (RE) assessed by alkaline phosphatase (AP) staining of good and bad old fibroblast cultures at passage 3. Pairs of good and bad old fibroblast cultures were treated either with their own CM (self CM) or with CM from the corresponding partner (swapped CM). Data shown are from a total of 6 good and 7 bad old cultures isolated from a total of 13 old mice from 2 cohorts, combined from 5 independent experiments. The cultures were combined to generate 8 unique pairs of good and bad old cultures. Note that 1 good and 1 bad old cultures were used in 2 different pairs. Each dot represents the fold difference in RE between a unique pair of good and bad old cultures. Line marks median fold change. *P*-values were calculated using a two-tailed Wilcoxon signed rank test. * *P* < 0.05, CM = conditioned media. **b**, RE, assessed as in a, of young cells at passage 3. Cells were treated with the indicated cytokines from day one post-infection at the concentration of 10ng/mL. Results are shown as log2 fold change in RE over RE of untreated cells. Data shown are from a total of 10 young mice from 3 cohorts, combined over 3 independent experiments. Each dot represents cells from an individual mouse. Lines depict median log2 fold change. *P*-values were calculated using a two-tailed Wilcoxon signed rank test and were adjusted for multiple hypothesis testing using Benjamini-Hochberg correction. * *P* < 0.05, ** *P* < 0.01. **c**, Fold change in RE, assessed as in a, between unique pairs of good and bad old fibroblast cultures. Pairs of good and bad old fibroblast cultures were treated with their own CM pre-treated with the indicated blocking antibodies. Data shown are from a total of 5 good and 5 bad old cultures isolated from a total of 10 old mice from 2 cohorts, combined over 4 independent experiments. The cultures were combined to generate 6 unique pairs of good and bad old cultures. Note that 1 good and 1 bad old culture were used in 2 different pairs. Each dot represents the fold difference in RE between a unique pair of good and bad old cultures. Line marks median fold change. *P*-values were calculated using a two-tailed Wilcoxon signed rank test. * *P* < 0.05, CM = conditioned media, n.s. = not significant. Line marks median fold change. **d**, Western blot analysis using the indicated antibodies, of old fibroblasts at passage 3, treated with the indicated cytokines at the concentration of 10 ng/mL for 30 min. **e**, Example images of ear wounds of young mice, fast-healing old mice (Old Fast) and slow-healing old mice (Old Slow) throughout the wound healing processes. Ink circles depict initial size of wounds. **f**, Wound healing in the ears of young (3-4 months) and old (24-26 months) mice. Full thickness wounds were induced on the dorsal side of both ears (see Experimental Procedures for further details), and the size of the wounds was monitored by imaging ear wounds every second day for 20 days. For each mouse, the average of both ear wounds was calculated. Data shown are from a total of 26 young and 28 old mice from 2 cohorts, combined over 2 independent experiments (for individual experiments, see Supplementary Table 7). Each dot represents an individual mouse. Line marks median day of wound closure. Statistical differences in variance between the age groups were calculated using the non-parametric Fligner-Killeen test. ** *P* < 0.01. **g**, FACS analysis as described in Fig. 3e to assess the % of PDGFRα+/Lin-/THY1+ in ears of young and old mice during basal conditions and at 7 days post-induction of wounds. Results are shown as percentage of THY1+/PDGFRα+/Lin-cells over PDGFRα+/Lin-cells. Data shown are from a total of 9 young basal, 8 young wounded, 10 old basal and 8 old wounded biological replicates (cells pooled from 2-3 mice) from 3 cohorts, combined over 3 independent experiments (for individual experiments, see Supplementary Table 7). Each dot represents cells pooled from 2-3 mice. Line depicts median percentage. *P*-values were calculated using a two-tailed Wilcoxon rank sum test. * *P* < 0.05. n.s. = not significant. Note that the % of PDGFRα+/Lin-/THY1+ in young and old basal conditions is the same as in Fig. 3e. **h**, Pathway enrichment analysis based on population RNA-seq of young and old THY1−/PDGFRα+/Lin-and THY1+/PDGFRα+/Lin-cells *in vivo*, during basal conditions and 7 day post-induction of wounds. The graph shows a subset of KEGG pathways that were found to be significantly enriched (FDR < 0.05). For a complete list of pathways, see Supplementary Table 5d. ** *P* < 0.01, *** *P* < 0.001. **i**, t-SNE clustering of single cell RNA-seq data of all live high quality PDGFRα+/Lin-cells (3036 cells in total) from young and old ear wounds, 7 days post-induction of wounds. Upper panel depicts the cells colored by age. depicts the cells colored by significant clusters identified using a KNN graph-based algorithm as implemented by Seurat. Middle panel depicts the cells colored by significant clusters identified using a KNN graph-based algorithm as implemented by Seurat. Lower panel depicts the log2 fold change in the two subpopulations between young and old wounds at day 7. Fold changes were calculated as log2[(percentage of the indicated subpopulation in old ear wounds)/(percentage of the indicated subpopulation in young ear wounds)]. **j**, Pathway and gene set overdispersion analysis (PAGODA) of the single cell RNA-seq data described in **i**. PAGODA was performed using all KEGG pathways, the “*In vitro* fibroblast aging”, the “Fibroblast activation” signatures (see Supplementary Table 2b, f). Hierarchical clustering is based on 58 significantly overdispersed gene sets (see Extended Data Fig. 9c for full list of significant KEGG pathways). Upper heatmap depicts single cells from young and old wounds and cell clusters identified by Seurat and PAGODA analyses. Middle heatmap show separation of cells based on their PC scores for a subset of the top significantly overdispersed gene sets. The PC scores are oriented so that high (maroon) and low (blue) values generally correspond, respectively, to increased and decreased expression of the associated gene sets. The lower heatmap shows the expression of a subset of secreted factor genes. Expression is depicted as VST-normalized read counts, and scaled row-wise. See Extended Data Fig. 9d-e for the full list of genes. The lower panel depicts the log2 fold changes in the number of cells in each of the six subpopulations identified by PAGODA, between young and old wounds at day 7. Fold changes were calculated as in i.

Among the factors secreted by activated fibroblasts, we tested if cytokines influence reprogramming efficiency (Fig. 3h). Purified IL6 enhanced reprogramming efficiency by ∼2 fold (as was previously reported)^39,61^, whereas TNFα, and IL 1β reduced reprogramming efficiency by ∼2 fold in both young and old fibroblasts (Fig. 4b, Extended Data Fig. 8a-d). Consistently, blocking IL6 with an antibody reduced reprogramming efficiency (Extended Data Fig. 8e-h), whereas blocking TNFα enhanced it (Extended Data Fig. 8e-h). To determine if IL6 and TNFα may contribute to the individual-to-individual variability in reprogramming efficiency, we compared pairs of good and bad old fibroblast cultures in the context of their own conditioned media. While blocking IL6 had a minor impact, blocking TNFα reduced the difference in reprogramming efficiency between pairs of good and bad old cultures by 44% (Fig. 4c). Hence, TNFα (and possibly its cumulative effect with opposing cytokines such as IL6) could contribute to the individual-to-individual variability in reprogramming efficiency with age. TNFα may negatively impact reprogramming efficiency by inducing NF−κB phosphorylation (Fig. 4d, Extended Data Fig. 8a) and expression of *Dot1l* (Extended Data Fig. 8j), which encodes an H3K79 methyltransferase known to impede reprogramming^35,62^. Indeed, the TNF signature in old fibroblasts co-occurs with the NF−κB signature (Extended Data Fig. 8k). Thus, inflammatory cytokines, including TNFα, could influence variability of reprogramming efficiency by activating a transcriptional/chromatin network that hampers the plasticity of cells in response to reprogramming factors.

**Extended Data Figure 8.**
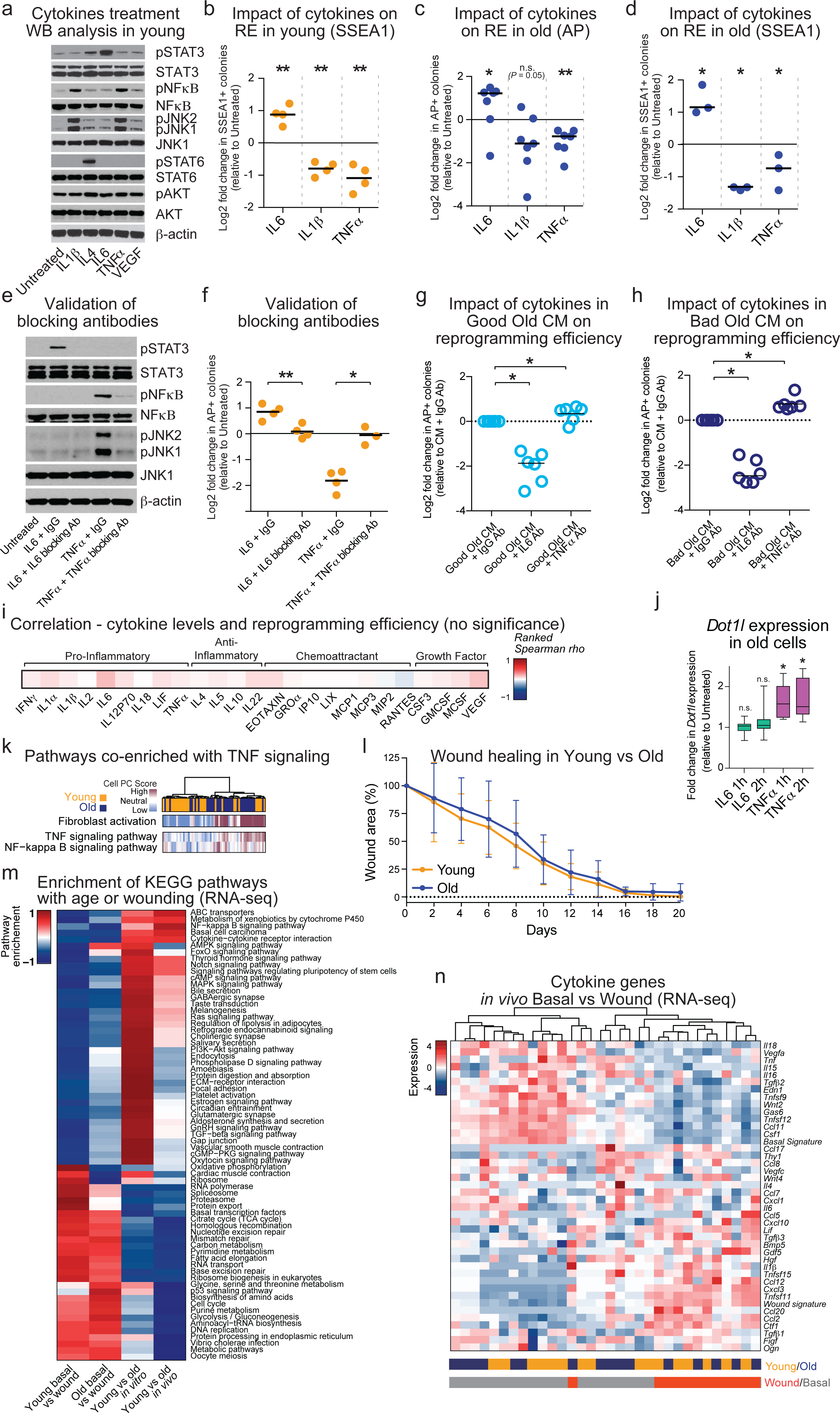
Old fibroblasts secrete cytokines, including IL6 and TNFα, that induce inflammatory signaling pathways and modulate reprogramming efficiency. **a**, Western blot analysis using the indicated antibodies of young fibroblasts at passage 3 treated with the indicated cytokines at the concentration of 10 ng/mL for 30 min. Representative of 3 independent experiments. **b**, Reprogramming efficiency (RE), assessed by stage-specific embryonic antigen 1 (SSEA1) staining, of young fibroblasts at passage 3 treated with the indicated cytokines from day one post-infection at the concentration of 10 ng/mL. Results are shown as log2 fold change in the RE over the RE of untreated cells. Data shown are from a total of 4 young mice from 2 cohorts, combined from 2 independent experiments (for individual experiments, see Supplementary Table 7). Each dot represents cells from an individual mouse. Lines depict median log2 fold change. *P*-values, when calculated using a one-tailed Wilcoxon signed rank test and adjusted for multiple hypothesis testing using Benjamini-Hochberg correction, were not significant (*P* = 0.06) because of low sample number (4). In this case, a t-test was performed and adjusted for multiple hypothesis testing using a Benjamini-Hochberg correction. ** *P*< 0.01. **c**, RE, assessed by alkaline phosphatase (AP) staining, of old fibroblasts at passage 3 treated with the indicated cytokines from day one post-infection at the concentration of 10 ng/mL. Results are shown as log2 fold change in the RE over the RE of untreated cells. Data shown are from a total of 7 old mice from 2 cohorts, combined from 3 independent experiments (for individual experiments, see Supplementary Table 7). Note that 1 old culture was used in 2 independent experiments. In this case, an average of the resultant measurements was determined so as to not inflate the statistical power. Each dot represents cells from an individual mouse. Lines depict median log2 fold change. To test if IL6 increased and IL1β and TNFα decreased RE also in old fibroblasts, *P*-values were calculated using a two-tailed Wilcoxon signed rank test and adjusted for multiple hypothesis testing using a Benjamini-Hochberg correction. * *P* < 0.05, ** *P* < 0.01. n.s. = not significant. **d**, RE, assessed as in b, of old fibroblasts at passage 3, treated with the indicated cytokines from day one post-infection at the concentration of 10 ng/mL. Results are shown as log2 fold change in the RE over the RE of untreated cells. Data shown are from a total of 3 old mice from 2 cohorts, combined from 2 independent experiments (for individual experiments, see Supplementary Table 7). Each dot represents cells from an individual mouse. Lines depict median log2 fold change. *P*-values, when calculated using a one-tailed Wilcoxon signed rank test and adjusted for multiple hypothesis testing using a Benjamini-Hochberg correction, were not significant (*P* = 0.08) because of low sample number (3). In this case, a t-test was performed and adjusted for multiple hypothesis testing using a Benjamini-Hochberg correction. * *P*< 0.05. **e**, Western blot analysis using the indicated antibodies of young fibroblasts at passage 3 treated with the indicated cytokines (10ng/mL) and blocking antibodies (Ab) (8μg/mL) for 30 min. Cytokines were pre-treated with either IgG or their corresponding blocking Ab for one hour prior to treatment. Representative of 2 independent experiments. **f**, RE, assessed as in c, of young cells treated with the indicated conditions from day one post-infection. Cytokines (10ng/mL) were pre-treated with either IgG or their corresponding blocking Ab (8µg/mL) for one hour prior to treatment. Results are shown as log2 fold change in RE over RE of untreated cells. Data shown are from a total of 4 young mice from 2 cohorts, combined from 2 independent experiments (for individual experiments, see Supplementary Table 7). Each dot represents cells from an individual mouse. Lines depict median log2 fold change. *P*-values, when calculated using a one-tailed Wilcoxon signed rank test and adjusted for multiple hypothesis testing using Benjamini-Hochberg correction, were not significant (P = 0.06) because of low sample number (4). In this case, a t-test with adjustment for multiple hypothesis testing using Benjamini-Hochberg correction was used. * *P* < 0.05, ** *P*< 0.01. **g-h**, RE, assessed as in c, of good old (g) or bad old (h) fibroblasts at passage 3, treated with the indicated conditions from day one post-infection. Conditioned media was pre-treated for one hour with the indicated blocking antibody before administration. Results are shown as log2 fold change relative to conditioned media + IgG. Data shown are from a total of 6 young mice (recipient cells) from 2 cohorts, combined from 3 independent experiments (for individual experiments, see Supplementary Table 7). Each dot represents cells from an individual mouse. Lines depict median log2 fold change. To test if blocking of IL6 decreased and blocking of TNFα increased RE, *P*-values were calculated using a one-tailed Wilcoxon signed rank test and were adjusted for multiple hypothesis testing using Benjamini-Hochberg correction. * *P* < 0.05, CM = conditioned media. **i**, Heatmap showing the Spearman rank correlation coefficients between the levels of individual cytokines and reprogramming efficiency in young and old cells. Data shown are from a total of 19 young and 18 old mice from 3 cohorts, combined from 2 independent experiments. *P*-values were computed using algorithm AS 89 and were adjusted for multiple hypothesis testing using Benjamini-Hochberg correction. None of the cytokines detected exhibited significant correlation with reprogramming efficiency. The lack of direct correlation between levels of individual cytokines and reprogramming efficiency could reflect the insensitivity of the multiplex fluorescent immunoassay, the cumulative effect of cytokines that act in opposing manner on reprogramming efficiency, and/or the fact that that cytokines can directly impact the levels of other cytokines^103,104^. **j**, Real-time quantitative PCR (RT-qPCR) of *Dot1l* expression of old fibroblasts at passage 3, treated with the indicated cytokines at the concentration of 10 ng/mL for the indicated time-point. Results are presented as fold change in expression of *Dot1l* compared to untreated cells, using a box-and-whisker plot to indicate the median and interquartile range with whiskers indicating minimum and maximum values. *Hprt1* was used as housekeeping gene. Data shown are from a total of 6 old mice from 2 cohorts of mice combined over 3 independent experiments (for individual experiments, see Supplementary Table 7). *P*-values were calculated using a two-tailed Wilcoxon signed rank test and were adjusted for multiple hypothesis testing using Benjamini-Hochberg correction. * *P*< 0.05. n.s. = not significant. **k**, Pathway and gene set overdispersion analysis (PAGODA) of single cell RNA-seq data from young and old fibroblasts performed using all KEGG pathways, the “*in vitro* fibroblast aging”, the “fibroblast activation”, and *de novo* gene sets after accounting for cell cycle phases. The dendrogram shows hierarchical clustering of 84 significantly overdispersed gene sets and the 248 genes driving the significantly overdispersed gene sets. The heatmaps show separation of the cells by the cell PC score (blue: low, maroon: high) for TNF and NF_κ_B signaling pathways. The full list of significant KEGG pathways is included in Extended Data Fig. 5b-c. **l**, Ear wound healing curve from young and old mice. Full thickness wounds were induced on the dorsal side of both ears (see Experimental Procedures for further details), and the size of the wounds was assessed by imaging ear wounds every second day for 20 days. For each mouse, the average of both ear wounds was calculated. Graph depicts the average % of wound area remaining at the indicated time-points. Data shown are from a total of 26 young and 28 old mice from 2 cohorts of mice combined over 2 independent experiments (for individual experiments, see Extended Data Fig. 7). Lines represent mean and standard deviation. **m**, Comparison between the transcriptomic changes that occur in fibroblasts with age *in vitro*, and *in vivo*, as well as changes that occur in upon wounding in young and old ears. The heatmap depicts the enrichment of the KEGG pathways that are present in at least two of the conditions described. For the complete list of significant KEGG terms, see Supplementary Table 2e, 4g, 5b. The scale for enrichment is indicated on the left. **n**, Heatmap of expression of a subset of cytokine genes from population RNA-seq of fibroblasts from young and old ears during basal and wounded conditions. Expression is depicted as log2 VST normalized read counts, and scaled row-wise. The scale for expression fold changes is indicated on the left. Basal and wound signatures refer to the average expression of the genes that go significantly down or up with wounding, respectively, in this dataset.

We next tested the *in vivo* relevance of these findings. Fibroblasts are critical *in vivo* for wound healing, a process associated with trans/dedifferentiation events of several cell types^63-65^. Fibroblasts participate in wound healing through remodeling of the extracellular matrix and secretion of inflammatory cytokines^4-6^. While the influence of aging on wound healing has been examined^4,66,67^, the variability of this response, and how it affects fibroblast heterogeneity, is not known. We first determined how old age impacts wound healing capacity in different individuals. Notably, there was an increased variability in ear wound healing capacity between old individual mice (Fig. 4e-f, Extended Data Fig. 8l) (median wound healing efficiency was not significantly affected by age). The proportion of THY1+ activated fibroblasts increased after wound healing in both young and old mice (Fig. 4g). Population RNA-seq on basal and wounded THY1+ cells from young and old mice showed that while wounded fibroblasts differed from basal ones (whether young or old) (Extended Data Fig. 8m-n, Supplementary Table 5a-b), cytokine-related signatures (e.g. cytokine-cytokine receptor interaction) were also significantly different between young and old wounded fibroblasts *in vivo* (Fig. 4h, Supplementary Table 5c-d).

Single-cell RNA-seq on freshly isolated fibroblasts pooled from 10 young or 10 old mice at seven days post-induction of wounds revealed age-associated changes in fibroblast heterogeneity (Fig. 4i-j). Analysis of 3036 high quality single cell transcriptomes identified 2 main clusters of cells in tSNE: one enriched for young cells and the other for old cells (Fig. 4i, Extended Data Fig. 9a-b). Unbiased PAGODA clustering confirmed these two cell populations, and could further divide them into six subpopulations (Fig. 4j, Extended Data Fig. 9c). Interestingly, two of the subpopulations that increased in proportions in old tissues were enriched for TNF signaling and cytokine expression (Fig. 4j, Extended Data Fig. 9c-e, g). Hence, fibroblasts are heterogeneous *in vivo* in old tissues, and the presence of fibroblasts with high TNF and cytokine signaling might contribute to the variability in wound healing between old individuals.

**Extended Data Figure 9.**
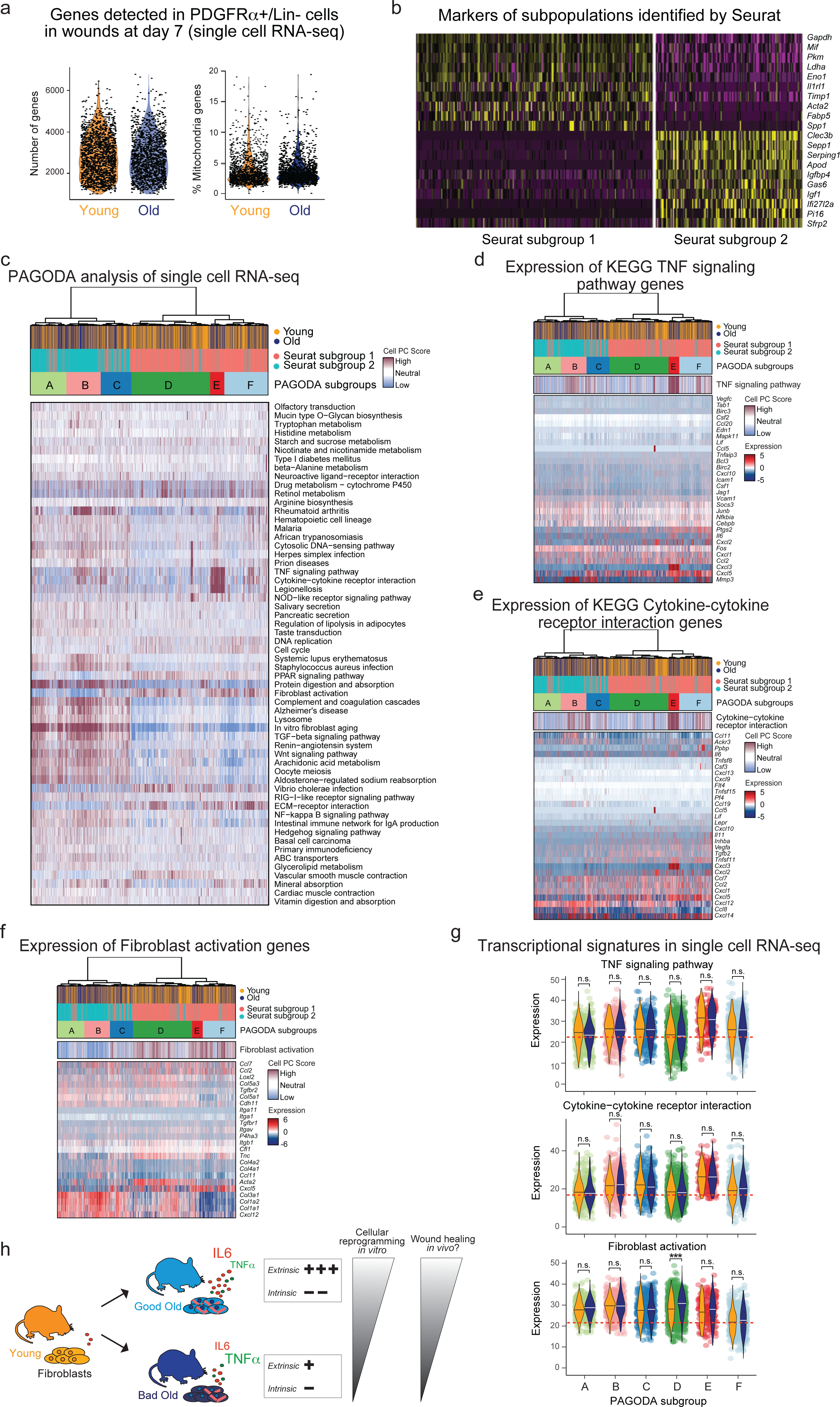
Single cell RNA-sequencing of PDGFRα+/Lin-cells in young and old wounds at day 7 identifies multiple subpopulations that change in proportion with age and that are enriched for different signaling pathways. **a**, Quality control for 10x Genomics single cell RNA-sequencing data of freshly isolated PDGFRα+/Lin-(CD45-/CD31-/EpCAM-/TER119-/TIE2-) cells from young (n=10) and old (n=10) ear wounds, 7 days post-induction of wounds. Unique gene counts (left panel) and percentage of mitochondrial genes (right panel) for each cell are shown, separated by age group. **b**, Seurat analysis of all live high quality PDGFRα+/Lin-cells (3036 cells in total) identified two main clusters of cells. Heatmap depicts the expression of the top 10 marker genes for each significant cell cluster identified by Seurat, which are defined as the genes that are most specific to each population. The cell subpopulation identity assigned to each cluster is indicated below each column. **c**, Pathway and gene set overdispersion analysis (PAGODA) of single cell RNA-seq data described in a. PAGODA was performed using all KEGG pathways, the “*In vitro* fibroblast aging”, the “Fibroblast activation” signatures (see Supplementary Table 2b, f). Hierarchical clustering is based on 58 significantly overdispersed gene sets and the 315 genes driving the significantly overdispersed gene sets. Upper heatmap depicts single cells from young and old wounds and cell clusters identified by Seurat and PAGODA analyses. Lower heatmap show separation of cells based on their PC scores for the significantly overdispersed gene sets. The PC scores are oriented so that high (maroon) and low (blue) values generally correspond, respectively, to increased and decreased expression of the associated gene sets. **d**, PAGODA as described in c. Middle heatmap shows separation of cells based on their PC scores for the KEGG TNF signaling pathway gene set. Lower heatmap shows expression of the genes that are part of the KEGG TNF signaling pathway, where expression is depicted as VST normalized read counts, and scaled row-wise. The scale for expression fold changes is indicated on the right. **e**, PAGODA as described in c. Middle heatmap shows separation of cells based on their PC scores for the KEGG Cytokine-cytokine receptor interaction gene set. The lower heatmap shows expression of the genes that are part of the KEGG Cytokine-cytokine receptor interaction, where expression is depicted as VST normalized read counts, and scaled row-wise. The scale for expression fold changes is indicated on the right. **f**, PAGODA as described in c. Middle heatmap shows separation of cells based on their PC scores for the “Fibroblast activation” signature. The lower heatmap shows expression of the genes that are part of the “Fibroblast activation” signature (see Supplementary Table 2f), where expression is depicted as VST normalized read counts, and scaled row-wise. The scale for expression fold changes is indicated on the right. **g**, Combined log-normalized expression values of genes in TNF signaling (upper panel), Cytokine-cytokine receptor interaction signatures (middle panel), and Fibroblast activation (lower panel) for specific cell populations. Age-associated differences in gene expression signatures were calculated using a two-tailed Wilcoxon rank sum test and were adjusted for multiple hypothesis testing using Benjamini-Hochberg correction. *** *P* < 0.001. n.s. = not significant. **h**, Schematic illustration of the main findings of our study: Aging is associated with an increased individual-to-individual variability in cellular reprogramming *in vitro* and in wound healing *in vivo*. *In vitro*, this increased variability could be a cumulative effect of a shift in the old fibroblast population to a more activated state that is intrinsically less plastic but that promotes reprogramming extrinsically through secretion of inflammatory cytokines. Particularly, the secretion of IL6 and TNFα can impact cellular reprogramming and contribute to the variability with age. *In vivo*, there is a similar age-associated shift in fibroblast populations and secretory phenotype, which could contribute to the variability in wound healing between old individuals.

Collectively, our data show that aging is associated with an increased individual-to-individual variability in cellular reprogramming *in vitro* and in wound healing *in vivo*. *In vitro,* this increased variability could be the cumulative effect of a shift in the old fibroblast population to a more activated state that is intrinsically less plastic but that promotes reprogramming extrinsically through secretion of inflammatory cytokines with opposing effects on the reprogramming process, such as TNFα and IL6. *In vivo*, the age-associated shift in fibroblast populations and secretory phenotype could contribute to the variability in wound healing (Extended Data Fig. 9h). As fibroblasts exhibit tissue-specific properties^68^, there could also be increased variability in different tissues with age. Increase in variability between individuals or cells is emerging as common feature of aging^18-21,45-49^. Our study shows for the first time that a key driver of inter-individual variability is the inflammatory cytokine TNFα (though other intrinsic and extrinsic factors may also exist), and this inflammatory cytokine may also regulate the variability of other aging phenotypes. In addition to genetic and environmental factors, inter-individual variability could reflect distinct aging trajectories between different individuals, and it illustrates the need for personalized treatments for age-related diseases.

Cellular reprogramming is critical as a potential rejuvenation strategy and for regenerative medicine^8-17^. As reprogramming efficiency is a key challenge in the generation of iPSCs from old donors, our findings raise the possibility that targeting inflammatory cytokines could help unify iPSCs generation from elderly patients, especially in old donors with poor reprogramming potential. Furthermore, cell composition and inflammatory cytokines could play an important role in determining the rejuvenation potential of an old tissue.

A subpopulation of activated fibroblasts that secrete cytokines could be a source of chronic inflammation in old individuals, contributing to the recruitment of immune cells to the tissue, and perhaps to cell senescence, which plays a key role in tissue aging^25,26,69^. Activated fibroblasts (which proliferate) differ from senescent fibroblasts (which are permanently arrested), and they appear to secrete overlapping, yet distinct sets of cytokines^24,70,71^. These different types of fibroblasts may contribute in different manners to physiological wound healing (e.g. wound closure and maturation)^4-6^, but also to pathologies such as fibrosis and chronic inflammatory diseases (e.g. rheumatoid arthritis and scleroderma)^4-6,51,72^. Wound healing, both for acute and chronic wounds, is a major issue for elderly individuals. Thus, the changes in fibroblast subpopulations during aging, and the cytokines involved, could provide individualized ways to improve wound healing and reduce fibrosis in old age.

## Author Contributions

S.M., E.M. and A.B. planned the study, with input from M.P.S. and J.W.. S.M. and E.M. performed the reprogramming experiments, analyzed and interpreted data. S.M., E.M., L.X. wrote the manuscript with the help of A.B.. S.M. generated, processed and analyzed the RNA-seq data sets, analyzed the metabolomics data, and performed most THY1-related and conditioned media experiments. E.M. generated and propagated transgene-free iPSC lines. All other studies were done by both E.M and S.M., unless otherwise noted. A.M. and S.M. performed wound healing experiments under the supervision of M.T.L.. L.X. helped with reprogramming, FACS, and IF experiments. F.K.J. generated the *in vitro* single cell RNA-seq data under the supervision of M.P.S.. R.S. generated the ChIP-seq libraries under the supervision of J.W.. K.H. helped with statistics and PAGODA analysis. X.L. performed metabolomics experiments and helped with metabolomics data analysis and validation under the supervision of M.P.S.. K.D. helped with reprogramming and western-blotting experiments. L.P. helped with reprogramming and RT-qPCR experiments. C.E.A. and Y.S. performed the iN reprogramming experiment under the supervision of M.W.. B.A.B. helped with analysis of the epigenomic data. A.L.S.C. identified and collected the human samples. All authors discussed the results and commented on the manuscript.

## Acknowledgments

We thank D. Wagh from Stanford Functional Genomics Facility for help with 10x Genomics single cell RNA-seq, C. Carswell-Crumpton and M. Weglarz from Stanford Shared FACS Facility for FACS support. We thank L. Liu for help with shRNA experiments. The piggyback OSKM transposon constructs were a kind gift from Dr. Keisuke Kaji. We thank V. Sebastiano and M. Kareta for help with iPSC generation and quality assessment. We thank V. Sebastiano, K. Andreasson, C. Weyand, and T. Wyss-Coray for helpful discussions and input on the manuscript. We thank L. Booth, T. Ruetz, P. Singh, X. Zhao, P. Navarro, B. Dulken and other members of the Brunet lab for helpful discussions and feedback on the manuscript. We thank S. Chen, A. Freund, J. Goudeaux, M. Pesch, A. Roux and J. Reuter for feedback on the initial manuscript. Work supported by NIH P01 GM099130 (A.B., M.P.S., J.W.), CIRM RB4-06087 (A.B.), a generous philanthropic gift from Michele and Timothy Barakett, the EMBO post-doctoral fellowship (S.M.), the WennerGren post-doctoral fellowship (S.M.), the Sweden-America Foundation post-doctoral fellowship (S.M.), the Dean’s Fellowship (E.M.), K99 AG049934 (B.A.B.), the Stanford Graduate Fellowship (L.X.), the NSF Graduate Research Fellowship (L.X.), R00 AG049934 and the Hanson-Thorell family fellowship (B.A.B.), the Hagey Laboratory for Pediatric Regenerative Medicine, The Gunn/Olivier fund, the Johnson Longaker fund and R01 GM116892 (M.T.L.).

## Accession Numbers

All raw sequencing reads for population RNA-seq, ChIP-seq and single cell RNA-seq data can be found under BioProject PRJNA316110.

## Experimental Procedures

### Animals

All animals were treated and housed in accordance to the Guide for Care and Use of Laboratory Animals. All experimental procedures were approved by Stanford’s Administrative Panel on Laboratory Animal Care (APLAC) and were in accordance with institutional and national guidelines. Male C57BL/6 mice at different ages (3-4 and 20-29 months) were obtained from the National Institute on Aging (NIA) colony, and were acclimated to the animal facility at Stanford University for at least 1 week before being processed. No live animals were censored.

### Collection of blood and plasma from young and old mice

To assess for systemic changes associated with age, whole blood was collected from young and old mice by cardiac puncture into a tube containing EDTA (Thermo Fischer Scientific, #AM9262) (for a final concentration of 5mM EDTA per blood sample). Blood cell composition, including white and red blood cell, granulocyte, monocyte, lymphocyte, and platelet counts were analyzed with a Hemavet Multispecies Hematology Analyzer (CDC Technologies) according to manufacturer’s instructions. Plasma was prepared from whole blood samples by two consecutive centrifugation steps at 500 rcf and 13,000 rcf, respectively, each for 10 min at room temperature, and then aliquoted and stored at −80°C for cytokine profiling (see below).

### Generation of primary cultures of fibroblasts from young and old mice

To investigate the impact of aging on tissue fibroblasts, primary fibroblast cultures were established from the ears and lungs of young and old mice as previously described^73^. To this end, the ears and lungs were cut into small fragments (∼1 mm^2^), and digested in Dulbecco’s Modified Eagle Media (DMEM, Invitrogen, #11965-092) supplemented with 0.14 Wunsch units/mL of Liberase TM (Roche, #05401127001) for 30-90 min. The fragments were washed with DMEM supplemented with 15% fetal bovine serum (FBS, Gibco, #16000-044, Lot# 551495, 1551824), and plated on tissue culture plates with DMEM supplemented with 15% FBS and 1% Penicillin/Streptomycin/Glutamine (PSQ) (Gibco, #10378). To isolate primary adult fibroblasts from the chest area^74^, the skin on the chest was dissected from the animals, the subcutaneous fat and fascia were removed, and the tissues were incubated overnight at 4°C with the epidermal layer of the skin facing down on top of a solution of 0.25% trypsin (Gibco, #25200-056). The following day, the epidermis was removed, tissues were cut into small fragments (∼1 mm^2^), and treated with 1000 U/mL Collagenase I (Gibco, #17100017) in DMEM for 60-90 min at 37°C. Digested fragments were funneled through a 70 µM nylon mesh (Fischer scientific #08-771-2), washed with fibroblast growth media (DMEM supplemented with 10% FBS and 1% PSQ) and plated using the same media. The cells were passaged once, before being aliquoted, frozen, and stored in liquid N_2_ (passage 1.5, P1.5). For all experiments, unless stated, fibroblasts were thawed (passage 2, P2) and cultured at 37°C in 5% CO_2_and 95% humidity in fibroblast growth media. All experiments, unless specifically noted, were performed at passage 3 (P3).

### Fluorescence-activated cell sorting and analysis of primary fibroblasts

To determine the purity of the primary fibroblasts from young and old mice, FACS analysis was performed on fibroblast cultures at passage 3. FACS analysis was performed using an LSR II flow cytometer (BD Biosciences), and analyzed using FlowJo version 10.0.7. For FACS analysis, fibroblasts were stained with phycoerythrin-conjugated CD140a (BioLegend, #APA5), in combination with the following allophycocyanin conjugated antibodies: B220 (eBioscience, #RA3-6B2), CD3 (BD Pharmingen, #SP34-2), Gr-1 (eBioscience, #RB6-8C5), F4/80 (eBioscience, #BM8), Siglec H (BioLegend, #551), CD11c (eBioscience, #N418) and propidium iodide staining solution (BD Pharmingen).

### Generation of primary cultures of fibroblasts from young and old human individuals

To determine if primary fibroblasts from humans also exhibit an inflammatory profile, we collected biopsies from humans at different ages. Stanford Human Subjects approval and informed consent was obtained prior to all study procedures (under Protocol ID#25269, IRB#350). Biopsies were collected from male participants of different ages with four biological grandparents of Ashkenazi Jewish descent, generally healthy without thyroid disease, diabetes, immunodeficiency, ongoing cancer or autoimmune disease, and no history of poor wound healing (Supplementary Table 1g). A 4 mm punch biopsy of pre-auricular skin was obtained after injection of 1% lidocaine with epinephrine (1:1,000,000). Skin biopsies were rinsed with PBS, cut into smaller fragments (∼1 mm^2^), and plated into a dry 6-well tissue culture plate. Excess PBS was removed, and fibroblast growth media (DMEM supplemented with 10% FBS and 1% PSQ) was added. Tissue was incubated at 37°C in 5% CO_2_ and 95% humidity. After 24 hours, tissue was supplemented with fibroblast growth media, and media changes were repeated every 3–4 days. The cells were passaged once, before being aliquoted, frozen, and stored in liquid N_2_ (passage 1.5, P1.5).

### Cytokine profiling analysis on plasma and conditioned media using Luminex multi-analyte

We examined the impact of aging on the inflammatory profiles by performing cytokine profiling on plasma and conditioned media from fibroblast and iPSC cultures from young and old mice. Plasma was collected as described above. Conditioned media from young and old mouse (ear and lung) and human (skin) fibroblasts was collected 48 hours after plating from 150,000 – 200,000 primary fibroblasts (passage 3 or 33) plated in a 6 cm dish with 2 mL of fibroblast growth media. Conditioned media from iPSCs (passage 23) was collected 24 hours after plating from 500 000 cells maintained in serum-and feeder-free culture condition in 2i media (see below for more information). Conditioned media from THY1+ and THY1− FACS-sorted young and old fibroblasts cultures (passage 4-6, see below for FACS sorting protocol) was collected 24 hours after plating from 500,000-1 million cells plated in a 15cm dish with 20 mL of media. Conditioned media was collected, centrifuged at 10,000 rcf for 10 min at room temperature, aliquoted, and stored at −80°C. For all of these conditions, cell number was determined for each plate by counting on hemocytometer for normalization purposes. In addition, cell-free media was used to assess background. All cytokine profiling was performed by the Stanford Human Immune Monitoring Center using a Luminex mouse 38-plex or a human 62-plex analyte platform (eBiosciences/Affymetrix) that detects 38 or 62 secreted proteins, respectively.

All plasma samples were measured in technical duplicates, and all conditioned media were measured in single technical replicates per recommendation of the Human Immune Monitoring Center at Stanford University. All our analyses were performed using mean fluorescent intensity (MFI) values because converting MFI to clinically-relevant measures (e.g. pg/mL) can introduce a degree of error [Luminex Corporation. xPonent logistic curve fitting: technical notes. Available at: http://www.luminexcorp.com/prod/groups/public/documents/lmnxcorp/207-xponent3.1-log-curve-fit.pdf. Accessed January 10, 2012]. We also report pg/mL conversions in Supplementary Table 1a-f to facilitate comparison with existing literature. To compare values across plates and independent experiments, the MFI values were normalized to the median of young (3 months) within each experiment, generating fold change values. In addition, the conditioned media levels were normalized to respective cell number. Two plasma samples from old mice were discarded, as the coefficient of variation (CV) was > 20% for most of the cytokines measured between the two technical replica for these two plasma samples. Ranked fold changes in cytokine levels were calculated by multiplying the log2fold median change (old/young) with the −log10(pvalue). Similarly, ranked spearman rho correlations were calculated by multiplying the spearman rho values with −log10(pvalue).

### Lentiviral production for reprogramming

To induce reprogramming in fibroblasts and generate iPSCs, we used the lentiviral vector 4F STEMCAA-LoxP, containing a floxed version of EF1α-STEMCCA allowing expression of human *OCT3/4*, *KLF4*, *SOX2* and *c-MYC*^37^. Lentiviruses were produced in Human Embryonic Kidney 293T (293T, ATCC, #CRL-11268) packaging cells as previously described^75^. Briefly, one day prior to transfection, 9 × 10^6^ 293T cells were seeded in a 10-cm dish in 293T media (DMEM supplemented with 10% FBS, 1% PSQ). The next day, the cells were transfected as follows: 100μL of 1 mg/mL polyethylenimine (PEI, Polysciences, #23966-2, linear 25kDa) was added to 2 mL of DMEM and incubated for 10 min at room temperature. The lentiviral vector of interest (20 µg) was mixed with lentiviral packaging vectors (1 µg of pHDM-tat1b (PlasmID), 1 µg of pRC-CMV-rev1b (PlasmID), 1 µg of pHDM-Hgpm2 (PlasmID)) and envelope vector (2 µg of HDM-VSV-G (PlasmID), then added to the PEI/DMEM mixture, and incubated for 15 min at room temperature. The PEI/DMEM/DNA mixture was then added drop wise to the 293T cells, and 12 hours post transfection the media was replaced by 8 mL fresh 293T media. Viral supernatant was collected at 24 and 36 hours post-transfection, centrifuged at 3000 rpm for 15 min, and carefully transferred into a fresh tube, and 0.7 ml of the crude virus supernatant was used to reprogram primary fibroblasts (see below).

### Reprogramming of young and old fibroblasts to iPSCs and iPSC characterization

We generated iPSC lines from 3 independent young fibroblast cultures and from 3 independent old fibroblast cultures (Supplementary Table 1f). Reprogramming of primary fibroblasts was induced as follows: 100,000 primary fibroblasts at passage 3 were seeded in a well of a 6-well plate, and were infected 24 and 36 hours post-seeding with 0.7 mL crude virus supernatant mixed with 8 µg/mL of Polybrene (Sigma, #H9268-5G). Forty-eight hours post-seeding (12 hours after the last round of infection), the infected primary fibroblasts were seeded at a density of 5,000 cells on a 10-cm dish containing 1.5 × 10^6^ γ-irradiated feeder cells (mouse embryonic fibroblasts, MEFs). Cells were maintained in fibroblast growth media for 7 days, and then switched to mouse embryonic stem cell (mESC) media, consisting of DMEM, GlutaMax (Life Sciences, #10569-010), 15% FBS, 1% PSQ, 5 × 10^5^ units of Leukemia Inhibitory Factor (LIF, EMD Millipore, #ESG10007), 1% MEM non-essential amino acids (Gibco, #11140-050) and 0.0008% β-mercaptoethanol (Sigma, #M-7522). Day 13-15 post-infection, colonies with distinct mESC morphology were manually picked from 10-cm dishes and each iPSC clone was transferred into a well of a 96-well plate (primary plate) in the presence of γ-irradiated MEFs. A minimum of 24 iPSC clones per parental fibroblast line were picked, and replicas of each 96-well primary plate were created. These replica plates were used to evaluate the number of viral integrations in each clone, while primary plates were temporarily frozen and stored at −80°C. To determine the number of viral integrations, an on-plate genomic DNA extraction was performed as previously described^76^, and the Mouse TaqMan^®^ Copy Number Reference Assays from Thermo Fisher was used to estimate the number of viral integrations from the genomic DNA extracted. A Taqman probe targeting the Human *KLF4* gene (FAM^™^ dye labeled) was used because the 4F STEMCAA-LoxP vector contains the human version of the reprogramming factors (Life Technologies, #4331182). A Taqman probe targeting the mouse Transferrin receptor gene (*Trfc*), which is known to be encoded by a single gene in the mouse genome, was used as the reference assay (VIC^®^ dye labeled) (Life Technologies, #4458366). Only iPSC clones with an estimated viral integration number equal to or lower than 3 were chosen for further analysis.

For 13 of these lines, we generated transgene-free iPSC lines by excising the reprogramming factor construct and performed long-term passaging (until passage 23), as this is known to improve the pluripotency state^13,32,77^. To this end, primary plates were quickly thawed and the iPSC clones were transferred into a fresh 96-well plate in the presence of γ-irradiated MEFs, and subsequently expanded. At passage 10, the integrated 4F STEMCCA lentiviral construct was excised using Cre-recombinase expressed under the CAG promoter (pCAG-Cre)^37^. The pCAG-Cre construct was transfected using a Mouse ES Cell Nucleofector Kit (LONZA, #V4XP-3012) according to manufacturer’s instructions. Transfected cells were then resuspended in mESC media, seeded on feeders at a very low density in a 10-cm dish (500 cells/dish), and cultured in mESC media until colonies appeared. For each iPSC clone, multiple subclones were isolated and expanded. The efficiency of Cre-recombinase excision was assessed by PCR using the Mouse TaqMan^®^ Copy Number Reference Assays as described above. Only transgene-free iPSC clones were further characterized. iPSC lines were maintained on ESC media for 10 passages post-excision, before being adapted to serum-and feeder-free culture condition in 2i media according to CReM Boston University ESC Culture Protocols (http://www.bu.edu/dbin/stemcells/protocols.php). All molecular characterizations of the iPSC lines were performed at passage 23.

To assess whether the derived iPSC lines could give rise to cell types from all three germ layers upon embryoid body formation (Extended Data Fig. 2b), embryoid body (EBs) formation was induced as previously described^78^. Briefly, iPSCs at passage 23 were incubated with Accutase (EMD Millipore) for 5 min at 37°C to obtain a single cell suspension, and 10 mL of iPSC suspension at a density of 10^3^ cells/mL was seeded on ultra-low attachment plates (Corning). Cells were allowed to form EBs. After 4 days, EBs were transferred into regular tissue culture grade plates in DMEM high glucose supplemented with 10% FBS, 100 U/mL penicillin, and 100 µg/mL streptomycin (Gibco) and were allowed to differentiate. At day 14 after EB differentiation, differentiated cells were collected and analyzed by qRT-PCR for the expression of endodermal, mesodermal, and ectodermal markers (see primers in Supplementary Table 6).

The inflammatory, transcriptomic, and metabolomic profiling of the iPSC lines was performed at passage 23.

### RT-qPCR on iPSCs and EB differentiated cells

To assess expression of specific genes in iPSCs and in embryoid body differentiated cells, RNA purification and cDNA synthesis was performed as previously described^79^. Briefly, total RNA was isolated using the RNeasy RNA Purification Kit (QIAGEN) and 0.5-1 µg of RNA was reverse-transcribed using the High Capacity cDNA Reverse Transcription Kit (Applied Biosystems) according to the manufacturer’s instructions. cDNA was used for RT-qPCR on the BioRad iCycler using iQ SYBR Green Mix (BioRad). All primer sequences are listed in Supplementary Table 6a.

### Assessment of reprogramming efficiency

To determine the impact of aging on iPSC generation, reprogramming efficiency (RE) was quantified by the 96-well assay as previously described^80^. Briefly, reprogramming was induced as described above. Forty-eight hours post-seeding (12 hours after the last round of lentiviral infection), the infected primary fibroblasts were seeded at a density of 20-40 cells per well into 96-well plates containing 1,000 γ-irradiated feeder MEFs per well. In experiments using cytokines and conditioned media, 0.1% gelatin-coated plates (Tribec Science, #TBS8004) without feeders were used to avoid confounds from the feeder cells.

Infected primary fibroblasts were maintained on fibroblast growth media until day 7 post-seeding and then switched to ESC media until day 13-15. In experiments assessing the impact of conditioned media on reprogramming efficiency, fresh conditioned media was collected from 10-25 cm dishes that were growing in parallel, centrifuged 10,000 rcf for 10 min at RT, and added every day, starting from day 1 of re-plating into 96-well plates until the end of experiment. In experiments testing the influence of specific cytokines on reprogramming efficiency, fresh media with the indicated cytokine was added every day until the switch to iPSC media.

To assess reprogramming efficiency, staining with alkaline phosphatase (AP) (an early marker of pluripotency^43^) was performed by fixing the cells in 4% paraformaldehyde (Santa Cruz, #sc-281692) for 15 min at room temperature, washing with citrate solution (Sigma Aldrich #3861) then staining with prepared diazonium salt solution (Sigma Aldrich #851) with Napthol (Sigma Aldrich #855) overnight. Quantification was performed by counting the number of wells containing at least one AP positive colony. To complement AP staining, we also used staining with stage-specific embryonic antigen 1 (SSEA1) (a later marker of pluripotency^43^). SSEA1 staining was performed using StainAlive mouse anti-mouse antibody (Stemgent, #09-0067) according to the manufacturer’s recommendations. Quantification was performed by counting the number of wells containing at least one SSEA1 positive colony using a Zeiss inverted microscope (Zeiss AxioVision A10).

Reprogramming efficiency (RE) was calculated as the number of AP-or SSEA1-positive clones, divided by the number of cells plated, and multiplied by the efficiency of viral infection (see below, Immunofluorescence of pluripotency markers). To compare RE across plates (and independent experiments), the RE of all individual cultures were normalized to the median RE of young cultures within a given experiment. Statistical differences in variance in reprogramming efficiency between the age groups were calculated using the non-parametric Fligner-Killeen test using R v.3.3.0. To assess whether the increased variability in RE with age was introduced by pooling multiple cohorts, we performed a permutation test where the null distribution was estimated by randomly assigning the age groups to the observed RE for individual cultures within each cohort, and the mean difference in standard deviation between young and old cells was calculated across the cohorts. This was repeated 1000 times, and the *P* value was calculated as the percentage of differences greater than or equal to the actual observed difference in standard deviation. This approach indicated that the increased variability in RE with age is not simply due to the pooling of multiple cohorts (*P* < 0.001).

### Immunofluorescence staining for reprogramming factors and pluripotency markers

For immunofluorescence (IF) staining of pluripotency markers, cells were fixed in 4% paraformaldehyde for 15 min at room temperature, then permeabilized with 0.5% Triton X-100 for 10 min, blocked in blocking solution (2% bovine serum albumin (BSA), 5% glycerol, 0.2% Tween-20, 0.1% sodium azide in PBS) for 1h, followed by incubation with primary antibodies. The following antibodies where used for IF: rabbit anti-OCT3/4 (Santa Cruz Biotech, #sc9081), SSEA1 StainAlive rabbit anti-mouse antibody (Stemgent, #09-0067), rabbit anti-SOX2 (Santa Cruz Biotech, #sc17320). The nuclei were stained with DAPI (Life Technologies). Cells were imaged using a Zeiss inverted microscope (Zeiss AxioVision A10) with AxioVision 4.7.2 software. For infection efficiency calculations, 5-10 images were randomly taken per sample and uploaded on ImageJ (v1.46r), and infection efficiency was calculated by dividing the number of OCT3/4 positive cells with the total number of cells (DAPI stain).

### Reprogramming of young and old fibroblasts to induced neurons (iNs)

To determine the ability of young and old primary fibroblasts to reprogram to induced neurons (iNs), iN reprogramming was induced as previously described^81^. Briefly, young and old fibroblast cultures (passage 3) were plated at a density of 60,000 cells per well of a 12-well plate. The following day, the fibroblasts were infected as described above with lentiviruses carrying TetO-FUW-ASCL1 (Addgene #27150), TetO-FUW-BRN2 (Addgene #27151), TetO-FUWMYT1L (Addgene #27152) and FUW-rtTA (Addgene #20342). The next day, doxycycline (2 µg/mL, Sigma) in fibroblast growth media was added to the wells. Media was changed to neuronal media [N2 + B27 + DMEM/F12 (Invitrogen) + 1.6 mL Insulin (6.25 mg/ml, Sigma)] + doxycycline (2 µg/mL) two days after the first doxycycline induction. Subsequently, neuronal media was changed every three days. To determine the number of iNs at day 7 for each fibroblast cultures, the cells were digested using 0.25% trypsin (Invitrogen) at 37°C for 5 min, and all cells were subjected to Magnetic activated cell sorting (MACS) to select for APC-conjugated PSANCAM positive cells (Miltenyi, #130-093-273), according to the manufacturer’s instructions. The number of PSA-NCAM-positive cells for each fibroblast culture was counted manually using a hemacytometer. The reprogramming efficiency for each line was obtained by dividing the total number of PSA-NCAM positive cells obtained at day 7 to the number of fibroblasts plated. The ability of the primary fibroblast cultures to undergo iN and iPSC reprogramming was assessed in parallel. Note that in this comparison, infection efficiency was not assessed and hence not included in the calculation of reprogramming efficiency.

### RNA-seq analysis

To profile transcriptomic changes in primary fibroblast cultures with age and upon iPSC reprogramming, total RNA was isolated from passage 3 fibroblasts and passage 23 iPSCs using RNeasy kit (QIAGEN) according to the manufacturer’s instructions. Total RNA (150 ng) was used to prepare RNA-seq libraries using the Encore Complete RNAseq library kit (Nugen Technology, #0333), according to the manufacturer’s instructions. Libraries were sequenced on HiSeq 2000 (2×101bp paired-end reads, Illumina).

Quality and adapter trimming of the fastq files was performed using TrimGalore version 0.2.8, maintaining reads with a minimum Phred score of 15. The trimmed reads were mapped to the mouse genome (mm9 build) using TopHat (v2.0.8b). Reads per genes were counted using HTSeq (v0.6.1). As annotation file, we used the genes.gtf downloaded from UCSC on March 6^th^, 2013. Gene expression was analyzed using DESeq2 (v1.20.0). For differential expression analysis of fibroblasts, batch-effect was accounted for by including a batch variable into the DESeq2 model (see Supplementary Table 2a). Genes with > 0.3 fpkm in at least one sample within a particular analysis, were considered expressed and included in the analysis. Heatmaps, hierarchical clustering and principal component analysis (PCA) were performed on log2 VST values (variance stabilizing transformation, implemented in DESeq2). Genes were considered significantly differentially expressed if they had FDR < 0.05 and an absolute fold change > 1.5, unless stated otherwise. Publicly available data sets were downloaded from the GEO database (Supplementary Table 6b), and processed as described above. Note these following RNA-seq samples were excluded from further analyses; (1) 2 old and 3 middle-aged RNA-seq libraries as they lacked any young samples, and hence batch-effects could not be corrected for. (2) RNA-seq libraries from 1 good old and 1 bad old fibroblast cultures as their reprogramming efficiency could not be confirmed over several independent experiments. (3) RNA-seq libraries from two iPSC lines (out of 13 total) failed at the QC stage because they exhibited large differences (for example in number of reads mapped) from the rest of the samples (Supplementary Table 2a).

Pathway enrichment analysis was performed using one-sided Fisher’s exact test, testing for the overrepresentation of significantly differentially expressed genes in a given gene list. As background, all the genes that were considered expressed (see above) were used. *P*-values adjusted for multiple hypothesis testing using Benjamini-Hochberg correction, and FDR = 0.05 was set as upper threshold. In Extended Data Fig. 3m, analysis of gene set enrichment was conducted using the Gene Set Enrichment Analysis (GSEA) (v2.2.2) tool. For this analysis, the log2 VST values (derived from DESeq2) was used, and enrichment statistics were calculated using the ‘classic’ method parameter. Nominal *P*-values were calculated based on 10,000 permutations. In Fig. 2i, 4h and Extended Data Fig. 5a, 6l, 8m, analysis of gene set enrichment was conducted by calculating the arithmetic mean of gene-wise test statistics (Wald test statistic from differential expression analysis using DESeq2) per gene set. To calculate a *P*-value for each gene set, we constructed a null distribution of test statistics by sampling 10,000 times n genes (n: number of genes in the respective gene set) and calculating the mean of the test statistics for these genes. A gene set-wise *P*-value was then calculated as the percentage of absolute (sampled) mean test statistics that are equal or greater than the absolute (observed) mean test statistic for that pathway. *P*-values were corrected for multiple hypothesis testing using the Benjamini-Hochberg algorithm using FDR = 0.05 as threshold. This method for gene set enrichment analysis has been shown to outperform many commonly used methods^82^. Kyoto Encyclopedia of Genes and Genomes (KEGG), Gene ontology (GO) terms were acquired from http://amp.pharm.mssm.edu/Enrichr/#stats.

Upstream regulator analysis was performed using QIAGEN’s Ingenuity Pathway analysis (IPA QIAGEN Redwood City) software, using the genes that passed the filter in our dataset as reference genome.

Motif analysis of promoter regions (TSS: +50 −1000 bp) of differentially expressed genes was performed using the Homer software (v4.8)^83^, using the genes that passed the filter in our dataset as background.

### ChIP-seq and analysis of the epigenomic landscape

To profile changes in the epigenomic landscape in primary fibroblast cultures with age, we performed ChIP experiments using H3K4me3 (Active Motif, #39159) and H3K27me3 (Active Motif, #39536) antibodies. Briefly, 1-2 × 10^6^ fibroblasts were crosslinked with 1% formaldehyde for 10%min at room temperature, and formaldehyde was quenched by addition of glycine to a final concentration of 0.125%M. Chromatin was sonicated to an average size of 0.5–2%kb, using Bioruptor (Diagenode). A total of 5%µg of antibody was added to the sonicated chromatin and incubated overnight at 4°C on a rotating platform. 10% of chromatin used for each ChIP reaction was retained as input DNA. Subsequently, 100%µL of protein G Dynal magnetic beads were added to the ChIP reactions and incubated for four additional hours at 4°C. Magnetic beads were washed, followed by reversal of crosslinks and DNA purification. Resultant ChIP DNA was dissolved in water. ChIP and input libraries were generated according to Illumina protocol and sequenced single-end 50bp using the Illumina HiSeq 2000 platform.

For analysis, Fastq reads were quality-trimmed using the trim-galore software (v0.2.1), with a Phred score threshold of 15, and a minimum remaining read length of 36bp. Trimmed reads were mapped to the mm9 genome assembly using bowtie v0.12.7^84^. Duplicate reads were eliminated using the FIXSEQ software with default parameters^85^. ChIP-seq peaks were called in all samples using the MACS (v2.08) software with default settings and the “--broad” option^86^. Input datasets were used as baseline.

To identify H3K4me3 and H3K27me3 ChIP-seq peaks with differential intensity in young vs. old or good old vs. bad old samples, we used the “DiffBind” R package (v.1.12.3)^87^. Reads were quantified in each sample over “meta-peaks”, *i.e.* peaks called using pooled reads from one specific mark (respectively H3K4me3 reads and H3K27me3 reads) over all samples. Meta-peaks help best determine peak boundaries^88^. ChIP-seq read counts normalized to Input reads counts by DiffBind were then analyzed using the “DEseq2” package (v1.6.3)^89^ to identify peaks with significantly different intensity. Hierarchical clustering and principal component analysis (PCA) were performed on log2 VST values (variance stabilizing transformation, implemented in DESeq2). The differential peak intensity and pathway analyses were restricted to the peaks that extended at least 100bp into the promoter regions of the nearest genes (defined as transcriptional start site +/− 2000bp), and were performed as mentioned above (see RNA-seq analysis).

Broad H3K4me3 domains are genomic regions coated with H3K4me3, and they are enriched at genes involved in cell identity/function^90-92^. To compare H3K4me3 breadth of samples across young and old samples, we used the approach described in^90^. Briefly, we used the H3K4me3 meta-peaks to compare the signal-to-noise ratio across samples. This revealed that sample number 2 for H3K4me3 from 3-month old fibroblasts was the noisiest sample of the 5 samples. We therefore down-sampled all other 4 samples to match the coverage histogram of that specific sample. We then called peaks as described above using the calibrated files in MACS (v2.08), and isolated the top 5% broadest H3K4me3 domains (broad H3K4me3 domains) from each peak file. We identified reproducible broad H3K4me3 domains by retaining only those that were present in all young or all old samples, respectively.

Bivalent domains are genomic regions coated with both H3K4me3 and H3K27me3^93,94^. To identify differential bivalent regions between young and old samples, we defined robust bivalently marked regions in each age group. For this, we called H3K4me3 and H3K27me3 “meta-peaks” separately at each age. Then, at each age and for each mark, we identified peaks that were supported by all the individual experimental samples (i.e. reproducible peaks). Bivalent peaks were obtained by the intersection of H3K4me3 and H3K27me3 reproducible peaks in all young or old samples.

### Metabolomic analysis

To profile changes in metabolomic features in cultured fibroblasts with age, frozen cell pellets were mixed with 80% methanol (mass-spec grade) in a ratio of 10 µL/mg cell pellet (a million cells weighs roughly 13 mg). The suspension was then processed by three rounds of 1 min vortex at max speed, chilled briefly on ice. The mixture was incubated at 4°C for 20 min before centrifuging at 20,000 × g for 20 min at 4°C. The supernatants were used as metabolite extracts for liquid chromatography-mass spectrometry (LC-MS) analysis. For LC-MS analysis, the metabolite extracts were transferred to 150 µL deactivated glass insert housed in 2-mL brown MS vials (Waters). Chemical standard solution (for quality control) was prepared from synthetic complete mixture from Sigma-Aldrich (Y1501) at a concentration of 19 µg/mL in 80% mass-spec grade methanol (Fischer Scientific). Metabolite extract was analyzed in a platform that consists of Waters UPLC-coupled Exactive Orbitrap Mass Spectrometer (Thermo), using an OPD2 HP-4B column (4.6×50mm) and an OPD2HP-4A guard column (Shodex, Japan). The column temperature was maintained at 45°C. Five µL of each sample maintained at 4°C was loaded by the autosampler in partial loop mode for three times at positive mode and negative mode, respectively. The binary mobile phase solvents were: A, 10 mM NH_4_ OAc in 10:90 Acetonitrile:water; B, 10 mM NH_4_ OAc in 90:10 Acetonitrile:water. Both solvents were modified with 10 mM HOAc for positive mode acquisition, or 10 mM NH_4_ OH for negative mode. The 30-min gradient for both modes was set as: flow rate, 0.1 mL/min; 0-15 min, 99% A, 15-20.5 min, 99% to 1% A; 20.5-25 min, 1% A; 25-25.5 min, 1% to 99% A; 25.5-30 min, 99% A. The MS acquisition was in profile mode and performed with an electrospray ionization probe, operating with capillary temperature at 275°C, sheath gas at 40 units, spray voltage at 3.5 kV for positive mode and 3.1 kV for negative mode, Capillary voltage at 30V, tube lens voltage at 120V, and Skimmer voltage at 20V. The mass scanning used 100,000 mass resolution, high dynamic range for AGC Target, 500 milliseconds as Maximum Inject Time, and 75-1200 m/z as the scan range. The system was operated by Thermo Xcalibur v2.1 software. The raw data files generated from LC-MS were centroided with PAVA program^95^ and converted to mzXML format by an in-house R script (distribution upon request). Mass feature extraction was performed with XCMS v1.30.3^96^. Differential analysis was performed on signal intensity values derived from XCMS using the nonparametric Wilcoxon rank sum test for positive and negative mode separately and adjusted for multiple hypothesis testing using q-value correction using the R package q-value (v2.0.0). The mass features that were found significantly different were manually searched against the Metlin metabolite database (29381867) using 5ppm mass accuracy. Retention time matching with compounds in the standard mixture was also performed for a subset of the metabolite hits. Prior to PCA and hierarchical clustering analysis, signal intensity values derived from XCMS were range-scaled^97^. Pathway analysis was performed using the integrated pathway analysis tool in the MetaboAnalyst 3.0 software^98^, using all the putatively identified metabolites that were found significantly different (FDR < 0.05, absolute fold change > 1.5) together with all the differentially expressed genes from the transcriptomic analysis (see above).

### Single cell RNA-seq analysis in primary cultures of fibroblasts

To assess for cell composition and heterogeneity in primary fibroblast cultures, single cells were isolated from 3 young and 3 old fibroblast cultures at passage 3 (see Supplementary Table 3h). Briefly, 20 single cells per culture were isolated manually by picking isolated cells under a Zeiss inverted microscope (Zeiss AxioVision A10). Single cell RNA-seq libraries were generated using SMARTer Ultra Low Input RNA Kit for Sequencing – v3 (Clontech, #634853), according to the manufacturer’s instructions. Briefly, single cells were directly lysed in 2.5 µL of Clontech reaction buffer and the volume was brought up to 10 µL with sterile water. First strand cDNA synthesis was carried out in 96 well PCR plates as follows: 1 µL of 3’ SMART CDS Primer II A (24 µM) was added and the resulting mix was incubated into a preheated thermocycler at 72°C for 3 min, and then put at 4°C. Next, 7.5 µL of first strand master mix was added (SMARTScribe Reverse Transcriptase, 5X First-Strand Buffer, dNTP Mix and SMARTer IIA Oligonucleotide), mixed, and incubated at 42°C for 90 min and 70°C for 10 min. Finally, the first-strand cDNA was purified with SPRI Ampure XP beads, where 36 µL of the SPRI bead was added into each 20 µL of single-stranded cDNA sample, mixed, and incubated 8 min at room temperature. The samples were placed on a Promega MagnaBot II Magnetic, the supernatant was discarded, and the single-stranded cDNA sample bound to the beads was directly used for double strand cDNA generation. Next, 50 µL of PCR master mix was added to each sample and mixed. Plates were placed in a preheated thermal cycler with a heated lid using the following program: 95°C for 1 min, 18 cycles of 95°C for 30 s, 65°C for 30 s, 68°C for 6 min, followed by 72°C for 10 min and hold on 4°C. Amplified double stranded cDNA was purified using SPRI Ampure Beads (Beckam Coulter), and eluted in 12 µL of purification buffer and kept in −20°C. The quantity and quality of 1 µL of the amplified purified double-stranded cDNA were measured using the Agilent 2100 BioAnalyzer and Agilent’s High Sensitivity DNA Kit (Agilent, #5067-4626). Double-strand cDNA libraries whose bioanalyzer results showed no contamination and a distinct peak at ∼2,000 bp and with ∼2–7 ng of cDNA were selected. This resulted in 8-12 single cell cDNA from each culture. To generate RNA-seq libraries, we next used Nextera XT DNA Library Preparation Kit and Nextera XT Index Kit (Illumina, #FC-131-1096 and #FC-131-1002). Brielfy, 5 µL purified double-stranded cDNA (∼1 ng total) from the previous step was added into each sample well of a 96-well plate, and 10 µL Tagmentation (TD) Buffer was added into each sample and mixed gently. Next, 5 µl amplicon tagment mix was added to the wells and mixed gently. The 96 well plate was sealed and placed in a thermal cycler and incubated at 55°C for 5 min and held at 10°C. The Tn5 transposase was inactivated by adding 5 µL of Neutralization buffer. The tagmented DNA was then amplified using 15 µL of Nextera PCR Master Mix, 5 µL index 1 primers (i7) and 5 µL index 2 primers (i5) into each sample. The final PCR was performed using the following program on a thermal cycler. 72°C for 3 min, 95°C for 30 s, 12 cycles of: 95°C for 10 s, 55°C for 30 s, 72°C or 30 s, and 72° C for 10 min. The PCR products were then purified with Ampure beads. The final libraries were assessed using the Agilent 2100 BioAnalyzer and Agilent’s High Sensitivity DNA Kit. We generated three pooled libraries and sequenced them on 3 lanes of Illumina HiSeq 2000 paired-end 2×101 base-pairs sequencing reads. Quality and adapter trimming of the fastq files was performed using TrimGalore version 0.2.8, maintaining reads with a minimum Phred score of 15. The trimmed reads were mapped to the mouse genome (mm9 build) using TopHat (v2.0.8b). Reads per genes were counted using HTSeq (v0.6.1). As annotation file, we used the genes.gtf downloaded from UCSC on March 6^th^, 2013. On average, 7000 genes were expressed per cell. Genes with at least 10 reads in 3 single cells were considered expressed. Heatmaps, hierarchical clustering and PCA were performed on VST values (variance stabilizing transformation, implemented in DESeq2).

### tSNE and PAGODA analysis of single cell RNA-seq data from cultured cells

To analyze the single cell RNA-seq data, we performed t-distributed stochastic neighbour embedding clustering using the Rtsne R package (v0.14). Single cell RNA-seq data were analyzed using pathway and gene set overdispersion analysis (PAGODA)^55^. PAGODA identifies pathways and sets of genes that are overdispersed in the data and separate the cells based on their expression patterns^55^. We applied PAGODA on the raw counts for all genes that were considered expressed. For gene sets, we used all KEGG pathways as well as an “*In vitro* fibroblast aging” gene set that we defined from comparing the population RNA-seq data from young and old fibroblast cultures (Supplementary Table 2b. Additionally, we used the list of “Fibroblast activation” genes, which are genes that have previously been associated with fibroblast activation (Supplementary Table 2f). We used the PAGODA pipeline with default parameters, unless stated otherwise, and used the SCDE package v0.99.1 in R v3.2.2. PAGODA revealed a relatively strong cell clustering by KEGG cell cycle as well as two *de novo* gene sets (clusters 37 and 119, see Extended Data Fig. 5b) consisting of many cell cycle-related genes. We accounted for this cell cycle aspect of heterogeneity using the pagoda.subtract.aspect() method (see Supplementary Table 3g for the lists of genes in these gene sets). After accounting for cell cycle phases, PAGODA identified 74 KEGG pathways, 8 *de novo* gene sets and the “*In vitro* fibroblast aging” and “fibroblast activation” signatures as significantly overdispersed in the dataset (Extended Data Fig. 5c).

### Immunofluorescence staining for fibroblast activation marker and EdU incorporation

Immunofluorescence (IF) staining was performed as described above (see immunofluorescence of pluripotency markers). The following antibodies where used for IF: mouse anti-αSMA (Abcam, #ab7817). The nuclei were stained with DAPI (Life Technologies). Cells were imaged using a Zeiss inverted microscope (Zeiss AxioVision A10) with AxioVision 4.7.2 software.

EdU incorporation in fibroblast cultures was visualized using the Click-iT EdU Plus Alexa Fluor 594 Imaging Kit (Invitrogen, C10639). Fibroblasts were plated onto glass coverslips (Bellco Glass, #194310012A) in wells of 24-well plates at a density of 20,000 cells per well. After allowing the cells to attach overnight, fibroblasts were incubated in media containing 5-ethynyl-2’-deoxyuridine (EdU, 10 µM) for 4 hours. Cells were then fixed (4% paraformaldehyde in PBS) and permeabilized (0.1% Triton X-100 in PBS). EdU was detected by click reaction according to manufacturer instructions. Cells were incubated in blocking buffer (2% BSA, 5% glycerol, 0.2% Tween-20, 0.1% sodium azide in MilliQ water) and stained with Alexa Fluor 488-conjugated anti-alpha smooth muscle actin antibody (Abcam, #ab184675). Coverslips were mounted onto slides using ProLong Gold with DAPI (Invitrogen, #P36931) and imaged on a Nikon Eclipse Ti/Andor CSU-W1 spinning disk confocal microscope using Andor Zyla and NIS Elements AR software (v4.30.02).

### Senescence in young and old fibroblast cultures

We assessed for senescence-associated β-galactosidase activity (SA-β-gal) in fibroblast cultures using a histochemical staining kit (Sigma-Aldrich, #CS0030) according to the manufacturer’s recommendations. The nuclei were stained with DAPI (Life Technologies). For determining the proportion of senescent cells, 5-10 images were randomly taken per sample and uploaded on ImageJ (v1.46r). Senescence rate was calculated by dividing the number of SA-β-gal positive cells with the total number of cells (DAPI stain).

### Fluorescence-activated cell sorting and analysis of primary fibroblasts

We performed FACS analysis and sorting of THY1+/PDGFRα+and THY1−/PDGFRα+ cells from primary fibroblast cultures at passage 3. FACS analysis was performed on an LSR II flow cytometer (BD Biosciences), and FACS sorting was performed on a BD FACS Aria II sorter, using a 100 µm nozzle at 13.1 PSI. FACS data were analyzed using FlowJo version 10.0.7. Gating was determined using fluorescent-minus-one controls for each color used in each FACS experiment to ensure that positive populations were solely associated with the antibody for that specific marker (see Extended Data Fig. 10). Antibodies conjugates and matched isotype controls were obtained from BioLegend. For FACS analysis of cultured cells, fibroblasts were stained with phycoerythrin-conjugated CD140a (BioLegend, #APA5) and fluorescein isothiocyanate– conjugated CD90.2 (BioLegend, #30-H12).

**Extended Data Figure 10.**
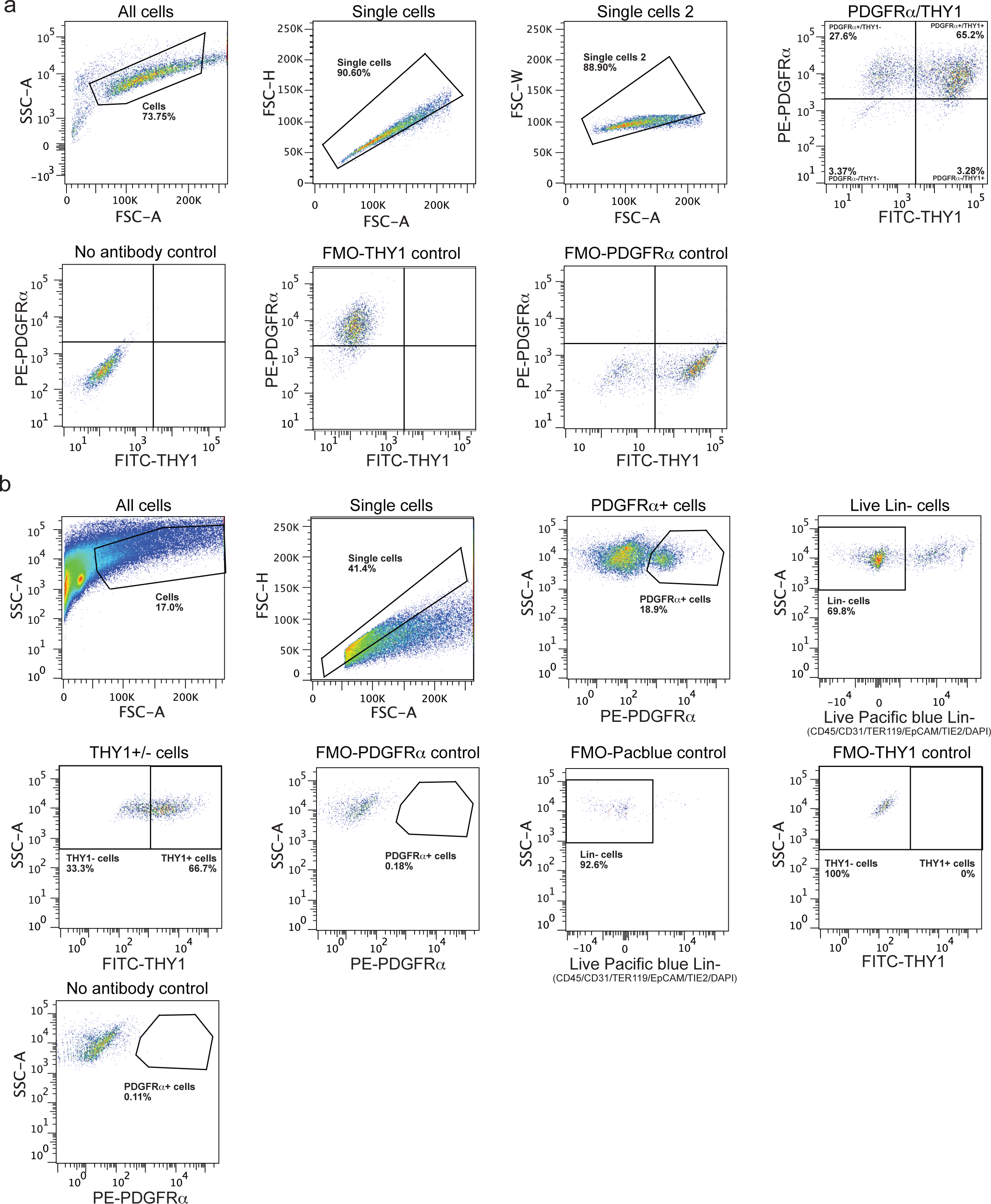
FACS scheme for *in vitro* and *in vivo* fibroblast analysis and sorting. **a**, FACS scheme for analysis and sorting of THY1−/PDGFRα+ and THY1+/PDGFRα+ cells in young and old cultures at passage 3. Gate shown on each plot is indicated above the plot. Marker and fluorophore are shown on each axis. FMO = fluorescence minus one. **b**, FACS scheme for analysis and sorting of live THY1−/PDGFRα+/Lin-and THY1+/PDGFRα+/Lin-cells from young and old fresh tissues (ears), used for population RNA-seq and single cell RNA-seq analyses. Gate shown on each plot is indicated above the plot. Marker and fluorophore are shown on each axis. FMO = fluorescence minus one.

EdU incorporation in fibroblast cultures was assessed by FACS using the Click-iT EdU Plus FACS PacBlue Kit (Invitrogen, #C10636) in accordance with manufacturer’s instructions. Briefly, fibroblasts were incubated in media containing 5-ethynyl-2’-deoxyuridine (EdU; 10 µM) for 4 hours. Cells were then dissociated and resuspended in FACS buffer (1% BSA in PBS). Cell surface markers were stained with phycoerythrin-conjugated CD140a (BioLegend, #APA5) and fluorescein isothiocyanate–conjugated CD90.2 (BioLegend, #30-H12). Cells were then fixed (4% paraformaldehyde, PBS) and permeabilized, followed by click reaction to detect EdU, according to the manufacturer’s instructions.

### RT-qPCR on cultured fibroblasts

To assess expression of fibroblast subpopulation specific genes, young and old fibroblast cultures at passage 3 were FACS-sorted into THY1+ and THY1− fibroblasts (see above for details) and purified fibroblast subpopulations were expanded until passage 5-9. THY1+ and THY1− cells were then plated at a density of 50,000 cells/well in a well of a 6-well plate. After 4 days RNA was isolated from these fibroblast cultures and cDNA synthesis was performed as described above (see RT-qPCR on iPSCs and EB differentiated cells). All primer sequences are listed in Supplementary Table 6a.

### Knock-down of the transcription factor EBF2

To test the functional implication of specific transcriptional regulators, we performed shRNA knock-down experiments. To this end, FACS-purified THY1+/PDGFRα/and THY1−/PDGFRα+ fibroblasts at passage 4-9 were plated at a density of 50,000 cells/well in a well of a 6-well plate. One day after plating, cells were infected by lentiviruses expressing shRNAs. Two independent lentiviral shRNA vectors against *Ebf2* were used (Sigma Aldrich, #TRCN0000081515, # TRCN0000081514). As control, a lentiviral shRNA vector against luciferase was used (Sigma Aldrich, # SHC007V). To produce lentiviruses, we followed the protocol described above (see Lentiviral production for reprogramming). Viral supernatant was collected at 24 hours post-transfection, centrifuged at 3,000 rpm for 15 min, and transferred into a fresh tube. Next, 0.7 ml of the crude virus supernatant was added to THY1+ and THY1− fibroblasts. The media was changed 24 hours post-infection, and the cells were maintained in fibroblast growth media for another 48 hour prior to RNA collection. RNA collection, purification, and RT-qPCR were performed as described above (see RT-qPCR on cultured fibroblasts).

### Proliferation rate of young and old fibroblast cultures

Proliferation rate was assessed by plating young and old fibroblasts at a density of 50,000 cells per well of a 6-well plate in fibroblast growth media. Every second day for up to six days, independent cultures were trypsinized and the number of cells in the cell suspension was counted manually using a hematocytometer. A growth slope was determined as the slope of the regression line based on the data points (cell numbers).

### Fluorescence-activated cell sorting and analysis of fibroblasts *in vivo* in tissues

We isolated fibroblasts from the ears of young and old mice for FACS for quantification and for transcriptomic analysis. Briefly, ears were dissected from animals, cut into small fragments (∼1 mm^2^), and then digested in Dulbecco’s Modified Eagle Media (DMEM, Invitrogen, #11965-092) supplemented with 0.14 Wunsch units/mL of Liberase DL (Roche, # 5401160001) for 30 min at 37°C. The fragments were washed with DMEM supplemented with 20% fetal bovine serum (FBS, Gibco, #16000-044), funneled through a 100 µm nylon mesh (Fischer scientific #08-771-19), and washed with fibroblast growth media (DMEM supplemented with 10% FBS and 1% PSQ). A second filtering was performed using a 40 µm nylon mesh (Fischer scientific #08-771-1), followed by a washing step with fibroblast growth media. Finally, cells were washed with FACS buffer (PBS, 1% BSA, 500nM EDTA), and then re-suspended in FACS buffer to be stained for FACS analysis. FACS analysis and sorting was performed on a BD FACS Aria II sorter, using a 100 µm nozzle at 13.1 PSI. FACS data was analyzed using FlowJo version 10.0.7. Gating was determined using fluorescent-minus-one controls for each color used in each FACS experiment to ensure that positive populations were solely associated with the antibody for that specific marker (see Extended Data Fig. 10). For *in vivo* FACS analysis and sorting the following antibodies were used: CD140a (BioLegend, #APA5), CD90.2 (BioLegend, #30-H12), TER119 (Biolegend, #116234), CD326 (Thermo Fischer Scientific #50-163-76), CD45 (Biolegend, #103126), CD31 (Biolegend, #102422), CD202b (Thermo Fischer Scientific, #15-5987-82), Brilliant Violet 421 Streptavidin (Biolegend, #405226) and DAPI staining solution (Thermo Fischer Scientific, # 62248).

### Bulk RNA-seq of freshly isolated THY1−/PDGFRα+ and THY1+/PDGFRα+ cells from the ear of young and old mice, before and after wounding

To determine if cells could express cytokine *in vivo*, we profiled changes in transcriptomic features in fibroblast subpopulations in tissues from young and old mice, before and after wounding (see below for details on wounding experiments). To this end, RNA-seq was performed on freshly isolated THY1+/PDGFRα+ or THY1−/PDGFRα+ (defined as PDGFRα+/CD45-/CD31-/EpCAM-/TER119-/TIE2-) (see above for isolation). Two to three young or old mice were pooled together to obtain 500 cells of each population for each biological replicate. RNA isolation and generation of RNAseq libraries were performed using the Clontech SmartSeq v4 Ultra-Low Input RNA kit (Clontech). Cells were FACS-sorted directly into lysis buffer, and cDNA was prepared as described by the manufacturer. Each cDNA library was analyzed via bioanalyzer on a High Sensitivity chip on an Agilent 2100 Bioanalyzer. To generate sequencing libraries, 0.15 ng of each cDNA library was used as input in the Nextera XT kit, following manufacturer’s recommendations. Cells were indexed using the Nextera XT Index Kit v2 Set A, and were subsequently multiplexed and sequenced on Illumina NextSeq-500 (400m), using 75bp paired-end reads.

### Assessment of reprogramming efficiency of THY1+ and THY1− fibroblasts

To determine the intrinsic reprogramming efficiency of THY1+ and THY1− fibroblasts, cells were FACS-purified (see above) and plated at a density of 100,000 cells per well of a 6-well plate. Reprogramming was induced as described above (see Reprogramming of young and old fibroblasts to iPSCs and iPSC characterization). In these experiments, 0.1% gelatin-coated plates (Tribec Science, #TBS8004) without feeders were used to avoid confounds from the feeder cells. Fibroblasts infected with lentiviruses expressing the OSKM factors were maintained on fibroblast growth media until day 7 post-reseeding and then switched to ESC media until day 12-13. Reprogramming efficiency was assessed by AP staining as described above (see Assessment of reprogramming efficiency).

### Assessment of reprogramming efficiency of fibroblasts with swapped conditioned media

To assess the contribution of extrinsic factors for reprogramming efficiency, we performed experiments swapping conditioned media. For the conditioned media experiments THY1+/− fibroblasts (passage 5-9), or ‘good’ or ‘bad’ old fibroblasts (passage 3) were plated at a density of 350-400,000 cells per 10 cm plate or at a density of 500,000-1 million cells per 15 cm plates in fibroblast growth media. In parallel, cells were seeded at a density of 100,000 cells per well of a 6-well plate for inducing reprogramming as described above (see Reprogramming of young and old fibroblasts to iPSCs and iPSC characterization). Starting from day one after infection with OSKM, conditioned media was collected from 10 cm or 15 cm dishes from the indicated cultures that were growing in parallel, centrifuged at 10,000 rcf for 10 min at RT, and added every day to the recipient cells replacing their media. At day 7 post-reseeding and onwards, ESC media was made using the conditioned media from fibroblast cultures as a base. Due to the positive effect of conditioned media on cellular reprogramming, reprogramming efficiency was assessed earlier than in other experiments, at day 9-10 post-infection. Reprogramming efficiency was assessed by AP staining as described above (see Assessment of reprogramming efficiency). For experiments using THY1+/− fibroblasts, comparisons were made between pairs of THY1− and THY1+ from the same original culture. For experiments using conditioned media from THY1+/− fibroblasts, conditioned media was collected from the THY1− or THY1+ fibroblasts from the same original culture, and comparisons were made between the effect of the different conditioned media on the specific populations.

### Non-viral reprogramming

To test if variation in reprogramming efficiency could be due to lentiviral infection, we used a non-viral reprogramming protocol. Non-viral reprogramming was induced using the piggyback transposon system containing OSKM^99^. Briefly, FACS purified THY1+ or THY1− cells (passage 4-6) were seeded in a well of a 6-well plate at a density of 100,000 cells per well, and were transfected by the piggyback transposon vector using Lipofectamine 3000 (Life sciences, #11668027), according to manufacturer’s instructions. Transfected fibroblasts were maintained on fibroblast growth media until day 7, and then switched to ESC media until day 16-19. Reprogramming efficiency was calculated by counting the number of AP-positive colonies in the well.

### Effect of cytokines and blocking antibodies on reprogramming efficiency

To test how cytokines impact reprogramming efficiency, the following recombinant cytokines and blocking antibodies were purchased from R&D systems and used according to the manufacturer’s recommendations: recombinant mouse IL6 (R&D systems, #406-ML-025), recombinant mouse IL1β(R&D systems, #401-ML-025), recombinant mouse IL4 (R&D systems, #404-ML-050), recombinant mouse TNFα(R&D systems, #410-NT-050), recombinant mouse VEGF (R&D systems, #493-MV-025), normal polyclonal goat IgG (R&D systems, #AB-108-C), goat polyclonal mouse IL6 blocking antibody (R&D systems, #AB-406-NA), goat polyclonal mouse TNFα blocking antibody (R&D systems, #AB-410-NA). For all experiments, the recombinant cytokines were re-suspended according to manufacturer’s instructions, and then used in culture media at a final concentration of 10 ng/mL. For the blocking antibody experiments, the blocking antibodies were pre-incubated with the corresponding cytokine or conditioned media for 60-90 min prior to treatment. The blocking antibodies (or control IgG) were all used at a concentration of 8 µg/mL. For these experiments, young and old fibroblasts at passage 3 were seeded at a density of 100,000 cells per well of a 6-well plate. To avoid confounds from the feeder cells, cells were plated on 0.1% gelatin-coated plates (Tribec Science, #TBS8004). Reprogramming was induced as described above (see Reprogramming of young and old fibroblasts to iPSCs and iPSC characterization). Starting from day one after infection with OSKM, cells were treated with specific cytokines or with conditioned media together with blocking antibodies. Reprogramming efficiency was calculated by counting the number of AP-or SSEA1-positive colonies in the wells as described above (see Assessment of reprogramming efficiency). For the cytokine experiments, comparisons were made between treated and untreated cells originating from the same infected pool of cells and thus infection efficiency was not taken into account.

### Western blot analyses

To test whether the cytokines used in this study induce their cognate pathways in fibroblasts, we performed western blot analyses. To this end, young and old fibroblasts at passage 3 were seeded at a density of 100,000-150,000 cells in a 6 cm dish in fibroblast growth media. Twenty-four hours post-plating, cells were treated with the indicated cytokines or antibodies for 30 min. Cells were then lysed directly in the culture plates using ice-cold RIPA buffer (50 mM Tris-HCL pH 7.5, 150 mM NaCl, 2 mM EDTA, 1% NP-40, 0.1% SDS supplemented with 1mM aprotinin, PMSF and PhosphoStop (Pierce), scraped and transferred to Eppendorf tubes. Following addition of sample buffer (0.0945 M Tris-HCl [pH 6.8], 9.43% glycerol, 2.36% w/v SDS, and 5% β-mercaptomethanol), samples were resolved on 10% SDS-page gels, transferred onto nitrocellulose membranes, and blotted, according to the following protocol (PMID: 26456332), with the following antibodies: phospho-STAT3 (Tyr705) (Cell Signaling Technology, #9145), STAT3 (Invitrogen, #44-364G), phospho-STAT6 (Tyr641) (Cell Signaling Technology, #9361), STAT6 (Cell signaling Technology, #5397), phospho-AKT (Ser473) (Cell Signaling Technology, #4060), AKT (Cell Signaling Technology, #4691), phospho-NF_κ_B (Ser536) (Cell Signaling Technology, #3033), NF_κ_B (Cell Signaling Technology, #8242), phospho-JNK1&2 (Thr183 & Tyr185) (Invitrogen, #44-682G), JNK1 (Invitrogen, #44-690G), β-actin (Novus Biologicals, #NB600-501). Membranes were incubated with HRP-conjugated anti-mouse or anti-rabbit secondary antibodies (Calbiochem) and visualized using enhanced chemiluminescence detection reagent (Amersham ECL, GE Healthcare).

### Wounding and wound healing experiments

To assess wound healing ability with aging, young (3-4 months) and old (24-26 months) C57BL/6 male mice from the NIA colony were anesthetized in standard fashion by inhalation of 1-4% of isoflourane^100^. The hair on the dorsal aspect of both ears was shaved and cleaned with a 70% ethanol solution. Symmetric full thickness skin wounds were induced on both ears by first gently pressing a 4mm punch biopsy onto the dorsum of the ear at its cartilaginous base. Sharp scissors were then used to dissect away the wheel of skin while leaving the underlying connective tissue, cartilage, and anterior skin. No dressing was applied post-operatively, and the wounds were then allowed to heal without further intervention. Wound healing was assessed by imaging (using a standard iPhone 8S camera) every other day for 20 days. Wound closure was analyzed by comparing the relative wound size at a given time to the original immediate post-operative size, performed as previously described^100^. A wound was considered closed when it became re-epithelialized >95% of its original size^101^. The rate of individual wound healing was determined using the average of the resultant measurements from both ears per individual.

### Single cell RNA-seq from young and old wounds using 10x Genomics Chromium

We assessed cell heterogeneity and cell composition in wounds from young and old mice. To this end, we performed single cell RNA-sequencing of all live PDGFRα+ Lin-cells in the wounded area from young and old mice, 7 days after wounding. We pooled cells from 10 young (3-4 months), or 10 old (25-27 months) male C57BL/6 mouse from the NIA aged colony. FACS sorting was performed as described above. Cells were sorted into fibroblast growth media. Cells were then spun down at 300xg for 5 min at 4^°^C and resuspended in fibroblast growth media at a concentration of 263 cells/µL. Young (18,000) and old (18,000) cells were loaded onto a 10x Genomics Chromium chip per manufacturer’s recommendations. Reverse transcription and library preparation was performed using the 10x Genomics Single Cell v2 kit following the 10x Genomics protocol. One library from young mice and one library from old mice were multiplexed and sequenced on one lane of Illumina NextSeq-500 with a high output (400m) kit.

### Quality control of 10x Genomics single cell RNA-seq

For mapping, sequences obtained from sequencing using the 10x Genomics single cell RNA-sequencing platform were de-multiplexed using the Cell Ranger package from 10x Genomics and mapped to the mm10 transcriptome using the Cell Ranger package (10x Genomics). Cells were removed from subsequent analysis if they were expressing fewer than 500 unique genes or expressed more than 10% mitochondrial reads. Levels of mitochondrial reads and numbers of Unique molecular identifiers (UMIs) were similar between the young and old conditions (Extended Data Fig. 9a), indicating that there were no systematic biases in the libraries from young and old mice. Average gene detection in each cell type was similar between young and old (Extended Data Fig. 9a). Based on these criteria, 3036 high-quality cells were used for downstream analyses.

### tSNE analysis of single cell RNA-seq datasets and identification of cell clusters

To analyze the single cell RNA-seq data, we performed t-distributed stochastic neighbour embedding clustering using the Seurat R Package (version 2.3.4) (tSNE) with the first 30 principal components^102^. Identification of significant clusters was performed using the FindClusters() algorithm in the Seurat package which uses a shared nearest neighbour (SNN) modularity optimization based clustering algorithm^102^. Marker genes for each significant clusters were found using the Seurat function FindAllMarkers(). This analysis identified 2 main clusters of cells.

### PAGODA on single cell RNA-seq data from in vivo cells

We applied PAGODA on the raw counts for all genes that were considered expressed. As for gene sets, we used all KEGG pathways as well as the “*In vitro* fibroblast aging” and “Fibroblast activation” genes sets described above (see Supplementary Table 2b, f). We used the PAGODA pipeline with default parameters, unless stated otherwise, and used the SCDE package v1.99.1 in R v3.3. PAGODA did not reveal strong cell clustering by KEGG cell cycle (see Extended Data Fig. 9c). We therefore did not account for cell cycle in this analysis. PAGODA identified 58 KEGG pathways, “*In vitro* fibroblast aging” and “Fibroblast activation” signatures as significantly overdispersed in the dataset (Extended Data Fig. 9c). Based on these pathways, 2 main clusters of cells were identified, which corresponded to the two clusters identified by Seurat, and 6 subclusters.

### Violin plots for gene expression in single cells

To visualize the expression of individual genes, cells were grouped by their cell type (as determined by PAGODA). Log-transformed and normalized gene expression values as calculated by Seurat were plotted for each cell as a violin plot with an overlying dot plot in R.

### Statistical analysis

For the majority of experiments, young and old mice/sample were processed in an alternate manner rather than in two large groups, to minimize group effect. While we did not do a *bona fide* power analysis, we took into account previous experiments to determine the number of animals needed in each experiment. The exception is the wound healing experiment in Fig. 4, in which a power analysis was performed based on an initial experiment to determine the sample size required to detect a difference in variability with a 95% confidence interval. For all quantifications that were done with FACS or automated image quantification, no blinding was performed, including Fig. 3b, e, 4g. The other experiments were blinded, with the exception of Fig. 1d, e, 4b, Extended Data Fig. 2d, 6e, 8b-d, f. Statistical analysis of the differences between age groups was performed using a two-tailed non-parametric Wilcoxon rank sum test (unpaired) or a two-tailed or a one-tailed non-parametric Wilcoxon signed rank test (paired), unless otherwise stated. The statistical test applied was determined prior to the performing experiments. In cases when the same recipient fibroblast culture was used in 2 independent experiments (Fig. 3g, i, Extended Data Fig. 6a, 7c, 8c), an average of the resultant measurements was determined so as to not inflate the statistical power. The non-parametric Fligner-Killeen test was used to test for differences in variance in reprogramming efficiency. *P*-values were corrected for multiple hypothesis testing using Benjamini-Hochberg correction, unless otherwise stated, and were considered significant when *P* < 0.05.

## Supplementary Table Legends

**Supplementary Table 1: Cytokine profiles of young and old plasma, and of conditioned media from young and old fibroblasts at passage 3.**

**Supplementary Table 2: Analysis of transcriptomic, epigenomic, and metabolomic profiles of young and old fibroblasts at passage 3.**

**Supplementary Table 3: Differential gene expression and pathway enrichment analyses of good and bad reprogramming cultures.**

**Supplementary Table 4: Differential gene expression and pathway enrichment analyses of non-activated and activated fibroblasts *in vitro* and *in vivo*.**

**Supplementary Table 5: Differential gene expression and pathway enrichment analyses of *in vivo* fibroblasts during basal and wound condition.**

**Supplementary Table 6: List of primers used in this study, and information on RNA-seq data used in this study.**

**Supplementary Table 7: Experimental list.**

## References

1. Franceschi, C. et al. Inflamm-aging. An evolutionary perspective on immunosenescence. Ann N Y Acad Sci 908, 244–54 (2000).

2. Franceschi, C., Garagnani, P., Parini, P., Giuliani, C. & Santoro, A. Inflammaging: a new immune-metabolic viewpoint for age-related diseases. Nat Rev Endocrinol 14, 576–590 (2018).

3. Franceschi, C. & Campisi, J. Chronic inflammation (inflammaging) and its potential contribution to age-associated diseases. The journals of gerontology. Series A, Biological sciences and medical sciences 69 Suppl 1, S4–9 (2014).

4. Eming, S.A., Martin, P. & Tomic-Canic, M. Wound repair and regeneration: mechanisms, signaling, and translation. Sci Transl Med 6, 265sr6 (2014).

5. Gurtner, G.C., Werner, S., Barrandon, Y. & Longaker, M.T. Wound repair and regeneration. Nature 453, 314–21 (2008).

6. Lynch, M.D. & Watt, F.M. Fibroblast heterogeneity: implications for human disease. J Clin Invest 128, 26–35 (2018).

7. Staerk, J. et al. Reprogramming of human peripheral blood cells to induced pluripotent stem cells. Cell Stem Cell 7, 20–4 (2010).

8. Lapasset, L. et al. Rejuvenating senescent and centenarian human cells by reprogramming through the pluripotent state. Genes Dev 25, 2248–53 (2011).

9. Mahmoudi, S. & Brunet, A. Aging and reprogramming: a two-way street. Current opinion in cell biology 24, 744–56 (2012).

10. Mertens, J. et al. Directly Reprogrammed Human Neurons Retain Aging-Associated Transcriptomic Signatures and Reveal Age-Related Nucleocytoplasmic Defects. Cell Stem Cell 17, 705–18 (2015).

11. Miller, J.D. et al. Human iPSC-based modeling of late-onset disease via progerin-induced aging. Cell Stem Cell 13, 691–705 (2013).

12. Suhr, S.T. et al. Mitochondrial rejuvenation after induced pluripotency. PLoS One 5, e14095 (2010).

13. Lo Sardo, V. et al. Influence of donor age on induced pluripotent stem cells. Nat Biotechnol 35, 69–74 (2017).

14. Prigione, A. et al. Mitochondrial-associated cell death mechanisms are reset to an embryonic-like state in aged donor-derived iPS cells harboring chromosomal aberrations. PLoS One 6, e27352 (2011).

15. Ocampo, A. et al. In Vivo Amelioration of Age-Associated Hallmarks by Partial Reprogramming. Cell 167, 1719–1733 e12 (2016).

16. Rando, T.A. & Chang, H.Y. Aging, rejuvenation, and epigenetic reprogramming: resetting the aging clock. Cell 148, 46–57 (2012).

17. Ocampo, A., Reddy, P. & Belmonte, J.C.I. Anti-Aging Strategies Based on Cellular Reprogramming. Trends Mol Med 22, 725–738 (2016).

18. Chaleckis, R., Murakami, I., Takada, J., Kondoh, H. & Yanagida, M. Individual variability in human blood metabolites identifies age-related differences. Proc Natl Acad Sci U S A 113, 4252–9 (2016).

19. Ong, M.L. & Holbrook, J.D. Novel region discovery method for Infinium 450K DNA methylation data reveals changes associated with aging in muscle and neuronal pathways. Aging Cell 13, 142–55 (2014).

20. Li, R. et al. Linking Inter-Individual Variability in Functional Brain Connectivity to Cognitive Ability in Elderly Individuals. Front Aging Neurosci 9, 385 (2017).

21. Hultsch, D.F., MacDonald, S.W. & Dixon, R.A. Variability in reaction time performance of younger and older adults. J Gerontol B Psychol Sci Soc Sci 57, P101–15 (2002).

22. Childs, B.G. et al. Senescent cells: an emerging target for diseases of ageing. Nat Rev Drug Discov 16, 718–735 (2017).

23. He, S. & Sharpless, N.E. Senescence in Health and Disease. Cell 169, 1000–1011 (2017).

24. Coppe, J.P. et al. Senescence-associated secretory phenotypes reveal cell-nonautonomous functions of oncogenic RAS and the p53 tumor suppressor. PLoS Biol 6, 2853–68 (2008).

25. Baar, M.P. et al. Targeted Apoptosis of Senescent Cells Restores Tissue Homeostasis in Response to Chemotoxicity and Aging. Cell 169, 132–147 e16 (2017).

26. Baker, D.J. et al. Naturally occurring p16(Ink4a)-positive cells shorten healthy lifespan. Nature 530, 184–9 (2016).

27. Xu, M. et al. Senolytics improve physical function and increase lifespan in old age. Nat Med 24, 1246–1256 (2018).

28. Jeon, O.H. et al. Local clearance of senescent cells attenuates the development of post-traumatic osteoarthritis and creates a pro-regenerative environment. Nat Med 23, 775–781 (2017).

29. Schwartz, S.D. et al. Human embryonic stem cell-derived retinal pigment epithelium in patients with age-related macular degeneration and Stargardt’s macular dystrophy: follow-up of two open-label phase 1/2 studies. Lancet 385, 509–16 (2015).

30. Barker, R.A., Drouin-Ouellet, J. & Parmar, M. Cell-based therapies for Parkinson disease-past insights and future potential. Nat Rev Neurol 11, 492–503 (2015).

31. Cheng, Z. et al. Establishment of induced pluripotent stem cells from aged mice using bone marrow-derived myeloid cells. J Mol Cell Biol 3, 91–8 (2011).

32. Kim, K. et al. Epigenetic memory in induced pluripotent stem cells. Nature 467, 285–90 (2010).

33. Li, H. et al. The Ink4/Arf locus is a barrier for iPS cell reprogramming. Nature 460, 1136–9 (2009).

34. Sharma, A. et al. The role of SIRT6 protein in aging and reprogramming of human induced pluripotent stem cells. J Biol Chem 288, 18439–47 (2013).

35. Soria-Valles, C. et al. NF-kappaB activation impairs somatic cell reprogramming in ageing. Nat Cell Biol 17, 1004–13 (2015).

36. Trokovic, R., Weltner, J., Noisa, P., Raivio, T. & Otonkoski, T. Combined negative effect of donor age and time in culture on the reprogramming efficiency into induced pluripotent stem cells. Stem Cell Res 15, 254–62 (2015).

37. Somers, A. et al. Generation of transgene-free lung disease-specific human induced pluripotent stem cells using a single excisable lentiviral stem cell cassette. Stem Cells 28, 1728–40 (2010).

38. Abad, M. et al. Reprogramming in vivo produces teratomas and iPS cells with totipotency features. Nature 502, 340–5 (2013).

39. Mosteiro, L. et al. Tissue damage and senescence provide critical signals for cellular reprogramming in vivo. Science 354 (2016).

40. Mosteiro, L., Pantoja, C., de Martino, A. & Serrano, M. Senescence promotes in vivo reprogramming through p16(INK)(4a) and IL-6. Aging Cell 17 (2018).

41. Banito, A. et al. Senescence impairs successful reprogramming to pluripotent stem cells. Genes Dev 23, 2134–9 (2009).

42. Marion, R.M. et al. A p53-mediated DNA damage response limits reprogramming to ensure iPS cell genomic integrity. Nature 460, 1149–53 (2009).

43. Brambrink, T. et al. Sequential expression of pluripotency markers during direct reprogramming of mouse somatic cells. Cell Stem Cell 2, 151–9 (2008).

44. Bahar, R. et al. Increased cell-to-cell variation in gene expression in ageing mouse heart. Nature 441, 1011–4 (2006).

45. Hernandez-Segura, A. et al. Unmasking Transcriptional Heterogeneity in Senescent Cells. Curr Biol 27, 2652–2660 e4 (2017).

46. Martinez-Jimenez, C.P. et al. Aging increases cell-to-cell transcriptional variability upon immune stimulation. Science 355, 1433–1436 (2017).

47. Wiley, C.D. et al. Analysis of individual cells identifies cell-to-cell variability following induction of cellular senescence. Aging Cell 16, 1043–1050 (2017).

48. Aguayo-Mazzucato, C. et al. beta Cell Aging Markers Have Heterogeneous Distribution and Are Induced by Insulin Resistance. Cell Metab 25, 898–910 e5 (2017).

49. Busuttil, R., Bahar, R. & Vijg, J. Genome dynamics and transcriptional deregulation in aging. Neuroscience 145, 1341–7 (2007).

50. Enge, M. et al. Single-Cell Analysis of Human Pancreas Reveals Transcriptional Signatures of Aging and Somatic Mutation Patterns. Cell 171, 321–330 e14 (2017).

51. Wynn, T.A. & Ramalingam, T.R. Mechanisms of fibrosis: therapeutic translation for fibrotic disease. Nat Med 18, 1028–40 (2012).

52. Bronte, V. & Zanovello, P. Regulation of immune responses by L-arginine metabolism. Nat Rev Immunol 5, 641–54 (2005).

53. Phang, J.M., Liu, W., Hancock, C. & Christian, K.J. The proline regulatory axis and cancer. Front Oncol 2, 60 (2012).

54. Rath, M., Muller, I., Kropf, P., Closs, E.I. & Munder, M. Metabolism via Arginase or Nitric Oxide Synthase: Two Competing Arginine Pathways in Macrophages. Front Immunol 5, 532 (2014).

55. Fan, J. et al. Characterizing transcriptional heterogeneity through pathway and gene set overdispersion analysis. Nat Methods 13, 241–4 (2016).

56. Koumas, L., Smith, T.J., Feldon, S., Blumberg, N. & Phipps, R.P. Thy-1 expression in human fibroblast subsets defines myofibroblastic or lipofibroblastic phenotypes. Am J Pathol 163, 1291–300 (2003).

57. Katsumata, L.W., Miyajima, A. & Itoh, T. Portal fibroblasts marked by the surface antigen Thy1 contribute to fibrosis in mouse models of cholestatic liver injury. Hepatol Commun 1, 198–214 (2017).

58. Mizoguchi, F. et al. Functionally distinct disease-associated fibroblast subsets in rheumatoid arthritis. Nat Commun 9, 789 (2018).

59. Huynh, P.T. et al. CD90(+) stromal cells are the major source of IL-6, which supports cancer stem-like cells and inflammation in colorectal cancer. Int J Cancer 138, 1971–81 (2016).

60. Driskell, R.R. et al. Distinct fibroblast lineages determine dermal architecture in skin development and repair. Nature 504, 277–81 (2013).

61. Brady, J.J. et al. Early role for IL-6 signalling during generation of induced pluripotent stem cells revealed by heterokaryon RNA-Seq. Nat Cell Biol 15, 1244–52 (2013).

62. Onder, T.T. et al. Chromatin-modifying enzymes as modulators of reprogramming. Nature 483, 598–602 (2012).

63. Donati, G. et al. Wounding induces dedifferentiation of epidermal Gata6(+) cells and acquisition of stem cell properties. Nat Cell Biol 19, 603–613 (2017).

64. Li, J.F. et al. HGF accelerates wound healing by promoting the dedifferentiation of epidermal cells through beta1-integrin/ILK pathway. Biomed Res Int 2013, 470418 (2013).

65. Li, Y., Jalili, R.B. & Ghahary, A. Accelerating skin wound healing by M-CSF through generating SSEA-1 and -3 stem cells in the injured sites. Sci Rep 6, 28979 (2016).

66. Keyes, B.E. et al. Impaired Epidermal to Dendritic T Cell Signaling Slows Wound Repair in Aged Skin. Cell 167, 1323–1338 e14 (2016).

67. Nishiguchi, M.A., Spencer, C.A., Leung, D.H. & Leung, T.H. Aging Suppresses Skin-Derived Circulating SDF1 to Promote Full-Thickness Tissue Regeneration. Cell Rep 24, 3383–3392 e5 (2018).

68. Rinn, J.L., Bondre, C., Gladstone, H.B., Brown, P.O. & Chang, H.Y. Anatomic demarcation by positional variation in fibroblast gene expression programs. PLoS Genet 2, e119 (2006).

69. Baker, D.J. et al. Clearance of p16Ink4a-positive senescent cells delays ageing-associated disorders. Nature 479, 232–6 (2011).

70. Waldera Lupa, D.M. et al. Characterization of Skin Aging-Associated Secreted Proteins (SAASP) Produced by Dermal Fibroblasts Isolated from Intrinsically Aged Human Skin. J Invest Dermatol 135, 1954–1968 (2015).

71. Hernandez-Segura, A., Nehme, J. & Demaria, M. Hallmarks of Cellular Senescence. Trends Cell Biol 28, 436–453 (2018).

72. Lagares, D. et al. Targeted apoptosis of myofibroblasts with the BH3 mimetic ABT-263 reverses established fibrosis. Sci Transl Med 9 (2017).

73. Seluanov, A., Vaidya, A. & Gorbunova, V. Establishing primary adult fibroblast cultures from rodents. J Vis Exp (2010).

74. Collins, C.A., Kretzschmar, K. & Watt, F.M. Reprogramming adult dermis to a neonatal state through epidermal activation of beta-catenin. Development 138, 5189–99 (2011).

75. Mostoslavsky, G. et al. Efficiency of transduction of highly purified murine hematopoietic stem cells by lentiviral and oncoretroviral vectors under conditions of minimal in vitro manipulation. Mol Ther 11, 932–40 (2005).

76. Ramirez-Solis, R. et al. Genomic DNA microextraction: a method to screen numerous samples. Anal Biochem 201, 331–5 (1992).

77. Chin, M.H., Pellegrini, M., Plath, K. & Lowry, W.E. Molecular analyses of human induced pluripotent stem cells and embryonic stem cells. Cell Stem Cell 7, 263–9 (2010).

78. Kurosawa, H. Methods for inducing embryoid body formation: in vitro differentiation system of embryonic stem cells. J Biosci Bioeng 103, 389–98 (2007).

79. Webb, A.E. et al. FOXO3 shares common targets with ASCL1 genome-wide and inhibits ASCL1-dependent neurogenesis. Cell Rep 4, 477–91 (2013).

80. Kareta, M.S. et al. Inhibition of pluripotency networks by the Rb tumor suppressor restricts reprogramming and tumorigenesis. Cell Stem Cell 16, 39–50 (2015).

81. Vierbuchen, T. et al. Direct conversion of fibroblasts to functional neurons by defined factors. Nature 463, 1035–41 (2010).

82. Ackermann, M. & Strimmer, K. A. general modular framework for gene set enrichment analysis. BMC Bioinformatics 10, 47 (2009).

83. Heinz, S. et al. Simple combinations of lineage-determining transcription factors prime cis-regulatory elements required for macrophage and B cell identities. Mol Cell 38, 576–89 (2010).

84. Langmead, B., Trapnell, C., Pop, M. & Salzberg, S.L. Ultrafast and memory-efficient alignment of short DNA sequences to the human genome. Genome Biol 10, R25 (2009).

85. Hashimoto, T.B., Edwards, M.D. & Gifford, D.K. Universal count correction for high-throughput sequencing. PLoS Comput Biol 10, e1003494 (2014).

86. Feng, J., Liu, T., Qin, B., Zhang, Y. & Liu, X.S. Identifying ChIP-seq enrichment using MACS. Nat Protoc 7, 1728–40 (2012).

87. Ross-Innes, C.S. et al. Differential oestrogen receptor binding is associated with clinical outcome in breast cancer. Nature 481, 389–93 (2012).

88. Q Li, J.B., H Huang, PJ Bickel Measuring reproducibility of high-throughput experiments. Annals Applied Statistics 1752–1779 (2011).

89. Love, M.I., Huber, W. & Anders, S. Moderated estimation of fold change and dispersion for RNA-seq data with DESeq2. Genome Biol 15, 550 (2014).

90. Benayoun, B.A. et al. H3K4me3 breadth is linked to cell identity and transcriptional consistency. Cell 158, 673–88 (2014).

91. Chen, K. et al. Broad H3K4me3 is associated with increased transcription elongation and enhancer activity at tumor-suppressor genes. Nat Genet 47, 1149–57 (2015).

92. Bernstein, B.E. et al. Genomic maps and comparative analysis of histone modifications in human and mouse. Cell 120, 169–81 (2005).

93. Azuara, V. et al. Chromatin signatures of pluripotent cell lines. Nat Cell Biol 8, 532–8 (2006).

94. Bernstein, B.E. et al. A bivalent chromatin structure marks key developmental genes in embryonic stem cells. Cell 125, 315–26 (2006).

95. Guan, S., Price, J.C., Prusiner, S.B., Ghaemmaghami, S. & Burlingame, A.L. A. data processing pipeline for mammalian proteome dynamics studies using stable isotope metabolic labeling. Mol Cell Proteomics 10, M111 010728 (2011).

96. Smith, C.A., Want, E.J., O’Maille, G., Abagyan, R. & Siuzdak, G. XCMS: processing mass spectrometry data for metabolite profiling using nonlinear peak alignment, matching, and identification. Anal Chem 78, 779–87 (2006).

97. van den Berg, R.A., Hoefsloot, H.C., Westerhuis, J.A., Smilde, A.K. & van der Werf, M.J. Centering, scaling, and transformations: improving the biological information content of metabolomics data. BMC Genomics 7, 142 (2006).

98. Xia, J., Sinelnikov, I.V., Han, B. & Wishart, D.S. MetaboAnalyst 3.0––making metabolomics more meaningful. Nucleic Acids Res 43, W251–7 (2015).

99. O’Malley, J. et al. High-resolution analysis with novel cell-surface markers identifies routes to iPS cells. Nature 499, 88–91 (2013).

100. Lam, M.T., Nauta, A., Meyer, N.P., Wu, J.C. & Longaker, M.T. Effective delivery of stem cells using an extracellular matrix patch results in increased cell survival and proliferation and reduced scarring in skin wound healing. Tissue Eng Part A 19, 738–47 (2013).

101. Barret, J.P. et al. Accelerated re-epithelialization of partial-thickness skin wounds by a topical betulin gel: Results of a randomized phase III clinical trials program. Burns 43, 1284–1294 (2017).

102. Butler, A., Hoffman, P., Smibert, P., Papalexi, E. & Satija, R. Integrating single-cell transcriptomic data across different conditions, technologies, and species. Nat Biotechnol 36, 411–420 (2018).

103. Confalone, E., D’Alessio, G. & Furia, A. IL-6 Induction by TNFalpha and IL-1beta in an Osteoblast-Like Cell Line. Int J Biomed Sci 6, 135–40 (2010).

104. Turner, N.A. et al. Mechanism of TNFalpha-induced IL-1alpha, IL-1beta and IL-6 expression in human cardiac fibroblasts: effects of statins and thiazolidinediones. Cardiovasc Res 76, 81–90 (2007).

